# An *Xist*-dependent protein assembly mediates *Xist* localization and gene silencing

**DOI:** 10.1101/2020.03.09.979369

**Authors:** Amy Pandya-Jones, Yolanda Markaki, Jacques Serizay, Tsotne Chitiashvilli, Walter Mancia, Andrey Damianov, Costantinos Chronis, Bernadett Papp, Chun-Kan Chen, Robin McKee, Xiao-Jun Wang, Anthony Chau, Heinrich Leonhardt, Sika Zheng, Mitchell Guttman, Douglas L. Black, Kathrin Plath

**Affiliations:** Department of Biological Chemistry, University of California, Los Angeles, Los Angeles, CA 90095, USA; Department of Microbiology, Immunology and Molecular Genetics, University of California, Los Angeles, Los Angeles, CA 90095, USA; Division of Biology and Biological Engineering, California Institute of Technology, Pasadena, CA 91125, USA; Department of Biology and Center for Integrated Protein Science, LMU Munich, 82152 Munich, Germany; Molecular Biology Institute, Jonsson Comprehensive Cancer Center, Brain Research Institute, Graduate Program in the Biosciences, Eli and Edythe Broad Center of Regenerative Medicine and Stem Cell Research, David Geffen School of Medicine at the University of California Los Angeles, Los Angeles, CA 90095, USA

**Author notes:** École Normale Supérieure de Cachan, Université Paris-Saclay, Saclay, France. Current Address: The Gurdon Institute and Department of Genetics University of Cambridge, Cambridge CB2 1QN, United Kingdom. Department of Neurological Surgery, University of California San Francisco, San Francisco, CA 94143, USA. Department of Oral Biology, College of Dentistry, University of Florida, Gainesville FL 32610, USA. Division of Biomedical Science, University of California, Riverside, Riverside, CA 92521, USA.

## Abstract

Nuclear compartments play diverse roles in regulating gene expression, yet the molecular forces and components driving compartment formation are not well understood. Studying how the lncRNA *Xist* establishes the inactive-X-chromosome (Xi)-compartment, we found that the *Xist* RNA-binding-proteins PTBP1, MATR3, TDP43, and CELF1 form a condensate to create an Xi-domain that can be sustained in the absence of *Xist*. The E-repeat-sequence of *Xist* serves a multivalent binding-platform for these proteins. Without the E-repeat, *Xist* initially coats the X-chromosome during XCI onset but subsequently disperses across the nucleus with loss of gene silencing. Recruitment of PTBP1, MATR3, TDP-43 or CELF1 to ΔE-*Xist* rescues these phenotypes, and requires both self-association of MATR3 and TDP-43 and a heterotypic PTBP1-MATR3-interaction. Together, our data reveal that *Xist* sequesters itself within the Xi-territory and perpetuates gene silencing by seeding a protein-condensate. Our findings uncover an unanticipated mechanism for epigenetic memory and elucidate the interplay between RNA and RNA-binding-proteins in creating compartments for gene regulation.

## Main text

The function of long non-coding RNAs (lncRNAs) and the mechanisms by which they act remain largely unknown. One of the best studied lncRNAs is *Xist*, which orchestrates X-chromosome inactivation (XCI) in placental female mammals^1–7^. By spreading across one X-chromosome and mediating chromosome-wide gene silencing, *Xist* equalizes X-linked gene expression with that of males^8–12^. XCI initiates when *Xist* is induced on one of the two X-chromosomes in pluripotent cells of the implanting blastocyst, or upon induction of differentiation in embryonic stem cells (ESCs)^4, 13^, the latter providing a powerful model for the mechanistic dissection of XCI-initiation. Intriguingly, *Xist* shapes nuclear organization during XCI-initiation. *Xist* establishes a transcriptionally silent, intra-chromosomal domain (or compartment) by specifically localizing to the X-chromosome from which it is transcribed and inducing the compaction of the forming inactive X-chromosome (Xi), the enrichment of heterochromatin proteins, the repositioning of silenced genes into the center of the Xi, and the exclusion of active transcriptional regulators, such as RNA polymerase II^1, 2, 14–20^. Yet, the mechanisms that drive and maintain the *Xist* RNA within a spatially confined region to establish this Xi-domain remain unclear.

Mechanistically, *Xist* is thought to act as a modular scaffold for diverse proteins each confering discrete functions on the RNA. These functions include chromatin localization, gene silencing, recruitment of chromatin modifiers, and sub-nuclear-positioning of the Xi^6, 14, 20–27^. Consistent with this model, functional regions within the RNA, structured as six conserved repeat arrays termed A – F, have been identified^1, 6, 28–33^, along with many interacting proteins^22, 25, 34^ and critical RNA-protein interactions^21, 22, 25, 29, 35, 36^. However, which of the *Xist*-interacting proteins play roles in formation of the Xi-domain and the spatial confinement of *Xist* is unclear. Interestingly, during the initiation of XCI, but after initial spreading and gene silencing (within 72h of ESC differentiation), a state is reached where gene silencing remains stable even if *Xist* is experimentally turned off^5^. This transition during XCI initiation from *Xist*-dependent to *Xist-* independent gene silencing^5^ cannot currently be explained. This is partially due to the fact that *Xist*-interacting proteins and functional *Xist* domains have primarily been examined during the *Xist*-dependent phase of XCI initiation.

Among the *Xist*-interacting proteins are RNA-binding proteins (RBPs) with known functions in regulating mRNA processing, including the splicing regulators Polypyrimidine Tract Binding Protein 1 (PTBP1), MATRIN-3 (MATR3), CUG-Binding Protein 1 (CELF1), and TAR-DNA Binding Protein (TDP-43)^22, 25, 34^. X-linked gene silencing is largely unaffected by siRNA-mediated depletion of these factors during the *Xist*-dependent phase of XCI initiation (Extended Data. Fig. 1a-c)^25, 37^, raising the question of what roles these proteins play in XCI. Interestingly, these RBPs have all been shown to form extensive higher-order assemblies, particularly when concentrated by RNAs containing multivalent protein binding sites^38–43, 43^. Because *Xist* contains several highly repetitive sequences that could seed a high spatial concentration of such proteins, we hypothesized that interactions between *Xist* and proteins like PTBP1, MATR3, CELF1, and TDP-43 might create a higher-order assembly within the Xi to establish a nuclear compartment that functions to restrict *Xist* localization and enforce the silent state of the Xi.

Based upon our hypothesis, we first examined whether the depletion of PTBP1, MATR3, CELF1, or TDP-43 has an impact on *Xist* localization. To test this, we knocked down PTBP1, MATR3, CELF1, or TDP-43 at day 3 of female ESC differentiation (during the *Xist*-dependent phase of XCI), and observed nuclear dispersal of *Xist* compared to a control siRNA (Fig. 1a/b). To quantify the *Xist* dispersal, we derived the *Xist* aggregation score, by normalizing the projected *Xist* RNA fluorescent in situ hybridization (FISH) signal by the smallest circular area encompassing the *Xist* FISH mask (Fig. 1c and Extended Data Fig. 1d/e). *Xist* dispersal occurred with small or no changes in *Xist* transcript levels and splicing (Extended Data Fig. 1f-i). Consistent with this finding, PTBP1 knockdown in male ESCs expressing *Xist* from a cDNA transgene, which does not require a splicing event, resulted in a similar dispersal of the *Xist* RNA FISH signal (Extended Data Fig. 2). Depletion of PTBP1, MATR3, CELF1 or TDP-43 also revealed defects in the enrichment of heterochromatin marks on the forming Xi (Extended Data Fig. 1j/k). These findings demonstrate that PTBP1, MATR3, CELF1, and TDP-43 play a role in the enrichment of *Xist* within the X-chromosome territory, which is largely independent of their RNA-processing activities.

**Figure 1:**
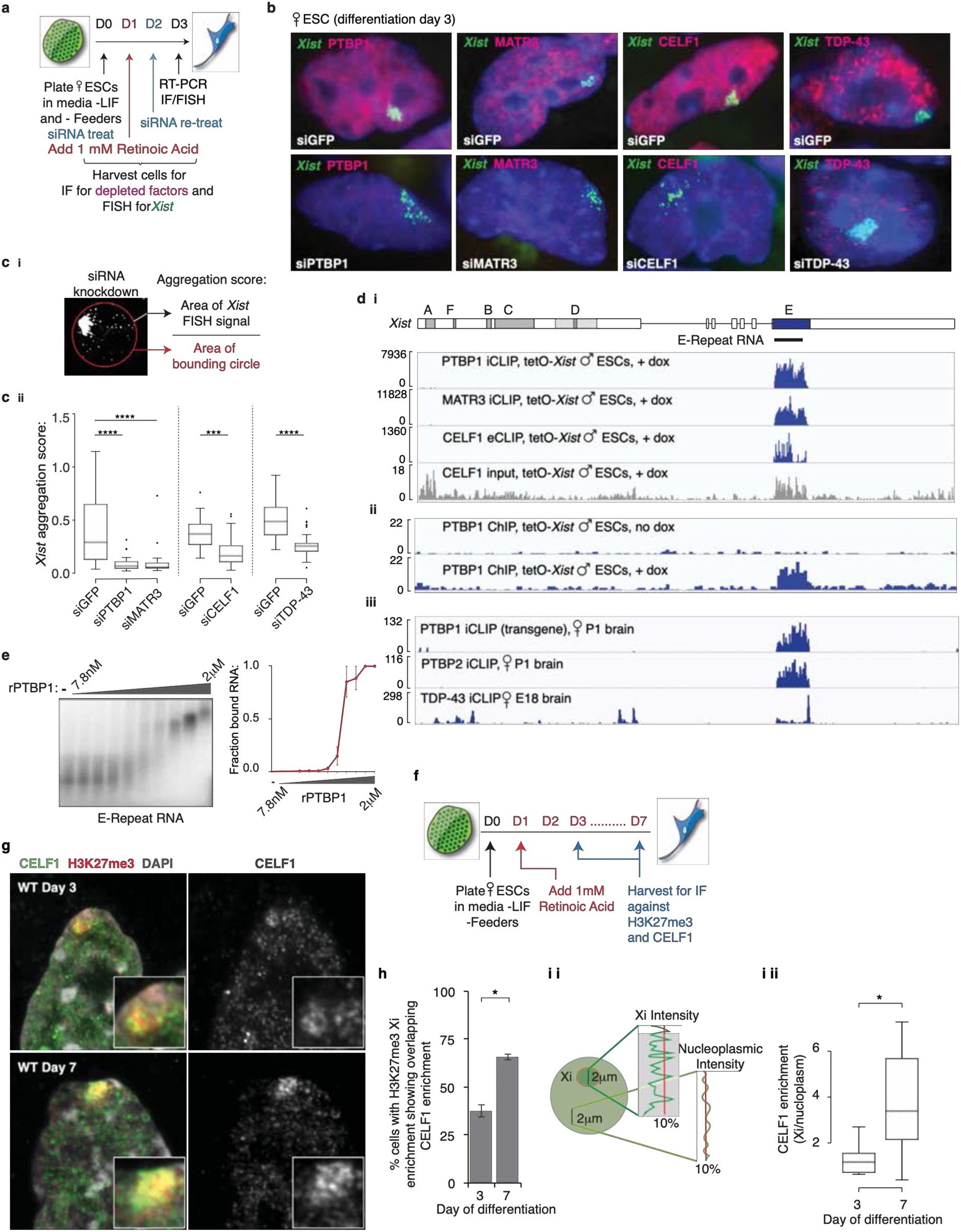
PTBP1, MATR3, TDP-43 and CELF1 bind the *Xist* E-repeat. **a,** Protein knockdown approach in differentiating female ESCs. **b,** Representative images of siRNA-treated differentiating female ESCs immunostained for indicated proteins (red) and probed for *Xist* by RNA FISH (green). **c i,** Depiction of aggregation score calculation. Maximum Intensity Projections (MIPs) of stacked images were used to mask the *Xist* RNA FISH signal and calculate the smallest bounding circle that encompassed the entirety of the *Xist* mask. The aggregation score is the ratio of the *Xist* mask (in pixels) to the bounding circle (in pixels). **ii,** Box plots showing *Xist* aggregation scores upon depletion of indicated proteins. Depletion experiments for CELF1 and TDP-43 were conducted separately to PTBP1 and MATR3, and measured relative to their paired siGFP (control) sample. Significance testing was performed using a 2-sample K-S test (*** p < 0.0005, **** p < 0.00005). No significant differences (p > 0.05) were observed between different siGFP treated samples. 25-36 clouds were measured per sample from one experiment. **d i,** Top: Diagram of the *Xist* locus with introns, exons and repeat arrays A-F of the RNA depicted. The repeat regions (A-D and F) within exon 1 are shown in grey. The sole conserved D-repeat in mouse (dark grey) is flanked by truncated versions (light grey coloring/dotted line). The 1.4kb E-repeat is shown in blue. The *in vitro* transcribed E-repeat RNA used in (e) is indicated. PTBP1 (iCLIP), MATR3 (iCLIP) and CELF1 (eCLIP) CLIP-seq profiles across the *Xist* locus after 6 hours of doxycycline (dox) treatment in male tetO-*Xist* ESCs are shown, together with the CELF1 input sample. **ii,** PTBP1 ChIP-seq profiles across the *Xist* locus before or after 20 hours of dox treatment in male tetO-*Xist* ESCs. **iii,** PTBP1, PTBP2 and TDP-43 iCLIP-seq profiles across the *Xist* locus in female mouse brain. **e,** Left: Radiograph of an EMSA with *in vitro* transcribed E-repeat RNA (see (d)) and either none, or increasing amounts of recombinant (r) PTBP1 (0, 7.8nM, 15.6nM, 31.3nM, 62.5nM, 125nM, 250nM, 500nM, 1μM and 2μM). Right: Quantification showing fraction of bound RNA. Quantification from two independent experiments, with standard error of the mean (SEM) shown. **f,** Depiction of wildtype female ESC differentiation experiment for immunostaining. **g,** Left: Confocal-airyscan optical sections through nuclei of WT female cells at day 3 or 7 of differentiation, immunostained for CELF1 (green) and H3K27me3 (marking the Xi, red) and counterstained with DAPI (gray). Inset: Enlargement of the Xi-territory. Right: Same as left only showing CELF1 staining in greyscale. Note the increased CELF1 staining intensity within the H3K27me3-enriched territory with differentiation time. **h,** Histogram showing the number of H3K27me3-marked Xi’s with a co-localizing CELF1 enrichment on days 3 and 7 of differentiation. 50-90 cells with H3K27me3 enrichment were counted per coverslip. 3 coverslips taken from 2 independent differentiations were counted for each sample. Error bars indicate SEM (p-value: p <0.05, 2-tailed students t-test). **i i,** Intensity values for CELF1 fluorescence were recorded across a 2µm line plot over the Xi (as marked by H3K27me3 staining) or in the nucleoplasm of the same nucleus in z-stack projections, as illustrated in the schematic. **ii,** Box plot showing the distribution of the ratio between the top 10% CELF1 intensity values within the Xi compared to the top 10% intensity values within the nucleoplasm for 12 cells per time point from one experiment.

As these findings indicated a role for these RBPs in controlling *Xist* localization, we next determined where on *Xist* these factors bind, by applying CrossLinking and ImmunoPrecipitation of RNA followed by high-throughput sequencing (CLIP-Seq). CLIP-seq for PTBP1, MATR3 and CELF1 was carried out in male ESCs that can inducibly express endogenous *Xist*, a model system for studying the *Xist*-dependent phase of XCI initiation^24^. Each of the three RBPs yielded a striking enrichment of reads on the E-repeat of *Xist* (Fig. 1d(i) and Extended Data Fig. 3a). Consistent with the binding signature of these proteins, the 1.4kb E-repeat comprises over 50 C/U/G-rich elements predicted to serve as PTBP1, MATR3 and CELF binding sites (Extended Data Fig. 3b-d)^44–46^. The binding of PTBP1 molecules to the E-repeat RNA was further confirmed with recombinant PTBP1 and *in vitro* transcribed E-repeat RNA in electrophoretic mobility shift assays (Fig. 1e). The E-repeat thus serves as a multivalent binding platform for PTBP1, MATR3, and CELF1. As *Xist* was expressed for just 6 hours prior to fixing cells for CLIP, our results indicated that these factors associate with *Xist* at the earliest stage of XCI initiation. To determine whether this interaction occurs co-transcriptionally, we performed Chromatin Immunoprecipitation followed by high-throughput sequencing (ChIP-seq) against PTBP1 and observed binding primarily over the genomic region encoding the E-repeat, upon induction of *Xist* expression, but not before (Fig. 1d(ii)). Taken together, these results demonstrate that PTBP1, and likely MATR3 and CELF1, engage *Xist* as it is transcribed.

To ascertain whether these factors also associate with *Xist* in cells with an established Xi, we examined published datasets. The PTBP1 CLIP-seq profile for *Xist* in differentiated female cells is strikingly similar to that observed for PTBP1 in ESCs, as is the CLIP-seq signature against PTBP2, the neural homolog of PTBP1^47^ (Fig. 1d(iii)). We further found that TDP-43 CLIP-seq in embryonic mouse brain displayed binding at the 3’ end of the E-repeat, where multiple (GU)n tracts presumably serve as TDP-43 binding motifs (Fig. 1d(iii) and Extended Data Fig. 3b/c)^48, 49^. These results indicate that binding of TDP-43 and PTBP1 to the E-repeat persists after XCI initiation is completed and that PTBP2 likely replaces PTBP1 on *Xist* RNA in brain^47^. We conclude that *Xist* is a multivalent binding platform for PTBP1, MATR3, CELF1, TDP-43 and their homologs throughout all stages of XCI.

Next, we applied immunofluorescent staining to differentiating female ESCs to determine whether recruitment of PTBP1, MATR3, CELF1 or TDP-43 by *Xist* could be microscopically oberved within the Xi. These experiments revealed that CELF1 accumulates on the Xi, which was delineated by the enrichment of H3K27me3, and that its accumulation increases in intensity from day 3 to day 7 of differentiation (Fig. 1f-i). We also observed PTBP1 to concentrate within the Xi-territory at later times of differentiation (day 7), in a mesh-like pattern distinct from the punctate staining seen earlier (day 3) (Extended Data Fig. 4a). Although clearly present in the Xi, MATR3 and TDP-43 did not appear enriched in the Xi relative to the nuclear background (Extended Data Fig. 4b/c). Xi-enrichment of these RBPs may not have been detected due to inassessibility of their epitopes or because high concentrations are present elsewhere in the nucleus. These findings indicate that, CELF1 and PTBP1 concentrate over time within the Xi-territory, consistent with the time-dependent formation of a spatially concentrated assembly.

We reasoned that, if PTBP1, MATR3, CELF1 or TDP-43 control the Xi-accumulation of *Xist*, loss of the E-repeat should disrupt XCI by reducing *Xist* enrichment within the X-chromosome territory. In support of this hypothesis, it has been shown that *Xist* lacking *Xist* exon 7, which contains the E-repeat (Fig. 2a), displays compromised nuclear localization during XCI initiation^32, 33^. To directly test the requirement of the E-repeat in *Xist* localization, we deleted it in a female hybrid mouse ESC line that carries one X-chromosome from the *129* background and a second X-chromosome from the *castaneous* (cas) background. We confirmed that the deletion of the E-repeat occurred on the *129* allele, which also harbors 11 copies of an MS2-RNA tag within *Xist*^15^, yielding the X*_129_*^Xist ΔE, MS2^ X*_Cas_*^Xist WT^ genotype (referred to as ΔE ESCs below) (Fig. 2a and Extended Data Fig. 5a-d). We ensured that ΔE ESCs maintained two X-chromosomes, and differentiated normally, as judged by morphological changes and loss of NANOG expression (Extended Data Fig. 5e-g). When transcribed from the X*_129_*^Xist ΔE, MS2^ allele, *Xist* exon 6 was spliced to a cryptic site within exon 7 to generate an RNA missing specifically the E-repeat (Extended Data Fig. 6).

**Figure 2:**
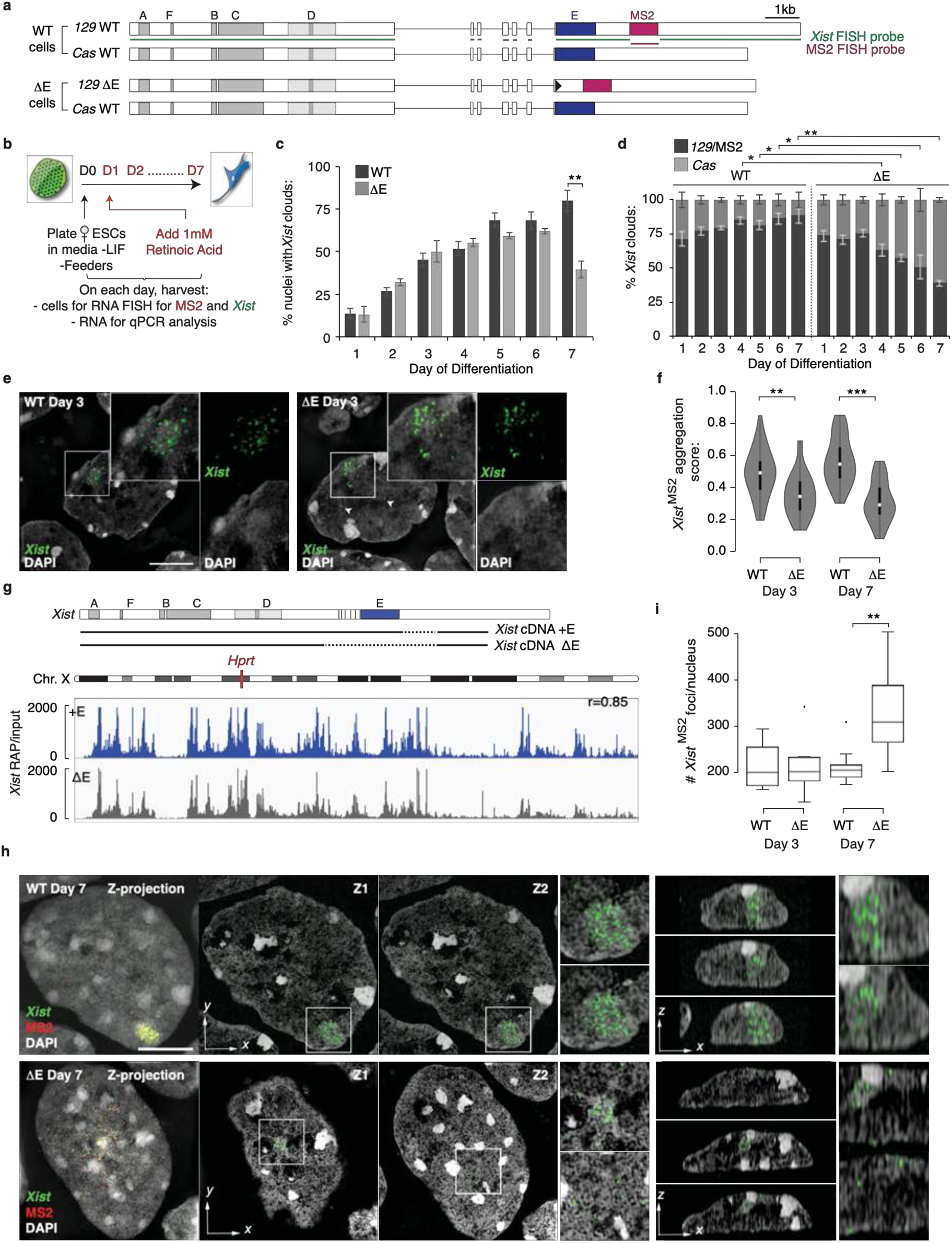
The E-repeat is required for *Xist* localization and controls the number of *Xist* foci. **a,** Diagram showing *Xist* alleles in female WT and ΔE ESCs. A 11xMS2 hairpin, located 1.2kb downstream of the 1.4kb E-repeat, is indicated in magenta. Green lines delineate the sequences within *Xist* RNA targeted by the *Xist* FISH probe. The magenta line represents the region targeted by the MS2 FISH probe. Note that the loxP site (black triangle) present in the genomic sequence after targeting is spliced out in ΔE *Xist* transcripts (see Extended Data Fig. 6). **b,** Scheme of Retinoic Acid (RA) differentiation experiment to assess *Xist* coating of the Xi in female WT and ΔE ESCs. **c,** Histogram showing the number of nuclei that display an *Xist* cloud, detected by RNA FISH against *Xist,* at the indicated day of differentiation. 100 – 250 cells were counted per time point across 3 independent differentiations. Error bars represent SEM (p-values: ** < 0.005, 2-tailed students t-test). **d,** Histogram showing the allelic origin of *Xist* RNA clouds for the data in (c), detected by RNA FISH for *Xist* and the MS2 tag. At later days of differentiation (4-7), MS2+*Xist^129^* (mutant) clouds in ΔE cells were scored if an *Xist* signal (which delivers a brighter signal than the MS2 probe due to target sequence length - see (a)) co-localized with even a weak MS2 signal. 50-80 *Xist* clouds were assessed per time point, across 3 independent experiments. Error bars represent SEM (p-values: * < 0.05, ** < 0.005, 2-tailed students t-test). **e,** 3D-SIM optical sections through WT and ΔE cells at day 3 of differentiation showing *Xist* RNA FISH signals (green) of MS2+*Xist^129^* WT or ΔE clouds, with DAPI counterstaining in grey. Note, that for simplicity, the MS2 signal is not shown, but the *Xist* clouds shown are derived from the MS2+*Xist^129^* allele. Inset: Enlargement of marked region corresponding to the *Xist*-coated-chromosome. Right: Same as inset except DAPI and *Xist* RNA FISH are presented independently, highlighting formation of the Barr body. Bar; 5µm. Arrowheads indicate ΔE-*Xist*^MS2^ foci located further from the core of the *Xist* cloud. **f,** Violin plots of aggregation scores of MS2+*Xist^129^* clouds formed in WT and ΔE cells at days 3 and 7 of differentiation. Only MS2+*Xist^129^* clouds were chosen for the analysis, but the signal quantified was derived from the co-localizing *Xist* RNA FISH signal. Significance testing was performed using a 2-sample K-S test. 30-34 clouds were analyzed per time point from one experiment (p-values: ** < 0.005, *** < 0.0005). **g,** Top: Diagram of the tet-inducible *Xist* cDNA transgenes that were inserted into the *Hprt* locus on the X-chromosome in male ESCs for RAP-seq. Bottom: RAP-seq profile of +E and ΔE *Xist* across the X-chromosome after 6 hours of dox treatment. Pearson correlation score between the two profiles is indicated. Data were collected from one experiment. **h,** Left panel: 3D-SIM MIP of the *Xist* RNA FISH signal from MS2+*Xist^129^* clouds in WT (top panel) and ΔE (bottom panel) nuclei at day 7 of differentiation. *Xist* is shown in green, MS2 in red, DAPI in grey. Bar; 5µm. Next two large panels: *Xist*/DAPI signals from two different Z-plane optical sections through the nucleus are shown. Next two small panels: Enlargements of the *Xist*-coated X-chromosome from each Z-plane (Z1, top; Z2, bottom). Rightmost panels: Y-plane sections through same cells, showing *Xist* localization relative to the apical and basal nuclear lamina with enlargements of area with *Xist* signal area shown on the far right (see Extended Data Figure 8a for additional images). **i,** Box plot showing the distribution of *Xist* RNA foci number derived from the MS2+*Xist^129^*-allele in WT and ΔE cells at day 3 and 7 of differentiation. Significance testing was performed using the 2-sample K-S test. 10 clouds were quantified per condition from one experiment (p-value: ** < 0.005).

RNA FISH over seven days of differentiation (Fig. 2b) revealed that the number of cells containing an *Xist*-coated X-chromosome increased gradually until day 4 of differentiation in both WT and ΔE cells (Fig. 2c). At later time points (days 5-7), we observed a significant decline in the proportion of cells containing an *Xist* enrichment in ΔE compared to WT cells (>50% reduction at day 7) (Fig. 2c). This reduction was specific to the *129*^Xist ΔE, MS2^ allele as revealed by RNA FISH against the MS2 tag (Fig. 2d). Although slightly less than in WT cells, the level of the MS2+ *Xist* transcripts in ΔE cells continually increased during differentiation and RNA half-life was the same for WT and ΔE *Xist* measured at day 3 of differentiation (Extended Data Fig. 7). Therefore, loss of ΔE *Xist* enrichment on the Xi was not an apparent consequence of decreased *Xist* transcript abundance or reduced RNA stability.

Early during initiation of XCI (differentiation day 3), the ΔE *Xist* was observed to enrich over the X-chromosome, while its aggregation measurements revealed a modest, but significant, defect in ΔE *Xist* localization compared to WT *Xist* (Fig. 2e/f). RNA antisense purification (RAP) of *Xist* and sequencing of the associated DNA in male ESCs inducibly expressing full length or ΔE *Xist* revealed highly correlated patterns of interaction across the X-chromosome early after *Xist* induction (6h induction, Pearson, r=0.85, Fig. 2g). This confirms that the E-repeat is not required for the initial transfer of *Xist* across the X-chromosome upon onset of XCI. By day 7, however, ΔE *Xist* was strikingly dispersed within the nucleus, often localizing at the nuclear lamina (Fig. 2h and Extended Data Fig. 8a-d). The ΔE *Xist* aggregation score decreased to less than half of WT by day 7 (p<0.00001, 2-sample ks test) (Fig. 2f and Extended Data Fig. 8e/f). H3K27me3 enrichment on the Xi is an *Xist-*dependent mark at all stages of XCI^17, 26^. Consistent with the *Xist* localization defect later in XCI initiation, we observed lower H3K27me3 enrichment and reduced chromatin compaction over the X-chromosome territory at day 7 of differentiation, despite having been established normally at day 3 of differentiation (Extended Data Fig. 9/10). Notably, the ΔE-*Xist* localization phenotype that becomes starkly apparent between day 4 and 7 of differentiation follows the transition from the *Xist*-dependent to the *Xist*-independent phase of XCI initiation^5^, and coincides with the described decrease in Xi-enrichment of the H3K27me3 methyltransferase complex PRC2^17, 50^ (Extended Data Fig. 11). Together, these results reveal a transition in the mechanisms that enrich *Xist* on the X-chromosome during XCI initiation, from a largely E-repeat-independent to an E-repeat-dependent phase. Moreover, the inability of *Xist* to remain associated with the X-chromosome in the absence of the E-repeat occurs after the induction of transcriptional silencing (see below), and upon transition to the *Xist*-independent-phase of XCI.

The *Xist* RNA signal has been previously resolved into 100-200 individual foci (or granules) when assessed with super-resolution imaging by 3D-SIM^18^. Quantifying the number of foci, we found that MS2+*Xist^129^* clouds at both D3 and D7 in WT cells, as well as the MS2+*Xist^129^* clouds in ΔE cells at D3, were comprised of ∼200 foci (Fig. 2i). However, at D7, we observed a striking 50% increase in the average number of MS2+*Xist^129^* foci in ΔE compared to WT cells (p=0.05, two-sample ks-test) (Fig. 2i). This change occurs after the number of cells with *Xist* clouds in the population has stabilized (Fig. 2c) and while *Xist* transcript abundance is still increasing (Extended Data Fig. 7a). Thus, the increased number of ΔE *Xist* foci at later times may result from disassembly of *Xist* foci that each contain multiple *Xist* transcripts^51, 52^. Taken together, our imaging data support a model in which the E-repeat is required for integration of multiple *Xist* transcripts into individual *Xist* foci and for stabilizing the localization of these foci within the X-chromosome.

The delayed timing of defective ΔE *Xist* localization raised several possibilities regarding gene regulation. Silencing of genes on the ΔE *Xist*-expressing X-chromosome (i) might not occur, (ii) might be initiated but not sustained, or (iii) might be both initiated and propagated normally with ΔE *Xist* dispersal occuring after silencing becomes *Xist*-independent^5^. To distinguish between these possibilities, we examined differentiating WT and ΔE ESCs using RNA FISH to measure nascent transcripts from five X-linked genes known to be transcriptionally silenced by XCI: *Gpc4, Rnf12, Mecp2, Chic1,* and *Atrx* (Fig. 3a/b). The ΔE *Xist* RNA was identified by FISH against the MS2 tag. In this assay, the presence of two nuclear nascent transcript foci is indicative of biallelic expression of the respective X-linked gene. One focus indicates silencing of one allele, which can be correlated to the Xi (*Xist-*expressing allele) by colocalization. In WT cells by day 3 of differentiation, some genes (*Gpc4* and *Rnf12*) already show mono-allelic and others (*Atrx* and *Mecp2*) biallelic expression, consistent with known differences in silencing kinetics (Fig. 3c and Extended Data Fig. 12a-d)^53^. Importantly, during the first three days of differentiation, we observed little difference in the level of gene silencing between WT and ΔE cells for most genes (Fig. 3c (compare top and bottom panels)/3d and Extended Data Fig. 12d), although, silencing of *Tsix,* a repressor of *Xist*^54^, and *Gpc4* were slightly delayed compared to WT (Fig. 3c and Extended Data Fig. 12c-e). Later stages of differentiation (day 4-7) presented a strikingly different pattern. The ΔE *Xist* expressing cells frequently failed to silence the five X-linked genes (Fig. 3c/d and Extended Data Fig. 12c/d). Consistent with this late silencing defect, we observed that RNA-polymerase II (RNA-PolII), which was largely excluded from the ΔE *Xist-*marked territory during early differentiation, was intermingled with the ΔE-*Xist* foci at later times (Fig. 3e and Extended Data Fig. 11b-d). The E-repeat is thus essential for sustaining silencing of X-linked genes and exclusion of RNA-PolII beyond the initial wave of transcriptional shutoff. Moreover, the results demonstrate that the *Xist*-independent state of XCI initiation^5^ is not established in the absence of the E-repeat, suggesting that *Xist* generates the epigenetic memory for the silent state through the E-repeat. These findings uncover a new and crucial layer of regulation during *Xist-*dependent silencing.

**Figure 3:**
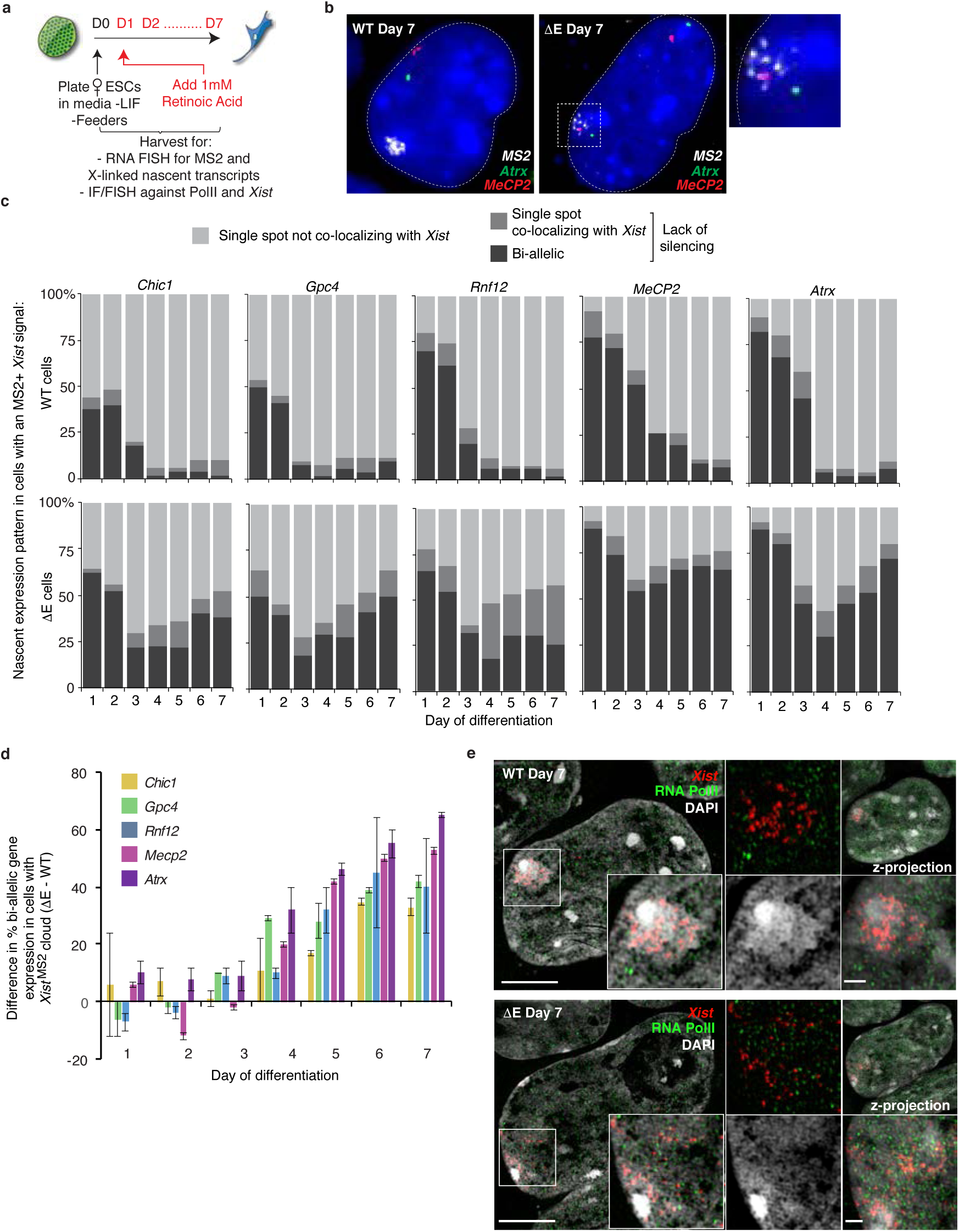
The *Xist* E-repeat establishes a memory of gene silencing during XCI initiation. **a,** Scheme of RA-differentiation experiment assessing gene silencing and Xi RNA-PolII presence in female WT and ΔE ESCs. **b,** Representative epifluorescence images showing the nascent expression patterns of the X-linked genes *Mecp2* (red) and *Atrx* (green) found in a majority of WT (mono-allelic expression on the active X chromosome) and ΔE cells (bi-allelic expression) with MS2+*Xist^129^* signal at day 7 of differentiation, together with the MS2 RNA FISH signal (white). DAPI staining is shown in blue. Note that MS2 signal in ΔE cells is faint as it is dispersed across the nucleus as described in Fig. 2. Far right: enlargement of square hatched region showing the MS2 FISH signal in ΔE-*Xist* cells. **c,** Histograms showing the quantification of nascent expression patterns of indicated X-linked genes in WT and ΔE cells displaying an MS2+*Xist^129^-*coated X-chromosome, across 7 days of differentiation. 50 cells with an MS2+*Xist^129^* clouds were counted per time point. (See Extended Data Fig. 12c/d). **d,** Histogram showing the percentage difference in bi-allelic nascent gene expression (lack of silencing) between ΔE and WT cells with an MS2+*Xist^129^* cloud, across 7 days of differentiation for the X-linked gene expression quantification shown in (c and Extended Data Fig.12d). Error bars represent SEM. **e,** 3D-SIM optical sections through nuclei of WT and ΔE cells showing *Xist* clouds derived from the MS2+*Xist^129^* allele at 7 days of differentiation, stained for RNA-polII (green) and probed for *Xist* (red). DAPI is shown in grey. Inset: Magnification of the marked region. Right, top: Same as inset but only showing PolII and *Xist* signals. Right bottom: Same as inset but only showing DAPI. Far right: z-stack projections of the whole nucleus (top panel) and Barr Body (lower panel). Size bars: 5µm; inset:1µm.

To test a causal relationship between the E-repeat-binding RBPs, *Xist* localization and gene silencing, we synthetically fused PTBP1, MATR3, CELF1, or TDP-43 to the MS2-coat protein (MCP)^55^. This allowed recruitment of these proteins to ΔE-*Xist* RNA during differentiation via the 11xMS2-tag (Fig. 2a, 4a and Extended Data Fig. 13a, c, d). In this gain-of-function approach, we were able to differentiate ESCs for seven days, which was not possible with RBP depletion. We then assessed whether synthetic recruitment of an RBP rescued the phenotypes associated with the loss of the E-repeat. We first confirmed *in vivo* recruitment of the synthetic protein to *129*^MS2^ *Xist* by expressing a Flag-MCP-GFP fusion protein in differentiating WT ESCs (Extended Data Fig. 13b). We found that continued expression of a Flag-MCP-PTBP1 fusion protein in ΔE cells, assessed at day 7 of differentiation, rescued *Xist* localization, gene silencing of *Gpc4* and *Atrx*, and H3K27me3 enrichment on the 129^ΔE,MS2^ X-chromosome at day 7 of differentiation (Fig. 4b-e and Extended Data Fig. 13e). Interestingly, we observed a similar rescue upon expression of Flag-MCP-MATR3, Flag-MCP-TDP-43, or Flag-MCP-CELF1 (Fig. 4b-e and Extended Data Fig. 13e). These data demonstrate that the E-repeat controls *Xist* localization, gene silencing, and heterochromatin formation via its binding proteins PTBP1, MATR3, TDP-43 and CELF1.

**Figure 4:**
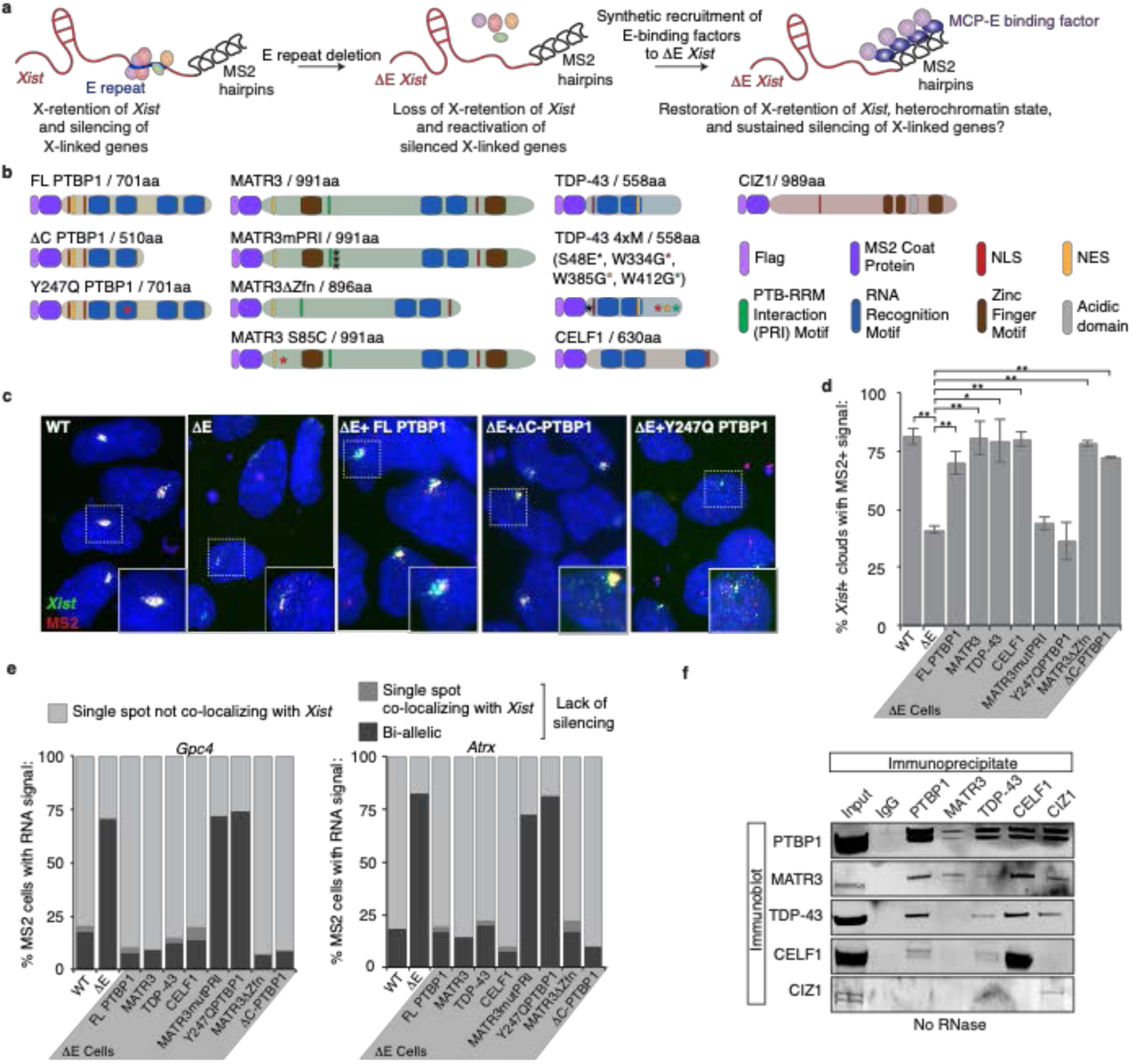
PTBP1, MATR3, TDP-43 and CELF1 confer function on the E-repeat to localize *Xist* and enable persistent X-linked gene silencing during XCI-initiation. **a,** Depiction of the MCP-fusion protein rescue strategy for ΔE *Xist*. **b,** Illustration of Flag-tagged MCP-fusion proteins used in this study and mutants thereof. Point mutations are indicated with asterisks (see methods for MATR3 PRI mutant sequence, denoted by 3 black asterisks). Amino acid length of fusion proteins includes Flag-tag and MCP sequence. **c,** Representative epifluorescence images of RNA FISH against *Xist* (green) and MS2 (red) in day 7 differentiated WT, ΔE, and ΔE lines expressing variants of Flag-tagged MCP-PTBP1 fusions. Inset: enlargement of MS2+*Xist^129^* clouds outlined with hatched lines. Chosen clouds are representative of the MS2+*Xist^129^* cloud phenotype in ΔE cells upon rescue with the indicated protein variant. **d,** Histogram showing the number of nuclei with an *Xist* cloud that also displayed a co-localizing MS2 signal at differentiation day 7, as detected by RNA FISH, in WT, ΔE, or ΔE lines expressing the indicated Flag-tagged MCP fusions (see methods for information on scoring). 83-103 cells were counted per sample. Error bars represent the SEM from two independent experiments (p-values: * <0.05, ** < 0.005, 2-tailed students t-test). **e,** Histograms showing the quantification of nascent *Gpc4* or *Atrx* nascent expression pattern in cells displaying an MS2+*Xist^129^* coated X-chromosome at differentiation day 7 as described in (d). 50 cells with an MS2+*Xist^129^-*coated X-chromosome were counted per sample from one experiment. **f,** Immunoprecipitation of PTBP1, MATR3, CELF1, TDP-43 and CIZ1 from ESC extracts (not RNase treated) and detection of co-precipitated proteins by immunoblotting, using the same antibodies. Image is representative of the results obtained from three independent replicates.

The ability of PTBP1, MATR3, TDP-43 and CELF1 to individually rescue phenotypes associated with loss of the E-repeat raised the question of whether they act redundantly in this process. To examine this more closely, we took advantage of a known direct interaction between PTBP1 and MATR3^44^. We found that Flag-MCP-MATR3 harboring a mutant PTBP1-RRM Interaction (PRI) sequence (MATR3mPRI)^44^, partially rescued H3K27me3 enrichment, but was unable to rescue the *Xist* localization and gene silencing defects observed upon loss of the E-repeat (Fig. 4b,d,e and Extended Data Fig. 13e/f). The converse mutation in PTBP1 (Y247Q) that prevents interaction of PTBP1 with MATR3^44^, was similarly unable to rescue the aggregation and silencing phenotypes associated with loss of the E-repeat (Fig. 4b-e and Extended Data Fig. 13e). These findings are supported by co-immunoprecipitation results demonstrating that PTBP1, MATR3, CELF1 and TDP-43 can interact with one another in the presence of RNA, whereas only PTBP1 and MATR3 robustly co-precipitate after RNase treatment (Fig. 4f and Extended Data Fig. 14). These findings show that PTBP1 and MATR3 are each necessary but insufficient for conferring function on the E-repeat and do not act completely redundantly in this process. Moreover, the data demonstrate that a specific protein-protein interaction between PTBP1 and MATR3 is critical for *Xist* localization, gene silencing and H3K27me3 Xi-enrichment.

The protein CIZ1 was previously suggested to anchor *Xist* to chromatin via the E-repeat in mouse embryonic fibroblasts (MEFs)^31, 56^. Interestingly, PTBP1 and MATR3 interact with CIZ1 in both the presence and absence of RNA in our co-immunoprecipitation assay (Fig. 4f and Extended Data. Fig. 14). However, expression of Flag-MCP-CIZ1 did not rescue *Xist* cloud formation or X-linked gene silencing in our rescue system (Extended Data Fig. 15). Moreover, unlike the CIZ1 Xi-enrichment observed in WT cells (Extended Data Fig. 16) ^31, 56^, CIZ1 did not accumulate on the ΔE-*Xist*-coated X-chromosome in cells expressing Flag-MCP-PTBP1, - MATR3, or -TDP-43 (Extended Data Fig. 16). Despite the RNA-independent interactions observed between CIZ1 and PTBP1/MATR3, these results suggest that distinct functional complexes assemble on the E-repeat and that rescue by PTBP1, MATR3, TDP-43 and CELF1 is independent of CIZ1. As a negative control, we also tested a bivalent MS2-CP-GFP-MS2-CP fusion in our rescue system (Extended Data Fig. 17), which was unable to rescue the ΔE-*Xist* localization and silencing defects. This result suggests that the linkage formed PTBP1 and MATR3 does not function to simply tether *Xist* transcripts together and that additional activities conferred by the recruited proteins likely facilitate compartmentalization of *Xist* and downstream events in XCI.

To define such additional features of the E-repeat-interacting RBPs, we performed a limited domain analysis. MATR3 encodes two zinc finger domains described as conferring DNA binding^57^ (Fig 4b). We found that the rescue of the ΔE-*Xist* phenotypes by MATR3 is independent of these Zinc finger domains (MATR3ΔZfn) (Fig. 4b,d,e and Extended Data Fig. 13e/f). We also tested whether the valency of the RNA binding by PTBP1 affects the ability of the protein to confer function on the E-repeat. PTBP1 lacking two of its four RNA recognition motifs (RRMs) (ΔC-PTBP1 without RRM 3 and 4, Fig. 4b) retains some splicing repression activity^58^ and, in our system, rescued *Xist* cloud formation, H3K27me3 Xi-enrichment, and gene silencing defects observed upon E-repeat deletion (Fig. 4c-e and Extended Data Fig. 13e). However, closer inspection of ΔE *Xist* clouds formed in the ΔC-PTBP1 rescue line, however, revealed the presence of dispersed *Xist* foci within the nucleus, normally not seen for WT *Xist* or in the rescue with full-length PTBP1 (see inset, Fig. 4c). This result indicates that binding valency by PTBP1 is an important parameter in the function of the PTBP1-*Xist* assembly and suggests that PTBP1 likely uses all of its RRMs to build a network of RNA-protein interactions in the Xi (Extended Data Fig. 4).

The formation of the *Xist* territory containing PTBP1 is interesting in light of the observation that PTBP1 can undergo liquid-liquid de-mixing *in vitro*, when incubated at high concentration with a binding RNA^41, 42^. Similarly, MATR3, TDP-43 and CELF1 are components stress granules and paraspeckles, membrane-less RNP granules that form via phase separation of RNA-protein assemblies^38–40, 43^. Therefore, we examined whether features that play a role in these processes apply to XCI. First, we asked whether rPTBP1 can form liquid-droplets upon interaction with the E-repeat of *Xist in vitro*. We found that in the absence of RNA, a solution of rPTBP1 (60uM) remained uniformly clear as assessed by light microscopy (Extended Data Fig. 18a). Addition of the E-repeat RNA at a concentration reported for RNA-PTBP1 droplet formation^42^ (3.2uM), produced RNA-protein aggregates within an hour (Fig. 5a). Lower RNA concentrations (0.1-0.5uM) produced droplets that resembled phase-separated liquids by multiple parameters: (1) upon addition of RNA, the PTBP1 solution became turbid (data not shown), (2) spherical, transparent droplets accumulated over time, (3) these droplets fused with other droplets, and (4) higher concentrations (0.3-0.5uM) of E-repeat RNA lead to larger droplets than lower concentrations (0.1uM) (Fig. 5a/b and Extended Data Fig. 18b). In contrast, very small droplets produced by a control RNA, containing 5 short CU-tracts, showed minmal changes in size between upon addition of 0.1 - 0.5uM of RNA and did not aggregate at the highest concentration of RNA. These findings indicate that the highly multivalent *Xist* E-repeat has a high propensity for condensation with PTBP1^59^.

**Figure 5:**
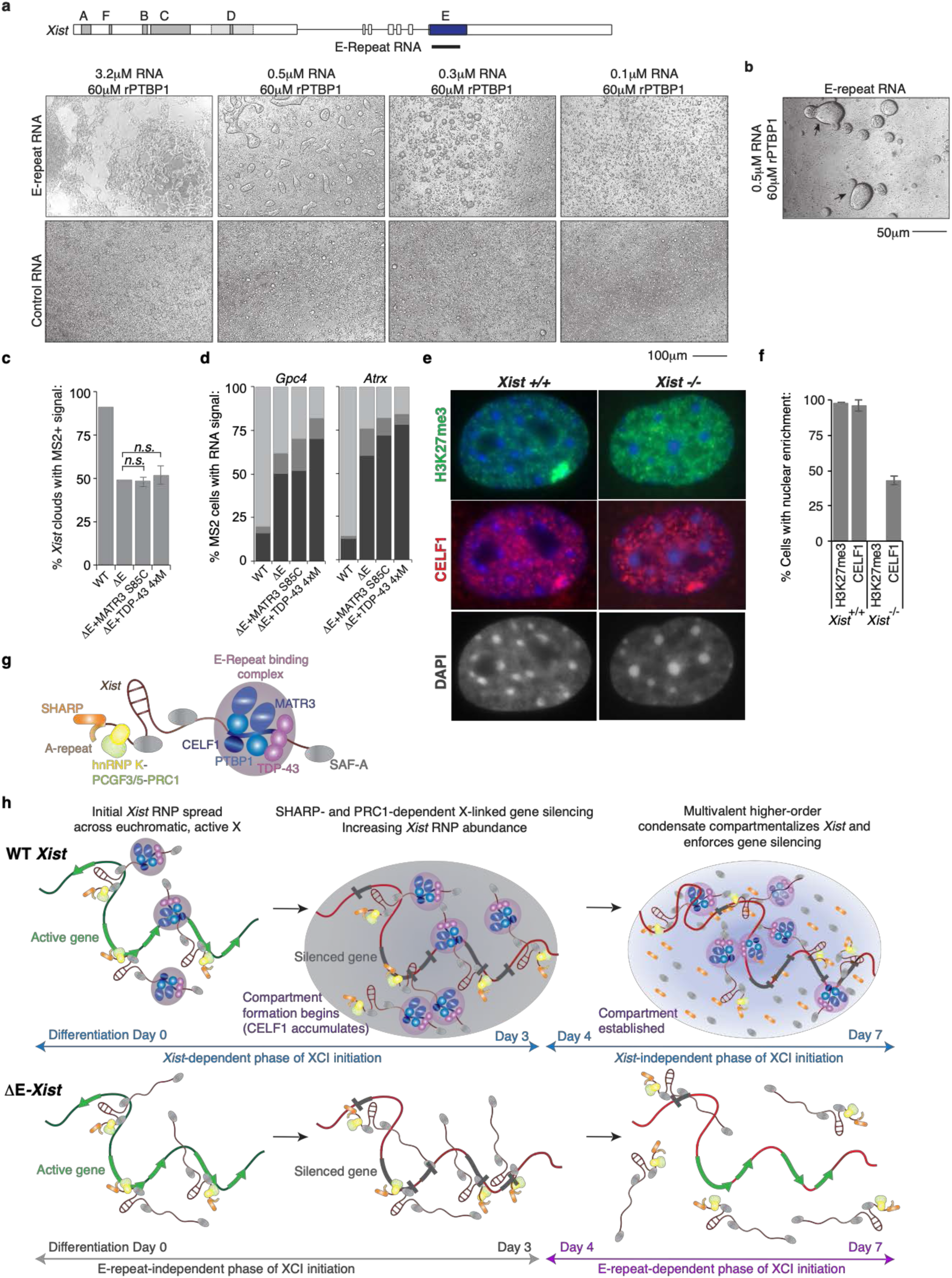
Self-association of E-repeat binding RBPs is critical for compartment formation. **a,** Top: Diagram of the *Xist* locus with introns, exons and repeat arrays A-F depicted as in Fig. 2a. The *Xist*-derived *in vitro* transcribed E-repeat used in droplet assays is indicated. Bottom: Brightfield (10x) images showing droplets formed after 40 minutes with 60μM rPTBP1 and decreasing concentrations of *Xist* E-repeat or control RNAs. **b,** Brightfield image of droplets formed after 40 minutes with 60μM rPTBP1 and 0.5μM E-repeat RNA, which have fused or are undergoing fusion, indicated by arrows. Images were taken at higher magnification than in (a), note scale bar. **c,** Histogram showing the number of nuclei with an *Xist* cloud that also displayed a co-localizing MS2 signal, as detected by RNA FISH, in WT, ΔE, or ΔE cells expressing Flag-tagged MCP-MATR3-S85C and TDP-43 4xM, respectively. 108-144 cells were counted per sample, from 2 independent experiments. Data for WT and ΔE samples were from a single experiment but was combined with data from Fig. 2 to calculate experimental standard error of the mean. There was no significant difference between the ΔE sample and experimental ΔE rescue lines (2-tailed students t-test). **d,** Histograms showing the quantification of nascent *Gpc4* or *Atrx* nascent expression patterns in cells described in (c) displaying an MS2+*Xist^129^* coated X chromosome at differentiation day 7. 50 cells displaying an MS2+*Xist^129^* were counted per sample, from one experiment. **e,** Representative fluorescence microscopy images showing H3K27me3 (green) and CELF1 (red) staining in *Xist*^2lox/2lox^*, Rosa26*^M2rtTA/tetO-Cre-recombinase^ before, or after 96h of dox treatment. Images of the DAPI stained nuclei (grey) are shown below. **f,** Histogram showing the percentage of cells with H3K27me3 or CELF1 enrichment before or after 96h of dox treatment (2ug/mL). Error bars represent SEM from two biological replicates. **g,** Illustration of the *Xist* ribonucleoprotein complex. PTBP1, MATR3, CELF1 and TDP-43 bound to the E-repeat are depicted in blue/purple. The A-repeat bound by SHARP is shown in yellow/brown, and hnRNPK-PGCF3/5-PRC1 bound to the B-repeat indicated in yellow/green. Proteins marked in grey indicate additional *Xist*-interactors, such as SAF-A/hnRNP-U. **h,** Model for how the E-repeat forms the Xi domain and facilitates the transition from the *Xist-* dependent to the *Xist-*independent phases of XCI-initiation. Top: At the onset of differentiation, *Xist* expression is induced and the *Xist* RNP spreads across the euchromatic X-chromosome (green line) using the 3D conformation of the chromosome and mechanisms involving SAF-A/hnRNP U to attach to chromatin. SHARP, bound at the A-repeat, recruits HDAC3 (not shown) to initiate transcriptional shutoff for genes. HnRNPK, bound at the B-repeat, instates heterochromatin formation (red line) via deposition of PRC1-dependent H2AK119ub and PRC2-dependent H3K27me3, and contributes to gene silencing. Further differentiation and likely DNA compaction, along with rising *Xist* levels, increase the overall local concentration of *Xist* RNPs, such that more *Xist* RNPs integrate into each *Xist* granule detected in 3D-SIM experiments. At a critical concentration of *Xist* transcripts and E-repeat-binding RBPs, the formation of a higher-order multi-molecular and multi-valent assembly occurs. This establishes a self-sustaining condensate that compartmentalizes *Xist* within the Xi-domain and enforces X-linked gene silencing, and that can be sustained in the absence of *Xist*. Bottom: Without the E-repeat, the *Xist*-retaining silencing compartment does not form, *Xist* localization cannot be reinforced, rendering the local concentration of *Xist* and its directly interacting proteins insufficient to maintain gene silencing, which ultimately results in reactivation of the Xi.

We also assessed whether self-assembly of TDP-43 affected XCI. TDP-43 forms higher-order complexes that undergo LLPS both *in vitro* and *in vivo*^60–62^ and this activity is reduced by mutations: S48E, W334G, W385G and W412G^60–62^. If formation of the Xi-domain also involves TDP-43 condensation, a mutant TDP-43 harboring these mutations (TDP-43-4xM, Fig. 4b) should be incapable of rescuing ΔE-*Xist* associated phenotypes during XCI initiation. Indeed, unlike WT TDP-43 (Fig. 4d), Flag-MCP-TDP-43-4xM did not restore *Xist* localization or gene silencing upon deletion of the E-repeat (Fig. 5c-d and Extended Data Fig. 19). Similar results were obtained with a MATR3 S85C mutant, previously shown to impair the formation of stress granules that also form from condensation ((Fig. 5c-d and Extended Data Fig. 19)^63–66^. From these data, we posit that by binding these proteins at high density via the E-repeat, *Xist* effectively concentrates PTBP1, MATR3, TDP-43 and CELF1 to establish a physical condensate, that matures over time during the initiation of XCI and relies on self-aggregation properties to compartmentalize *Xist* and enforce X-linked gene silencing during the subsequent *Xist*-independent phase of XCI.

Our results suggested that the condensate set up by the E-repeat could be retained in the absence of *Xist* at least through a few rounds of cell division, and this could explain how the *Xist-*independent phase of XCI initiation is achieved when *Xist* is deleted after day 3 of differentiation^5^. To test the possibility that the protein condensate might be retained in the absence of *Xist*, we immunostained CELF1 in primary female MEFs carrying a conditional *Xist* allele^67^ (Fig. 5e) to detect the E-repeat dependent condensate. As seen with differentiating ESCs, in the WT MEFs we observed CELF1 enrichment that co-localized with the H3K27me3 accumulation over the Xi (Fig. 5e). We then deleted *Xist* in the MEF cells using Cre-mediated recombination (Extended Data Fig. 20). As expected, loss of *Xist* led to loss of H3K27me3 accumulation on the Xi^17, 26^ but strikingly, in half of the *Xist-*deleted cells, CELF1 maintained the punctal enrichment (Fig. 5f). We conclude that the E-repeat-seeded protein condensate is stable without *Xist*, which likely explains how silencing can be enforced during the *Xist-*independent phase of XCI.

Our results define a new model for how *Xist* establishes the Xi-domain during XCI initiation (Fig. 5g/h -top). Upon induction of differentiation, *Xist* is upregulated, assembles with proteins across the RNA (Fig. 5g), and spreads along the X-chromosome^24^ (Fig. 5h, top left). Previous studies have elucidated funtions for several of these proteins. SAF-A/hnRNP-U mediates the chromatin attachment of *Xist*^68^, while the proteins SHARP^25^, bound at the A-repeat, and PRC1 recruited via hnRNP-K binding to the B-repeat^29^, silence transcription (Fig. 5h, top middle). We now define a function for the *Xist* E-repeat that recruits the RNA binding proteins PTBP1, MATR3, TDP-43 and CELF1 during the initiation of XCI. These factors each carry multiple RRMs allowing for the simultaneous engagement of distinct repeat motifs within in the long E-repeat sequence, whose multivalency will increase the avidity of binding to a single transcript (Fig. 5h, top middle). Our results also define homotypic and heterotypic interactions between these factors required for the faithful initiation of XCI. Together, these multivalent RNA-protein and protein-protein interactions will form a higher-order *Xist*–protein network. We propose that increasing *Xist* abundance with differentiation and, likely, compaction of the X-chromosome^18^ concentratrate the *Xist* binding factors within the confined nuclear region of the Xi and result in condensation of PTBP1, MATR3, TDP-43 and CELF1 around the nucleating *Xist* molecules.

After day 3 of differentiation and XCI initiation, the condensate formed by the E-repeat binding proteins is critical for XCI and sustained silencing of X-linked genes during the *Xist-* independent phase of XCI initiation (Fig. 5h, top right). At this point, the E-repeat has led to coalescence of *Xist* transcripts into the *Xist* granules and compartmentalization of the *Xist*-coated Xi. By binding and concentrating the factors needed for establishment of the Xi-domain, *Xist* enforces its own *cis-*limited spread and resultant gene silencing. The formation of the E-repeat-dependent condensate can allow *Xist* to remain associated solely on the X chromosome from which it is expressed. In the absence of the E-repeat, the loss of gene silencing and the dissociation of *Xist* from the X-chromosome domain only occur after trancriptional shutoff and heterochromatin formation (Fig. 5h bottom). We suggest that gene silencing, loss of active transcriptional regulators, and heterochromatinization alter the interaction of *Xist* with the Xi and induce a transition from an E-repeat-independent mode of association to one that is E-repeat-dependent.

Our model also suggests a mechanism for the epigenetic memory that perpetuates the silent state after the inducing molecule (*Xist*) has been deleted. We propose that continued gene silencing upon deletion of *Xist* after day 3 of differentiation^5^, is mediated by the E-repeat-seeded protein condensate. This is consistent with our finding that CELF1 enrichment on the Xi can be maintained in the absence of *Xist*. We hypothesize that the condensate integrates additional *Xist*-interacting proteins, such as SHARP, via specific protein interactions (Fig. 5h top/bottom right). Weak interactions between these different proteins might permit them to diffuse within the Xi-domain. In this way, *Xist*-interactors, such as SHARP, maintain association with the multi-molecular assembly independently of direct *Xist* interaction (Fig. 5h top right). Such a model could explain how 100-200 *Xist* granules (foci)^18, 52^ can silence >1000 genes across the 167Mb of X chromosome DNA. A full understanding of how the condensed silencing domain controls gene silencing will involve determining all of its components and their stoichiometry within the Xi as well further biophysical characterization of the condenstate. It will be particularly interesting to examine individual genes within the compartment and whether silenced or escaper genes exhibit different interations with the condensate or locations within it.

Our silencing domain model may also explain the finding that *Xist-*dependent silencing can only be triggered within a defined developmental window upon onset of XCI. PTBP1, MATR3 and CELF1 are highly expressed in ESCs and decline in abundance upon differentiation^69^. In differentiated cells, the lower levels of these RBPs may be insufficient to multimerize on *Xist* and condense into a silenced compartment.

Our work provides a new way of thinking about the mechanism of XCI, where the condensation of *Xist* with its interacting proteins drives the compartmentalization needed for sustained gene regulation. Our results also reveal how RBPs, known for their roles in RNA processing, mediate lncRNA localization and exert control over gene regulation via mechanisms independent of their previously described RNA processing activities.

## Author contributions

K.P., A.P-J, Y.M., and D.L.B conceptualized the project, A.P-J., Y.M., J.S., T.C., W.M., A.D., B.P., K.C., R.M., X-J.W., C-K.C., and A.C. acquired data, A.P-J., J.S., Y.M., T.C., H.L. and K.P., analyzed data, A.P-J, Y.M., J.S., M.G. and K.P. interpreted the data and contributed towards methodology and model creation, K.P., D.L.B, M.G. and H.L. acquired funding to support the project, A.P-J. and K.P. administered the project and A.P-J. and K.P. wrote the manuscript and included edits from all authors.

## Author Information

Authors declare no competing interests.

## Data and materials availability

All genomic data for Xist interactions and chromatin association will be deposited in the Gene Expression Omnibus (GEO) database before publication of the manuscript. All reagents will be made available upon request after the manuscripts has been accepted for publication.

## Materials and Methods

### Cell Culture

All mouse ESC lines were cultured in knockout DMEM (Life Technologies) supplemented with 15% FBS (Omega), 2 mM L-glutamine (Life Technologies), 1× NEAA (Life Technologies), 0.1 mM Beta-Mercaptoethanol (Sigma), 1xPenicillin/Streptomycin (Life Technologies), and 1000 U/mL murine LIF (homemade) on 0.3% gelatinized plates (porcine skin gelatin, Sigma) pre-plated with irradiated male DR4 feeders (homemade from day 14.5 embryos with appropriate animal protocols in place). For 3D-SIM microscopy experiments (see below), ESCs were maintained in 2i culture conditions. No differences in results upon cell differentiation (see below) were observed between the ESC propagation conditions. ESCs were maintained as small colonies and passaged with trypsin and single cell dissociation at 80% confluency.

### Female ESC Differentiation

Female WT F1 2-1 MS2*^129^* ^70^(and derivatives thereof) were trypsinized to single cells and counted. Cells were seeded in 2mL of mouse embryonic fibroblast (MEF) medium [DMEM (Invitrogen) supplemented with 10% FBS (Omega), 2 mM L-glutamine (Life Technologies), 1× NEAA (Life Technologies), 0.1 mM Beta-Mercaptoethanol (Sigma) and 1xPenicillin/Streptomycin (Life Technologies)] at a density of 20,000 - 200,000 cells/4cm^2^ (depending on the experiment) on tissue culture plates for Western Blotting or onto 18mm sterile glass coverslips for IF/FISH experiments, both of which were pre-coated with sterile 0.3% gelatin (porcine skin gelatin, sigma) or matrigel (Corning)(diluted 1:100). 24 hours post-seeding, the culture medium was changed and supplemented with 1μM all-trans Retinoic Acid (Sigma), which was changed daily thereafter until the cells were harvested for analysis.

### Female MEF culture

Female MEFs (*Xist*^2lox/2lox^, *Rosa26^M2rtTA/tetO-Cre-Recombinase^*)^67^ were maintained in MEF medium as described above. To delete *Xist,* cells were treated with 2μg/mL doxycycline (sigma) for 96h.

### RNA Fluorescence In Situ Hybridization

FISH against *Xist* RNA was performed using both RNA and DNA probes. FISH against MS2, *Atrx*, *Gpc4, Mecp2, Rnf12* and *Chic1* was performed using DNA probes. In undifferentiated ESCs, the DNA probe against *Xist* additionally detects *Tsix*.

#### RNA Probe Preparation

Strand-specific RNA probes were generated using a T3 *in vitro* transcription kit (Promega) in the presence of Chromatide AlexaFluor-UTP (ThermoFisher). Six ∼700nt transcription templates were generated from *Xist* exon 1 (Primers UCLA 1416 – 1429, Extended Data Table 1), and used in transcription reactions containing 0.5mM ATP, CTP, GTP, 0.1mM UTP, and 0.05mM Chromatide AlexaFluor488-UTP (Life Technologies) along with 1x T3 transcription buffer supplemented with 10mM DTT, 500U RNase inhibitor, 170U T3 RNA polymerase and 5μg of pooled template DNA in a final volume of 500μL at 37°C overnight in the dark. The transcription reaction was treated with 15U RNase-Free DNase for 15 minutes at 37°C prior to probe purification. To purify the probes, 1/3 of the transcription reaction was loaded on a pre-spun (700xg, 5minutes) Chromaspin-100 column (Clontech) and centrifuged (700g, 5 minutes). The eluates were combined and precipitated with 100% EtOH in the presence of 100mg tRNA and 1/10^th^ volume of sodium acetate (Sigma). We sometimes also purified RNA probes using a 2.5x volume of AMPure beads (ThermoFisher 09-981-123, reconstituted according to ^71^), which were washed twice on a magnet with 80% ethanol before elution of the probes from the beads with 50μL water, followed by ethanol precipitation. The RNA pellet was washed twice in 70% ethanol, resuspended in 400μL of RNase Free water, to which 1mL EtOH was added for storage at −20C. To make the final probe, 1/7^th^ of the Probe/EtOH solution was added to 90μL Salmon Sperm DNA (Sigma), 90μL mouse Cot1 DNA (Life Technologies), 40μL 3M RNase Free Sodium Acetate (Sigma), 40μL 10mg/mL tRNA (Life Technologies) and 1mL EtOH. After vigorous shaking, the solution was centrifuged at maximum speed for 10 minutes. The pellet was washed once with 70% EtOH and then once with 100% EtOH, allowed to dry completely, and then resuspended in 200μL deionized Formamide (VWR) and 200μL 2x Hybridization buffer [20% dextran sulfate (sigma), 4x SSC (Ambion), 0.1M NaH_2_PO_4_]. Probes were stored at −80C and denatured at 95°C for 5 minutes before use.

#### DNA probe preparation

For 3D-SIM and Airyscan experiments, FISH probes were labelled by nick translation as described in ^72^, using p15 cDNA plasmid as template and home-labelled Atto488-, Cy3- or Texas Red-conjugated dUTPs made as described in ^73^. For all other experiments, DNA probes were synthesized using the CGH Bioprime Array Kit (ThermoFisher) according to manufacturer’s instructions. Briefly, a 40μL solution containing 100ng of template plasmid DNA was denatured in the presence of 1x random primers at 95°C for 5 minutes and snap cooled on ice. 5μL of nucleotide mix, 5μL of 488, 555-, or 594- dUTP or dCTP chromatide fluorphore (Life Technologies) and 5U Klenow exo-enzyme were then added and incubated in the dark at 37°C for 6 hours, after which an additional 5U of Klenow Exo- were added. The reaction was incubated at 37°C overnight, quenched with 10μL stop solution, and then purified over a Chromatide-100 column or AMPure beads as described above. The eluate was precipitated in the presence of 100mg yeast tRNA (Life Technologies) and sodium acetate (Sigma). The DNA probe was then prepared as above to yield 400μL of probe solution in formamide/hybridization buffer.

The MS2 DNA template for DNA probe preparation was PCR amplified from genomic DNA purified from WT F1 2-1 MS2*^129^* female ESCs (see Extended Data Table 1 for primers). For *Xist*, the DNA probe was synthesized using a full-length mouse *Xist* cDNA plasmid (p15A-31-17.9kb *Xist*, unpublished). Probes against X-linked genes were synthesized using BACs RP23-467J21 (*Gpc4*), RP23-265D6 (*Atrx*), WIBR1-2150D22 (*Chic1*), WIBR1-2704K12 (*Rnf12*) and W11-894A5 and W11-1189K18 (*Mecp2*) (all obtained from CHORI-BACPAC). Note that the use of 2 BAC’s for the *Mecp2* probe sometimes resulted nascent FISH signals that appeared as doublets (see Fig. 3b).

#### RNA FISH procedure for Epifluorescence microscopy

Culture medium was changed 10 minutes prior to harvesting cells to remove dead cells and stimulate transcription. Upon collection, culture medium was aspirated and coverslips were gently rinsed twice with cold 1xPBS. Coverslips were then transferred to a new culture dish containing cold 1xPBS, which was then aspirated, and the cells were fixed in 4% paraformaldehyde (PFA) (Electron Microscopy Sciences) in 1× PBS for 10 minutes at room temperature (RT) under standard laboratory safety practices. After fixation, the cells were permeabilized in 0.5% Triton X-100 (Acros) in 1xPBS with 2mM Vanadyl Ribonucleoside Complex (NEB) for 10-20 minutes on ice. Coverslips were then stored in 70% ethanol at −20°C for 1 hour or until samples from all time points had been collected. Prior to hybridization with probe, the coverslips with cells were brought back to 4°C and serially dehydrated by 5-minute incubations in ice-cold 80%, 95% and 100% ethanol. Coverslips were removed from 100% ethanol and allowed to air dry prior to incubation with probe for 48 hours at 37°C in a sealed chamber humidified with 2xSSC / 50% Formamide. For RNA probes, coverslips were washed 3x 5 minutes in 50% formamide (Fisher) / 2xSSC (Ambion) and 3x 5 minutes in Wash Buffer II (10mM Tris, 0.5M NaCl, 0.1% Tween-20), prior to a 45 minutes incubation with 25 µg/mL RNaseA (ThermoFisher) in Wash Buffer II at 37°C. After RNaseA treatment, coverslips were washed 2x 5 minutes in Wash Buffer II, 2x 5 minutes in 50% formamide / 2× SSC, 3x 5 minutes in 2× SSC and 3x 5 minutes in 1× SSC before briefly drying excess 1× SSC off and mounting with Vectashield mounting media lacking DAPI (Vector Labs). Coverslips were sealed with Biotium Covergrip coverslip sealant (Thermo Fisher). For DNA probes, coverslips were washed 3x 5 minutes in 50% Formamide / 2xSSC, 3x 5 minutes in 2xSSC and 3x 5 minutes in 1xSSC prior to mounting. A 1:10,000 dilution of DAPI (0.5mg/mL) was included in all penultimate 1xSSC washes. All washes were conducted at 42°C, cells were protected from light. All procedures were performed, and used reagents disposed of, according to standard laboratory safety procedures.

#### RNA FISH procedure for 3D-SIM and Improved-Resolution microscopy

All coverslips were processed according to ^74^.

### Indirect Immunofluorescence staining

The cell culture medium was changed 10 minutes prior to harvesting. Upon collection, culture medium was aspirated and coverslips were gently rinsed twice with cold 1xPBS. Coverslips were then transferred to a new culture dish containing cold 1xPBS. If cells were CSK treated (MS2-CP-GFP expressing wild-type WT F1 2-1 MS2*^129^* ESCs), then coverslips were gently treated with 1mL (added dropwise) ice-cold CSK buffer [100mM NaCl, 300mM sucrose, 3mM MgCl_2_, 10mM PIPES pH 6.8] and incubated on ice for 30 seconds before aspiration. Coverslips were then similarly treated with 1mL ice-cold CSK-Trt Buffer (CSK+0.5% Triton X-100) for 30 seconds, followed with a second ice-cold CSK treatment. Coverslips were then processed as described in^75^. See Extended Data Table 2 for antibody information.

### Indirect Immunofluorescence staining -RNA Fluorescence In Situ Hybridization

Where immunostaining and FISH were combined, immunostaining preceded FISH.

#### Epifluorescence Microscopy

The immunostaining protocol was followed as outlined above, but coverslips were not mounted. Instead, after the last round of washes (omitting DAPI in the penultimate wash), coverslips were re-fixed in 4% PFA in 1× PBS for 10 minutes at room temperature and then dehydrated through a 70-85-95-100% ice-cold ethanol series prior to overnight incubation with probe as described above in the RNA FISH procedure section.

#### 3D-SIM and Improved-Resolution microscopy

All coverslips were processed according to ^74^.

### Plasmid Construction and Cell Line Generation

#### Xist-ΔE targeting construct

To create the targeting vector pCR2.1-Puro-*Xist*ΔE, 3kb upstream and 1.2kb downstream of the mouse *Xist* E repeat were PCR amplified from mouse genomic DNA using primers WRM163-166, modified for In-Fusion cloning (Clontech) using Kapa polymerase (Kapa biosystems) according to the manufacturer’s instructions. The upstream homology arm was integrated at the *EcoR*1 site and the downstream homology arm at the *BamH*1 site, in a 4-piece InFusion cloning reaction, into a vector containing a floxed puromycin resistance cassette (PCR2.1-loxP-pGK-Puro-pA-loxP). Positive recombinants were identified by restriction digest of with HindIII.

#### E-repeat deletion in WT F1 2-1 MS2^129^ ESCs

The *Xist* E repeat was deleted in female WT F1 2-1 MS2*^129^* ESCs derived from an F1 cross of mice from pure bread *129* and *castaneous* background, and then targeted to contain a 11x tandem repeat of the MS2 hairpin located 1.2kb downstream of the E-Repeat ^70^ via homologous recombination. The cells also harbor an M2-reverse tetracycline TransActivator cassette within the Rosa26 locus that confers neomcyin resistance on the cells. ½ of a confluent T75 flask of WT F1 2-1 MS2*^129^* female ESCs were electroporated with 40μg of PciI linearized PCR2.1-Puro *Xis*tΔE targeting plasmid (800v, 0.2ms, 4mm cuvette, Biorad X-Cell electroporation module) and plated at varying dilutions on 10cm plates of confluent irradiated DR4 feeders. 36 hours after plating, the cells were selected with 1μg/mL puromycin for 10 days. 100 clones were picked, expanded and subjected to southern blot analysis using a Sac1 digest and an external probe (amplified using primers WRM193/194 (Extended Data Table 1)) as outlined in Extended Data Fig.5. The positive clone #35 was expanded in culture, then transfected with a Cre-recombinase plasmid using Lipofectamine 2000 according to the manufacturer’s protocol (Thermo Fisher), to delete the puromycin resistance cassette. Transfected cells were serially diluted, 100 clones were picked, expanded and replica plated for growth in the presence or absence of puromycin. Sub-clone #96 was sensitive to puromycin. PCR analysis of genomic DNA confirmed the deletion of the puromycin cassette with primers APJ439/440 (Extended Data Table 1). Subsequent southern blot analysis and sequencing of wild-type *Xist* and ΔE *Xist* PCR amplicons from genomic DNA (Intron 6 to Exon 7 using APJ248/631 (Extended Data Table 1)) showed that the ΔE targeting construct integrated on the *129* allele of *Xist* upstream of the MS2 tag, preserving the 3’ splice site of intron 6, to yield the heterozygous E-repeat deletion ESC line X*_129_ ^Xist^* ^ΔE,MS2^ X*_Cas_ ^Xist^* ^WT^ (=ΔE ESCs) (Extended Data Fig. 5/6 and data not shown). Sequencing of the exon 6 - exon 7 RT-PCR amplicon (obtained from cDNA of differentiated ESCs) derived from the *129*^MS2^ *Xist* transcript, revealed the use of a cryptic 3’ splice site downstream of the LoxP site (Extended Data Fig. 6). This extended the E-repeat deletion within the *Xist* transcript (as initially designed) by 42nt and removed the LoxP site and additional vector sequences present in the genomic DNA from mature ΔE *Xist* transcripts, resulting in a scar-less ligation of the 3’ terminus of exon 6 to nucleotide 1479 of Exon 7 (Extended Data Fig. 6 and data not shown). We ensured that ΔE ESCs maintained two X chromosomes throughout the targeting process, and differentiated equally to wild-type as judged by changes in morphology, loss of NANOG upon induction of differentiation (Extended Data Fig. 5).

#### Engineering of WT and ΔE ESCs with a□FLP-FRT recombination platform for rescue experiments

WT F1 2-1 MS2*^129^* ESCs and ΔE ESCs described above (½ of a confluent T75 flask) were electroporated with 40μ<u>g</u> of Fsp1 linearized Flp-IN homing plasmid that integrates a FRT landing site downstream of the Col1A locus and carried a puromycin resistance cassette for targeting ^76^, at 800v, 0.2ms, 4mm cuvette using a Biorad X-Cell electroporation module before being serially diluted on 10cm plates, pre-coated with irradiated DR4 feeders. 36 hours after plating, the cells were selected with 2μg/mL puromycin for 10 days after which 200 clones were picked and expanded. Genomic DNA was isolated and EcoRI digested, before being subjected to southern analysis with the Col1A Xba/Pst1 3’ probe. Positive clones 1-61 (WT) and 137 (ΔE) were used for all subsequent experiments.

#### Generation of Flp-In plasmids encoding Flag-MS2-CP fusion proteins

The MS2 Coat Protein (CP) coding sequence was PCR amplified with a forward primer encoding a 3xFlag tag downstream of a Kozak-ATG start signal. The reverse primer contained an in frame Nhe1 site (primers APJ526/570 (Extended Data Table 1)) such that any fragment ligated into the site would be expressed in frame with the MS2-CP protein, separated by a 3-amino acid (Gly-Leu-Gly) linker. The Flag-MS2-CP-Nhe1 fragment was inserted into the EcoRI site of the pBS32 vector using Infusion cloning. This vector is similar to the pgkATGfrt vector described in^76^, except that the tet-inducible promoter was replaced with a CAGGS promoter, allowing constitutive expression of the fusion protein. The coding sequence for each protein (GFP, PTBP1, MATR3, TDP-43, and CELF1) fused to the Flag-MS2-CP was PCR amplified from cDNA with infusion overhangs, or synthesized (see below, Genewiz), and ligated into the Nhe1 site of the pBS32-Flag-MS2CP parent plasmid using InFusion cloning (Clontech). The PTBP1 Y247Q, MATR3mutPRI and MATR3ΔZfn mutants were generated using primer-directed mutagenesis. The WT PRI sequence (GILGPPP) was mutated to create the mutant PRI sequence (GAAAPPA)^44^. The coding sequences for the CELF1, MATR3 S85C, TDP43 4xM and MS2CP-GFP-MS2CP fusions were synthesized (Genewiz). All plasmids were verified by sequencing.

The ΔC-Terminal PTBP1 fragment that is fused to Flag-MS2-CP in our rescue system, is comprised of the first 299 amino acids of PTBP1, which includes the first two RRMs as well as the MATR3 interaction site, followed by 68 amino acids that are out of frame, and do not encode a functional linker region. A premature stop codon terminates the protein at residue 367.

#### Generating WT and ΔE ESCs expressing Flag-MS2-CP-fusions via Flp-In recombination

33μg of the pBS32 plasmid DNA encoding the Flag-MS2-CP-Fusions and 26μg of plasmid encoding the flpase FlpO were electroporated into WT ESCs carrying the FRT homing site (clone 1-61) for the GFP fusion and ΔE ESCs with the FRT homing site (clone 137) for all other fusion constructs (1/2 of a confluent T75 flask of ESCs per electroporation). Cells were plated on confluent irradiated DR4 feeders in a 10cm dish and 36 hours after plating selected with 170μg/mL hygromycin for 14 days, after which all colonies were picked and expanded. The resulting clones were tested for protein expression by immunoblot of lysates (RIPA buffer in 1X SDS lysis buffer (ThermoFisher)) using an anti-Flag antibody as well as antibodies against the respective fusion protein (Extended Data Table 2). Immunostaining confirmed nuclear localization of all fusion proteins that failed to rescue the phenotypes associated with loss of the E-Repeat. All clones used maintained two X chromosomes, as determined by FISH against *Tsix* in undifferentiated cells. For all rescue experiments, at least two clones were analyzed, which revealed that the data are robust. Therefore, in most cases only the results from one rescue clone per protein or mutant are shown.

#### Generation of tet-inducible Xist ΔTsix V6.5 male ESCs

Tet-On *Xist* male V6.5 ESCs carrying a tet-inducible promoter in place of the endogenous *Xist* promoter and a M2rtTA trans-activator as well as puromycin resistance in the R26 locus^24^ (½ of a confluent T75 flask) were electroporated with 40μg of Not1 linearized paa2Δ1.7 plasmid DNA (800v, 0.2ms, 4mm cuvette using a Biorad X-Cell electroporation machine) and plated on confluent irradiated DR4 feeders. 36 hours after plating, the cells were selected with Neomycin/G418 for 10 days after which 100 clones were picked and subjected to southern analysis described in^24^ (data not shown). Positive clone 70 was used for the PTBP1 ChIP-seq experiments.

### siRNA treatments

Silencer Select siRNAs (Thermo Fisher) against PTBP1, MATR3, CELF1, TDP-43 and SAF-A were diluted to 20nM in 1x siRNA buffer (60mM KCl, 6mM HEPES pH 7.5 0.2mM MgCl_2_), aliquoted and stored at −80C until further use. Under sterile conditions at room temperature, 2.5μL of 20nM siRNA were added to 80μL of fresh Opti-MEM solution (Gibco). 1.6μL siRNA MAX transfection reagent (Life Technologies) were added to 80μL Opti-MEM solution and subsequently added to the siRNA/opti-MEM solution after 5 minutes of incubation. The resulting solution was mixed by pipetting and left to incubate at RT for 20 minutes. The solution was then added to 200,000 cells in 0.8mL of culture medium and plated in 1 well of a 12 well plate on 18mm gelatinized coverslips and left overnight at 37C. For female ESCs undergoing differentiation, cells were plated in MEF medium and after 24h, the culture medium was changed (with the addition of 1μM all-trans Retinoic Acid)(Sigma) and a second round of siRNA treatment was performed. Knockdown efficiency was assessed by immunoblotting (Extended Data Table 2).

### Immunoblotting

Cells were harvested by trypsinization, pelleted (1000xg, 5 min), resuspended in 500μL 1xPBS to wash, and re-pelleted. The washed cell pellet was lysed in 5 pellet volumes of RIPA buffer and 40U benzonase (Novogen) and incubated at 4C overnight. The lysate was centrifuged at max speed to pellet the remaining insoluble material and the supernatant was transferred to a new tube and mixed with 4x Novex sample buffer containing 5% 14.3M Beta-mercaptoethanol (Sigma) to a final concentration of 1x. The samples were then denatured for 5 min at 95C and loaded onto a 4-12% Novex Bis-Tris acrylamide gel with 1x MES running buffer (Life Technologies) run at 120V for 1.5-2h. The gels were transferred to a protran BA-85 nitrocellulose membrane (Whatman) using a Novex XCell II transfer system for 1h at 30V, 4C (or overnight at 4C at 10V) in transfer buffer (25mM Tris-HCl, 192mM glycine, 20% methanol). Membranes were probed with primary antibody (Extended Data Table 2) in 1x Odyssey blocking buffer (LI-COR) overnight at 4C, washed 3×5 minutes in PBS+0.2% tween-20 and then incubated with appropriate secondary antibodies (1:10,000 dilution, Odyssey 700 and 800nm antibodies) in the dark at room temperature for 30 minutes before being washed again and scanned on a LI-COR infrared imaging system.

### Co-Immunoprecipitation

For co-immunoprecipitation experiments, Rabbit IgG and antibodies against PTBP1, MATR3, CELF1, CIZ1, TDP-43 (Extended Data Table 2) were crosslinked to ProteinG-Dynabeads (ThermoFisher) using the protocol provided by Abcam (http://www.abcam.com/protocols/cross-linking-antibodies-to-beads-protocol) with minor modifications. Briefly, 20μL of bead slurry were isolated on a magnet and washed 3x 5 minutes at room temp in 5 volumes of 1xPBS. Beads were then washed once in 5 volumes of binding buffer (100μL, 1xPBS containing 1mg/mL of BSA (NEB)) for 10 minutes at RT, and incubated in 100μL binding buffer supplemented with 5μg of Rabbit IgG or antibodies against PTBP1, MATR3, CELF1, CIZ1 or TDP43. Samples were rotated for 1 hour at 4°C. Beads were then washed in binding buffer for 5 minutes, followed by an additional 5 minute wash in 1xPBS. Next, the antibody was crosslinked by incubating in a 100μL of 1xPBS solution containing 0.2M triethanolamine (Sigma) and 6.5mg/mL Dimethyl pimelimidate (DMP) (Sigma) pH 8.5 for 30 minutes with rotation at RT. Beads were then washed in 250μL 0.2M triethanolamine in 1xPBS for 5 minutes. DMP incubation and wash steps were repeated two more times before samples were quenched in 100μL of 50mM ethanolamine in 1xPBS for 5 minutes. The quenching step was repeated and excess non-crosslinked antibody removed with 2x 10 minutes incubations in fresh 1M glycine pH 3.0. Beads were washed in 1xPBS 3x 5 minutes before use in immunoprecipitations. Immunoprecipitations were performed under non-denaturing conditions according to the Abcam protocol (http://www.abcam.com/ps/pdf/protocols/immunoprecipitation%20protocol%20(ip).pdf). 4×15cm plates of confluent WT F1 2-1 MS2*^129^* female ESCs were lysed by pipetting in 3mL of lysis buffer (10M Tris-HCl pH8, 137mM NaCl, 1% NP40, 2mM EDTA) supplemented with 1x Complete EDTA-free Protease Inhibitors (Roche) and incubated for 1h on ice with or without RNase (10μg/mL RNase A) (Thermo Fisher). Lysate was centrifuged at 4°C, 14,000rpm in a tabletop microfuge for 15 minutes to pellet insoluble material. The supernatant was transferred to new tubes and precleared with 20μL of washed ProteinG-dynabeads per 1mL of lysate with rotation at 4°C for 1 hour. 500μL of lysate were then added to each crosslinked antibody-proteinG Dynabead prep (described above) and rotated at 4°C overnight. The next day, crosslinked antibody-proteinG Dynabeads were isolated on a magnet and washed 4x 5 minutes in ice-cold wash buffer (10mM Tris-HCl pH 7.4, 1mM EDTA, 1mM EGTA, 150mM NaCl, 1% TritonX-100) supplemented with 1x Complete EDTA-free Protease Inhibitors. The co-purified proteins were eluted by boiling in 1x NuPage Protein Loading buffer (Thermo Fisher) supplemented with 5% Beta-mercaptoethanol, at 95°C for 5 minutes. Samples were assessed by immunoblotting. Input represents 4% of lysate added per immunoprecipitate. 1/4^th^ of eluate was loaded per lane.

### *In Vitro* RNA transcription (IVT)

For several *in vitro* experiments (Droplet assays, EMSA), RNAs encoding the E-repeat and other sequences were obtained by IVT. Templates for IVT were amplified from DNA using KAPA polymerase according to manufacturer’s instructions (KAPA Biosystems), and then gel purified and concentrated over AMPure beads (homemade). See Extended Data Table 1 for primer information. RNA was transcribed and UREA-PAGE purified as described in ^77^. For biotinylated RNAs, Biotin-UTP (Ambion) comprised 18% of the total UTP.

### Droplet Assays

Recombinant 6x-His tagged PTB was purified using Ni-NTA agarose (Invitrogen) according to manufacturer’s instructions. The purified protein was dialyzed and stored in Buffer DG (20 mM HEPES-KOH pH 7.9, 80 mM K. glutamate, 20% glycerol, 2.2 mM MgCl_2_, 1 mM DTT, and 0.1 mM PMSF). 10μL droplets were assembled in 1.5mL Eppendorf tubes as described in ^42^. Briefly, 5μL of a 2x buffer containing 200mM NaCl, 40mM Imidazole, 2mM DTT and 20% glycerol was supplemented with the E-Repeat or control IVT RNA (varying concentrations), PTBP1 (to a final concentration of 60μM) and water to 10μL (final volume). The solution was mixed by pipetting and transferred to one well of an 8 well chamber slide (Ibidi) that had been pre-coated with 3% BSA, washed 3x with RNase-Free water and dried. Droplets were imaged at 10x magnification.

### Electrophoretic Mobility Shift Assays

EMSAs were performed as described in ^78^ except that 40,000cpm of 5’ end labeled RNA were used per condition.

### Quantitative RT-PCR and ActinomycinD treatment

In several experiments we determined the levels of *Xist* by RT-PCR. For experiments with ActinomcyinD treatment, the drug was dissolved in DMSO at 1mg/mL and added to the culture medium to a final concentration of 1μg/mL. For RT-PCR, cells were harvested in 1mL TRIzol (Thermo Fisher), after culture medium removal and PBS wash. RNA was purified over RNAeasy columns (Qiagen). 1μg total RNA was used in a reverse-transcription (RT) reaction with SuperScript III and appropriate strand-specific reverse primer, according to manufacturer’s instructions (ThermoFisher). 1/20^th^ of the RT reaction was used in a quantitative PCR reaction, using either 480 SYBR Green LightCycler PCR mix (Roche), SsoAdvanced Universal SYBR mix (Bio-Rad) or SYBR Green Master Mix (Applied Biosystems) and appropriate primers (see Extended Data Table 1), in triplicate reactions. RT-qPCR experiments were normalized against *Gapdh* or *Rrm2* transcripts.

### Crosslinking and Immunoprecipitation of RNA and high throughput sequencing (iCLiP-Seq) for MATR3, PTBP1 and CELF1

iCLIP experiments were performed as described in ^79^. For CLiP-Seq all washes were conducted for 5 minutes per wash, at 4°C with ice cold buffers. Three confluent 15cm plates of male tetO-*Xist* V6.5 (pSM33) ESCs^24^ were used per immunoprecipitation upon 6 hours of induction of *Xist* expression with 2μg/mL doxycycline, and crosslinking was performed at 100mJ/cm^2^ at 4°C in a Stratalinker 1800 (Stratagene). Crosslinked cells where harvested by scraping in cold 1xPBS and pelleted at 700xg for 2min. Cell pellets were lysed in ice cold lysis buffer [20 mM HEPES-KOH pH 7.5 (Sigma), 150 mM NaCl (Sigma), 0.6% Triton X-100 (Sigma), 0.1% SDS (Sigma), 1 mM EDTA (GIbco), and 0.5 mM DTT (Sigma)] and sonicated in a bioruptor (Diagenode) for 2x 15 minutes (30 sec ON, 30 sec OFF) on high setting at 4°C. Sonicated lysates were cleared by centrifugation at 20,000g, 5min, 4C, supernatants transferred to 15mL falcon tubes and diluted in 5 volumes of buffer containing 20mM HEPES-KOH pH 7.5, 150 mM NaCl, 0.5 mM DTT, 1.25x complete protease inhibitors EDTA-free (Roche), 50 µg/ml yeast tRNA (Life Technologies) and 400U RNAase out (Life Technologies). Samples were briefly mixed and rotated overnight at 4°C. To prepare beads for pulldown, a magnet was used to isolate beads from 200μL of proteinG-dynabead slurry, which were then washed 3x in WB_150_ [20 mM HEPES-KOH pH 7.5, 150 mM NaCl, 0.1% Triton-X100] and incubated overnight at 4°C with 50μg antibody α-MATR3 (Abcam ab151714), α-PTBP1 (Abcam, ab5642) in 700μL WB_150_. Beads were washed 3x in WB_750_ (20mM HEPES-KOH pH 7.5, 750mM NaCl, 0.1% Triton-X100) and 1x with WB_150_ (20mM HEPES-KOH pH 7.5, 150mM NaCl, 0.1% Triton-X100) prior to incubation with lysate. After overnight incubation in lysate, beads were collected at the bottom of the falcon tube with a magnet and the supernatant was removed. Beads were then transferred to a 1.5mL eppendorf tube with 1mL of WB_150_, washed 5x in WB_750_ and 2x in PNK buffer (20 mM HEPES-KOH pH 7.5, 10 mM MgCl2, 0.2% Tween-20). The immunoprecipitated RNA was fragmented in 100 µl of 1x MNase buffer (NEB) containing 5.0 µg of yeast tRNA that was pre-warmed to 37°C in a thermomixer (Eppendorf) set to shake for 15sec ON/15 sec OFF at 750rpm (or minimum speed required to prevent settling of the beads). 50 µl of 1xMNase buffer containing 60 gel units/ml (6 Kunz units/ml) of Micrococcal nuclease (NEB M0247S) were added and incubated for exactly for 5 min. The reaction was stopped with the reaction with 500 µl of EGTA buffer (20 mM HEPES-KOH pH 7.5, 150 mM NaCl, 20 mM EGTA, 0.1% TritonX-100). The beads were then washed 4x in EGTA buffer and 2x in cold PNK buffer. The fragmented RNA was dephosphorylated in 100 µl of 1x FastAP buffer (Fermentas) containing 0.15 U/µl of Fast alkaline phosphatase (Thermo Scientific, EF0651) and 0.2 U/µl of RNaseOUT (LifeTechnologies, 10777-019), incubated in a thermomixer for 90 min at 37 °C, 15 sec shaking/20 sec rest. Beads were washed 4x in WB_750_ and 2x in cold PNK buffer. The dephosphorylated RNA was then ligated to a 3’biotinylated linker RNA in 40μL of buffer containing 1 mM ATP, 25% PEG4000 (Sigma, 202398), 0.5 U/µl T4 RNA ligase1 (NEB M0204S), 0.5 U/μl RNaseOUT, and 6.0 μM L3 linker (Extended Data Table 1). The ligation reaction was incubated in a thermomixer overnight at 16°C, 15 sec ON/4 min OFF at a speed that prevents beads from settling. The next day, beads were washed 4x in WB_150_ and 2x with cold PNK buffer. The RNA was then 5’ end labeled in 24μL PNK wash buffer with 16μL of 1x PNK buffer (NEB) containing 150μCi of gamma P32-ATP, 10U PNK and 1U/μL of RNase OUT. The reaction was incubated in a thermomixer for 20 minutes at 37°C set to shake for 15sec ON/20 sec OFF. The beads were then washed 3x with WB_150_. The immunoprecipitated complexes were eluted off the dynabeads in 50μL of buffer (100mM Tris-HCl pH7.5, 0.6% SDS, 5mM EDTA, 50mM DTT and 50ng/μL yeast tRNA) incubated for 10min at 85°C shaking continuously at 900rpm. The elute was transferred to a new tube and the beads were rinsed with 1200μL of buffer (50mM Tris-HCl pH 7.5, 150mM NaCl, 1.25x complete protease inhibitors (Roche), 50ng/μL yeast tRNA and 0.1% Triton X-100) which was added to the first eluate. The combined eluates were centrifuged for 5min at 4°C at maximum speed and the supernatant transferred to a new tube to prevent carry over of any remaining dynabeads. To prevent IgG heavy chain contamination that co-migrate with many proteins of interest, the biotinylated RNA-protein complexes were bound to monomeric avidin beads. To do this, 10μL packed monomeric avidin agarose beads (Thermo Fisher) were washed 3x with WB_150_. Beads were pelleted after each wash by spinning in a swing bucket rotor at 1000xg, 4°C (use of the swing bucket rotor helps prevent loss of agarose beads). One packed bead volume was mixed with an equal volume of WB_150_ and 15μL of the bead slurry was added to each combined eluate and rotated at 4°C for 4 hours. The beads were then pelleted as above and washed 3x with WB_150_. After the final wash, carefully remove the remaining 5-20μL of supernatant with a p10 pipette. The complexes were eluted off avidin beads by incubation in 30μL of buffer (10mM Tris-HCl pH 7.5, 10% glycerol, 2.2% SDS, 5mM EDTA) at 85°C for 10 minutes in a thermomixer shaking at 900rpm. After centrifugation to pellet the beads, the supernatant was transferred to a new tube and mixed with 5μL of 1xLDS sample buffer (Life Technologies) with 300mM DTT. Samples were incubated at 90C for 10min and then loaded on a pre-run (75v, 10min) NuPAGE Bis-Tris Gel (Life Technologies NP0307) with 1x MOPS running buffer and run for 10-15min at 75v and then 120V until each sample has been satisfactorily separated. The gel was then incubated in transfer buffer (25mM Bis-Tris, 25mM Bicine, 1mM EDTA pH7.2, 20% methanol) for 5 minutes and then transferred onto a protran BA-85 nitrocellulose membrane using a semi-dry transfer apparatus (Biorad 170-3940) for 75min at 400mA (not exceeding 15V). After completion of the transfer, the membrane was briefly washed in milli-Q water, wrapped in plastic film and exposed on a phosophoimager screen for 1 hour. The regions of interest were then excised from the membrane and transferred to an eppendorf tube. The RNA was eluted from the membrane by incubation in 300μL of buffer (100mM Tris-HCl pH 7.5, 50mM NaCl, 10mM EDTA and 2μg/μL proteinase K) for 30 minutes at 55°C in a thermomixer, shaking continuously. 300μL of pre-warmed buffer (100mM Tris-HCl pH 7.5, 50mM NaCl, 10mM EDTA, 7M urea and 2μg/μΛ proteinase K) was then added to the tube and incubated for a further 30min at 55°C. The supernatant was then transferred to a new tube and extracted with an equal volume of phenol:chloroform (5:1, pH 4.5). The separated aqueous phase was precipitated with 0.5μL of Glycblue (Life Technologies), 60μL Sodium Acetate pH 5.4 and 600μL isopropanol overnight at −20C. Next day, the RNA was pelleted by centrifugation at 4°C for 30min at max speed. The pellet was then washed with 1mL 75% EtOH before air drying for 2 minutes, and dissolved in 5.70 μL RNAse free water and left on ice for 5-10 minutes before being reverse transcribed. To do this, 0.5μL of 10mM dNTPs and 0.5μL of 2μM RT primer (Extended Data Table 1) were added to the RNA, mixed by pipetting and denatured for 5 min at 70°C before being snap cooled on ice. The RT primers contain an 11nt Unique Molecule Identifier (UMI) used in sequence analysis (see below). The sample was then equilibrated at 25C in a PCR machine before 3.5μL of RT mix were added (2μL 5x First Strand Buffer, 0.5μL 100mM DTT, 0.5μL 100U/μL Superscript III (Life Technologies) and 0.5μL 40U/μL RNase OUT (Life Technologies)) and incubated for 5 minutes at 25°C and then for 20 minutes at 42C, and then 20 minutes at 48°C. The reverse transcription reaction was then transferred to a new eppendorf tube containing 100μL TE, 11μL 3M Sodium Acetate and 2.5volumes of 100% EtOH. The cDNA was precipitated overnight at −20°C, pelleted and washed as described above, dissolved in 5μL RNase-free water and then mixed with 7.5μL of formamide containing 10mM EDTA, bromophenol blue and xylene cyanol tracking dyes. For size determination, ladder was prepared as follows: 2μL GeneScan 500LIZ size marker (Life Technologies 4322682), 3μL H20, 15μL Formamide containing 10mM EDTA with NO tracking dyes. The samples and ladder were denatured for 5min at 85°C and then loaded on a pre-run 5.5% (19:1 Bis:acrylamide) UREA-PAGE gel (1xTBE, 7.5M UREA) for 20 min at 21V. The gel was then scanned and a gel slice in the range of 70-120nt was excised, chopped into 1mm cubes and the cDNA eluted in 700μL of TE buffer rotating at RT overnight. The next day, the cDNA was precipitated overnight as described above. The washed pellet was then dissolved in 6.7μL of RNase free water and left on ice for 5-10 minutes before being transferred to a PCR tube. Subsequently, the RNA was circularized by addition of 1.5 μl of: 0.8μL of circligase II buffer, 0.4μL 50mM MnCl_2_ and 0.3μL of 100U/μl CicrLigase II ssDNA ligase (Epicentre CL9021K) and incubated in a PCR machine at 60°C for 60 minutes. The circularized cDNA was then digested with BamHI by addition of 30μL of: 4μL 10x FastDigest Buffer, 0.9μL of 10μM cut_oligo and 25.1μL RNase Free water. This mix was incubated for 4 minutes at 95C after which the temperature was decreased 1C each minute until 37°C after which 2μL of FastDigest BamH1 (Thermo Scientific FD0054) was added and incubated at 37°C for a further 30 minutes. The sample was transferred to an eppendorf tube and pelleted as described above. The pelleted DNA was dissolved in 12 μL RNase free water. 2μL were used to prepare a 42μL PCR mix, containing 1x PFU buffer, 0.2mM dNTPs, 0.2μM P3 and P5 solexa primers, and 0.5U PFU polymerase. A negative control containing water instead of cDNA was also prepared. The 42μL reaction was then split into 4×10μL reactions and PCR amplified for 20, 24, 28 and 32 cycles (94°C/3’; 94°C/30s; 63.5°C/15s; 72°C/30s; with final extension at 72°C for 7 minutes). The PCR amplicons were run on a 2% agarose gel in 0.5X TBE/EtBr and the number of cycles required to produce 50-200ng of PCR product from the remaining 10μL of cDNA template was calculated. The PCR reaction was repeated using 10μL remaining ssDNA template, run on a 2% gel as before and the 150-210bp size range was excised and purified using Zymoclean Gel DNA recovery kit (Zymo Research D4007). DNA concentration was determined by qubit using the dsDNA Broad Range assay and prepared for sequencing on an Illumina HiSeq2000 machine using a single end 100bp protocol.

### Enhanced Crosslinking and Immunoprecipitation of RNA and high throughput sequencing (eCLIP) for CELF1

eCLIP experiments against CELF1 (α-CELF1 (ab129115) were performed as described in ^80^ with a few modifications. As with iCLIP, male tet-inducible -*Xist* V6.5 ESCs (pSM33) were induced with 2μg/mL doxycycline for 6 hours prior to crosslinking at 100mJ/cm^2^ at 4°C in a Stratalinker 1800 (Stratagene). Cells were then processed according to the eCLIP protocol for input and immunoprecipitated samples until cDNA was obtained. We then used the iCLIP protocol from the gel-purification of the cDNA through to amplification and purification of the DNA library. For eCLIP samples were sequenced on an Illumina HiSeq2000 machine using the single end 50bp protocol.

### CLIP-seq analysis

CLIP-seq results were mapped using TopHat and processed with the publicly available fastq-tools, fastx-toolkit, Samtools, Bedtools, DeepTools and UCSC scripts ^81 82^. The first 11 bases of each sequenced read correspond to a UMI, composed of a library specific barcode (3 nt) flanked by 4 degenerate nucleotides. The UMI permitted removal of PCR duplicates from the total sequenced reads with the fastq-uniq command-line tool. The Fastx-toolkit was then used to clip 3’ adapter sequences. Sequences shorter than 20 nt were discarded. Reads were then de-multiplexed and mapped to the iGenome mm9 reference genome by Tophat with high stringency settings (no mismatches, reads with multiple genome hits were discarded). Library-depth normalized counts were generated and data was converted to bigWig format to visualize tracks in IGV. Peaks were called using CLIPper ^83^ using the --superlocal option.

### ChIP-seq

3x 15 cm plates of male tetOXist-Δ*Tsix* V6.5 ESCs (∼100 million cells) were used to prepare chromatin for PTBP1 ChiP-seq. Cells were induced for 0 or 20 hours with 2μg/mL doxycycline to induce *Xist* prior to harvesting by trypsinization. Cells were then pelleted by centrifugation at 700g for 5 minutes at RT and resuspended in a total volume of 10mL PBS. The wash step was repeated twice before resuspending in 10 mLs of 1x PBS and transferring to a 50mL falcon tube to which Disuccinimidyl glutarate (DSG) (Pierce) in DMSO was added for crosslinking to a final concentration of 2mM and incubated for 10 minutes at room temperature with gentle mixing under standard laboratory safety practices. Cells were then pelleted and the supernatant was safely disposed of. Cells were re-suspended in 10mL ESC medium and incubated for 10 minutes with 1% formaldehyde (16% methanol free, Pierce) at room temperature with gentle mixing under standard laboratory safety practices. The reaction was quenched by addition of freshly made 0.125M Glycine (Sigma) for 5 minutes at room temperature. Cells were pelleted and supernatant was safely disposed of. Cells were washed twice in 50mL PBS with protease inhibitors (Complete EDTA free, Roche) before being pelleted and flash frozen in liquid nitrogen and stored at −80C. The frozen pellets were processed for ChIP-seq as described in ^69^

### ChIP-seq analysis

Reads were mapped using Bowtie to the iGenome mm9 reference genome. Duplicate reads were removed and length extended to 49 nt. Normalized reads count were generated across 50nt bins. Tracks were visualized in IGV in bigWig format.

### RNA Affinity Purification (RAP)

RAP was performed as described in^24^. For the RAP-seq experiment, we used male T20 ESCs^6^ carrying homing site in the *Hprt* locus on the single X-chromosome as well as a tetracycline-inducible transactivator in the R26 locus. The *Hprt* homing site includes a bidirectional, tetracycline-inducible promoter for expression of a control gene (*EGFP*) and of the *Xist* cDNA transgene introduced later by site-specific recombination, as well as a *lox*P site neighboring the tet-promoter and linked to a truncated neomycin-resistance gene lacking a promoter and translation initiation codon. We integrated two different *Xist* cDNA transgenes into the homing site by electroporation of the respective *Xist* cDNA encoding plasmid and a Cre expression plasmid. The *Xist* transgene plasmid contained a promoter-less *Xist* sequence followed by a poly-adenylation signal and a PGK promoter and translation initiation codon linked to a loxP site. Site-specific recombination of the *lox*P sites in the *Xist* plasmid and the homing site linked the translation initiation codon and *Pgk1* promoter to the *neo* gene, which restored the antibiotic resistance marker. A single copy of the *Xist* cDNA transgene was thus integrated under the control of the inducible promoter. In this study, we employed a ∼14.5kb *Xist* cDNA with either a 4122 nucleotide deletion between BstEii sites within the *Xist* cDNA, deleting the E-repeat and surrounding sequences, or a 1237 nucleotide deletion ending at a similar region with *Xist* and not including the E-repeat, which was generating by deleting internal sequences within the cDNA by SnaBI digestion and re-ligations (see Fig. 2g). Cells were induced with 2μg/mL dox for 6h, prior to fixation for RAP-seq.

### Microscopy

#### Epifluorescence imaging

Immunofluoresence and RNA FISH samples were imaged using a Zeiss AxioImager M1 microscope with a 63x objective and acquired with AxioVision software. Epifluorescence images shown are sections and were analyzed, merged and quantified using ImageJ or Adobe Photoshop.

#### Xist aggregation analysis

To quantify the aggregation of *Xist* clouds, images were taken as Z-stacks and transformed in a Maximum Intensity Projection (MIP) image to detect the entire *Xist* FISH signal in one plane. The background was removed using a rolling ball radius of 50 pixels. *Xist* RNA cloud areas were measured by creating a binary mask over the *Xist* RNA FISH signal for each analyzed *Xist* cloud. Edges of each *Xist* cloud signal were determined by selecting a central pixel and all associated pixels of same intensity value (+/- 5 units). The ImageJ FracLac plugin was then used to calculate the area of a circle encompassing each cloud signal ^84^. The ratio of the *Xist* cloud area over its bounding circle area approximates the compaction of the *Xist* RNA cloud. Significant differences between WT/ΔE ESC or siRNA treated samples were tested with the non-parametric 2 sample Kolmogorov-Smirnov (K-S) test.

#### 3D-Structured Illumination Microscopy (3D-SIM)

3D-SIM super-resolution imaging was performed on a DeltaVision OMX V3 system (Applied Precision, GE Healthcare) equipped with a 100 Å∼/1.40 NA Plan Apo oil immersion objective (Olympus, Tokyo, Japan), Cascade II:512 EMCCD cameras (Photometrics, Tucson, AZ, USA) and 405, 488 and 593 nm diode lasers. Image stacks were acquired with a z-distance of 125 nm and with 15 raw images per plane (five phases, three angles). The raw data were computationally reconstructed with the soft-WoRx 6.0 software package (Applied Precision) using a wiener filter set at 0.002 and channel-specifically measured optical transfer functions (OTFs) using immersion oil with different refractive indices (RIs) as described in ^85^. Images from the different channels were registered using alignment parameters obtained from calibration measurements with 0.2µm diameter TetraSpeck beads (Invitrogen) as described in ^75^.

#### Improved Confocal microscopy

Improved confocal laser scanning microscopy was performed on a LSM880 platform equipped with 100x/1.46NA or 63x/1.4 NA plan Apochromat oil objectives and 405/488 diode and 594 Helium-Neon lasers using the Airyscan detector (Carl Zeiss Microscopy, Thornwood, NY). An appropriate magnification was used in order to collect image stacks from a region that encompassed the nucleus of interest thereby optimizing imaging time and reducing photobleaching. The pixel size and z-optical sectioning were set to meet the Nyquist sampling criterion in each case. Airyscan raw data were linearly reconstructed using the ZEN 2.3 software.

### Quantitative image analysis

All image analysis steps were performed on Fiji/ImageJ ^86, 87^.

#### CELF-1intensity plot profiles

Airyscan image stacks were imported into ImageJ and converted to 16-bit composites. The 3D-stacks were reduced into 2D images and 2µm intensity line plots were used to extract the intensity profiles over the Xi enriched signal in the CELF1 channel. The same line-plot was used in a random nucleoplasmic region to select for the average nuclear CELF-1 intensities. The ratio of the top 10% intensities of the signals were plotted after diving over the nucleoplasmic signal.

#### Xi DAPI intensities quantification

Wide-field image stacks were generated from 3D-SIM raw data of H3K27me3 and DAPI stained cells by average projection of five consecutive phase-shifted images from each plane of the first angle and subjected to an iterative 3D deconvolution using soft-WoRx 6.0 software. The reconstructed image stacks were imported to ImageJ and converted to 8-bit files. In order to measure the Xi underlying DAPI intensity, binary masks from the H3K27me3 channel were created to define the Xi territory of day 7 WT and ΔE ESC nuclei. A threshold was carefully applied selecting the boarder of H3K27me3 enriched region that demarcates the Xi territory. Subsequently the grey values of the corresponding masked region in the DAPI channel were extracted and plotted.

#### Segmentation of Xist RNA foci from 3D-SIM datasets

The 32-bit reconstructed 3D-SIM image stacks were imported into ImageJ were grey values were shifted to the positive range and converted to 16-bit composites after subtracting the mode grey value to remove background noise. Segmentation of *Xist* RNA foci was performed by using the TANGO plugin ^88^ on ImageJ according to the pipelines described in ^75^. In brief, nuclear masks were created by using the nucleus processing chain. *Xist* foci were segments by first pre-filtering with a TopHat filter with a radius of 1 pixel in all three dimensions (xyz), followed by a Laplace of Gaussian filter with a radius of 1-pixel (x,y,z). Segmentation of foci was performed using the spot detector 3D with Otsu auto-thresholding. Segmented objects were post-filtered with a size and edge filter of 5 pixels per spot and an SNR above 2.

#### Amira reconstructions

3D-reconstructions were performed using Amira 2.3 (Mercury Computer Systems, Chelmsford, Massachusetts, United States). Image stacks were imported into Amira as separate channels. *Xist* FISH or antibody stainings were reconstructed as surface renderings while DAPI was reconstructed as volume rendering using the Volren module that allows visualization of intensity in color maps.

### Genomics data

Genomics data (CLIP-seq, ChIP-seq and RAP-seq) will be made accessible with publication on the GEO database.

#### Technical Note 1

The *Xist* and MS2 DNA FISH probes were synthesized using Klenow Exo-polymerase primed with random hexamers from DNA templates. Specifically, for the *Xist* probe, we used a plasmid encoding the entire 17.9kb *Xist* cDNA as template. The MS2 probe was synthesized from a PCR product that amplified the 1.1kb MS2 repeat sequence from genomic DNA of targeted F1 2-1 ESCs. See Fig. 2a for illustration of the sequences covered by the two FISH probes. The difference in sequence length available for the probes to bind to *Xist* transcripts in cells meant that the *Xist* probe gave more intensely fluorescent FISH signals than the MS2 probe. To assign the *cas* or *129* allele-of-origin for the *Xist* cloud, we first looked for *Xist* probe signal then asked whether we could detect *<u>any</u>* co-localizing MS2 probe signal. Thus, while we show that 50-70% of the cells with a detectable *Xist* cloud at late stages of initiation (day 4 onward) still show MS2 co-localizing signal cloud formation in ΔE cells, we wish to emphasize that almost none of these clouds presented as morphologically intact and our results, as presented, under-represent the penetrance of the dispersal phenotype observe for ΔE-*Xist* transcript at later stages of XCI initiation.

---

**Extended Data Figure 1:**
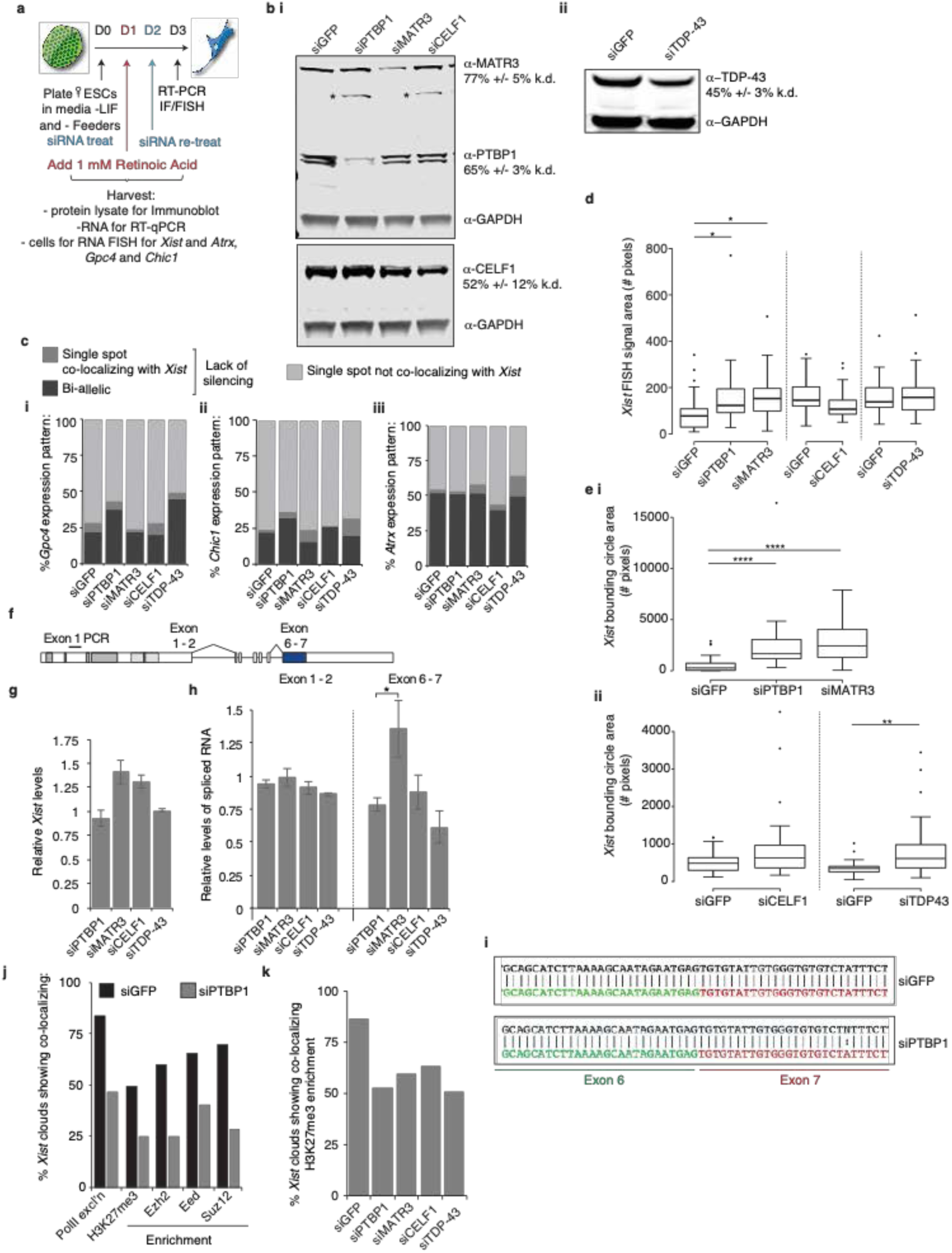
Depletion of PTBP1, MATR3, TDP-43 or CELF1 does not strongly affect X-linked gene silencing or Xist processing. **a,** Overview of siRNA approach used to deplete *Xist*-interacting RBPs in female differentiating mouse ESCs and to assess XCI. **b i,** Representative immunoblot showing the level of siRNA-mediated knockdown of PTBP1, MATR3, and CELF1 relative to siGFP treatment in cells differentiated for 3 days. The percentage depletion normalized to GAPDH is shown on the right. Error represents the SEM from 3 independent experiments. **ii,** As in (i), except for TDP-43. **c i,** Histogram showing the nascent expression pattern of X-linked gene *Gpc4* after 3 days of differentiation and knockdown of the indicated factor. 50 cells with *Xist* clouds were counted per sample from one experiment. **ii,** Same as (i) except showing data for the X-linked gene *Chic1*. **iii,** Same as (i) except showing data for the X-linked gene *Atrx*. **d,** Box plots showing the distribution of the area (mask) covered by *Xist* RNA FISH signal upon knockdown of PTBP1, MATR3, CELF1 or TDP-43, used to calculate the *Xist* aggregation scores displayed in Figure 1b. (p-value: * < 0.05, 2-sample KS test). 25-36 clouds were measured per sample, from one experiment. **e i,** Box plots showing the distribution of the area of the bounding circle encompassing *Xist* FISH signal/mask upon knockdown of PTBP1 or MATR3 used to calculate the *Xist* aggregation scores displayed in Figure 1b. **ii,** Same as (i) except for knockdown of CELF1 or TDP-43. (p-value: * < 0.05, **** < 0.00005, 2-sample KS test). 25-36 clouds were measured per sample, from one experiment. **f,** Diagram of *Xist* showing the splicing events assessed upon knockdown of PTBP1, MATR3, TDP-43 or CELF1, and amplicon used to assess *Xist* abundance (exon 1 PCR). The results are shown in (i) and (j). **g,** Histogram showing the relative abundance of total *Xist* transcripts upon knockdown of indicated RBPs. All samples are normalized to siGFP control treatment samples and *Snrnp27* RNA (encodes a U6 snRNP factor). Samples were assessed in triplicate from 3 independent depletions. Error bars represent the SEM. **h,** Histogram showing the relative level of *Xist* exon 1-2 and exon 6-7 splicing upon knockdown of indicated factors, normalized as in (j). Samples were assessed in triplicate from three independent depletions. Error bars represent the SEM (p-value: * < 0.05, 2-tailed students t-test). **i,** Snapshot of aligned sequences derived from the expected sequence upon splicing of exon 6 (green) - exon 7 (red) (colored, lower sequence), and the observed sequence upon siRNA treatment targeting GFP or PTBP1 after 72h in differentiating female ESCs (black, upper sequence). Two independent experiments were tested for splicing with identical results showing correct exon 6 - 7 ligation. **j,** Histogram showing the percentage of *Xist-*positive cells showing co-localizing exclusion of RNA-pol II or enrichment of PRC2 components EZH2, EED, SUZ12 and the histone modification H3K27me3 on the Xi, in female ESCs differentiated for 3 days and treated with siRNAs against GFP (control) or PTBP1. 54-64 cells were counted per sample from one experiment. **k,** Histogram showing the percentage of *Xist-*positive cells with H3K27me3 Xi-enrichment in day 3 differentiated female ESCs treated with siRNAs against GFP (control), PTBP1, MATR3, TDP-43 or CELF1. 109-146 cells were counted per sample, from one experiment independent of that shown for (f).

**Extended Data Figure 2:**
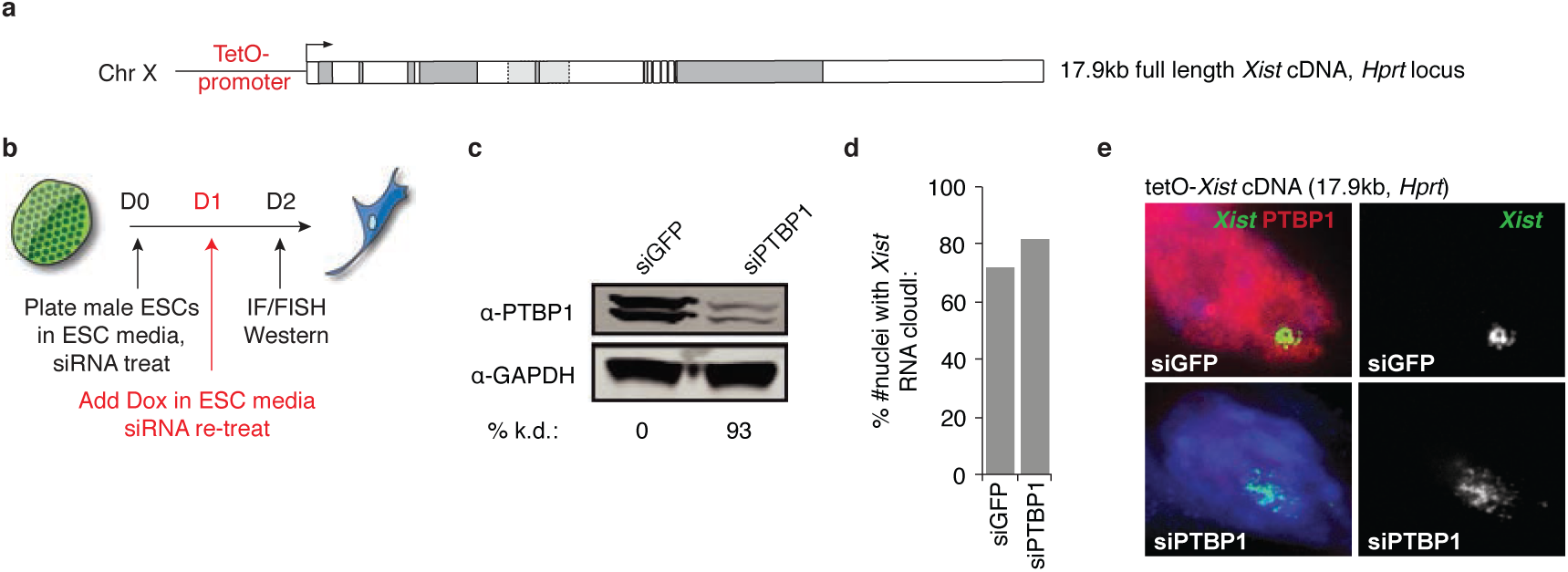
*Xist* dispersal upon PTBP1 knockdown is independent of splicing. **a,** Diagram of a tet-inducible full length (17.9kb) *Xist* cDNA transgene inserted into the *Hprt* locus on the X-chromosome in male ESCs. **b,** Approach used to deplete PTBP1 in the cells described in (a). **c,** Immunoblot and quantification showing level of PTBP1 knockdown in the experiment diagrammed in (b). **d,** Histogram showing the percentage of cells with an *Xist* RNA cloud upon knockdown of PTBP1 and 24 hours of dox treatment in the experiment as described in (b). 82-90 cells were counted per sample in one experiment. **e,** Representative images of *Xist,* detected by RNA FISH, after being treated with siRNAs against GFP or PTBP1 and dox-addition, as outlined in (b). Note the dispersal of *Xist* upon PTBP1 knockdown occurs in the absence of splicing regulation, as *Xist* is expressed from a cDNA construct in this experiment.

**Extended Data Figure 3:**
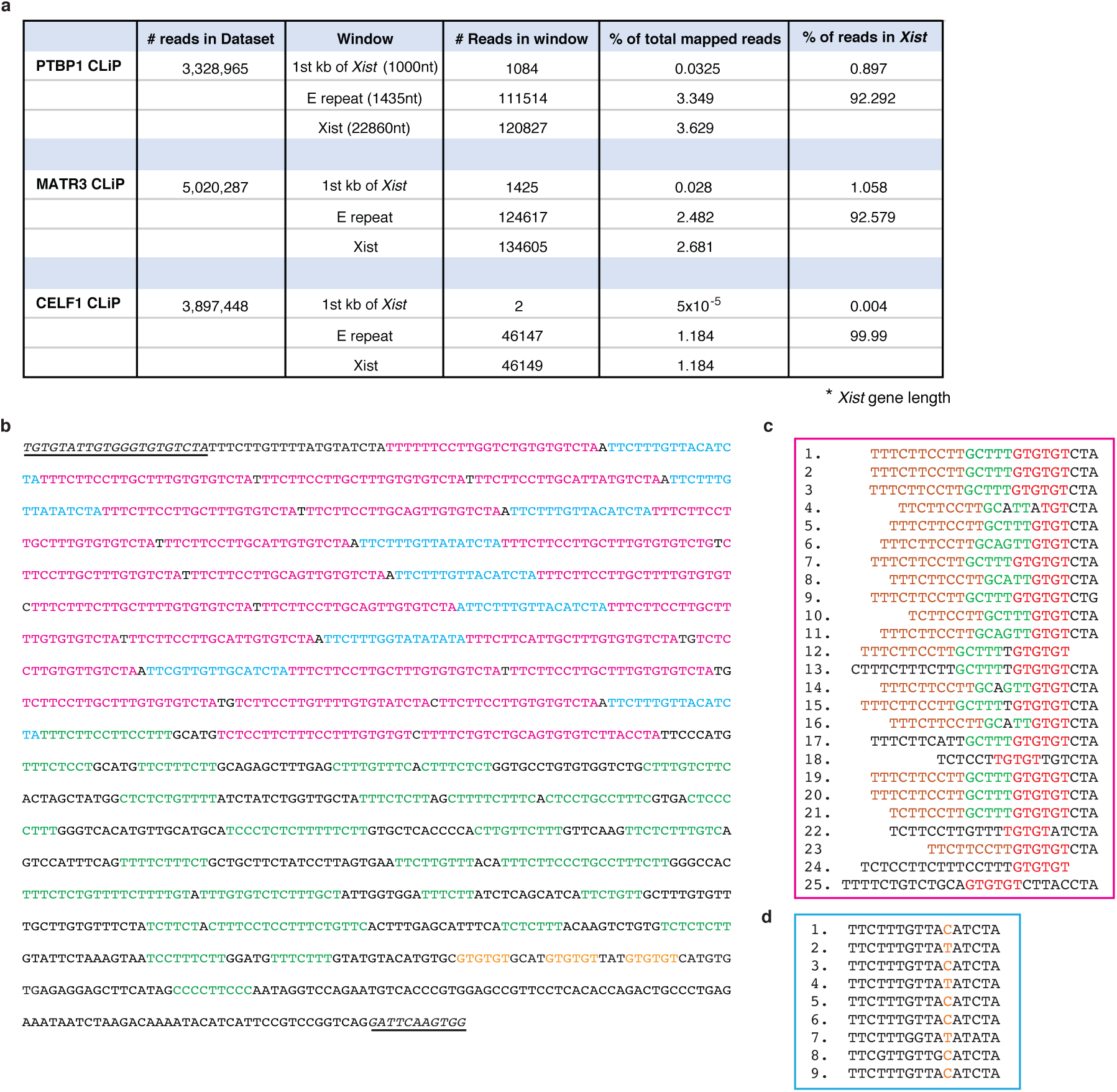
A tandem array of 20-25nt C/U/G-rich elements comprises the *Xist* E-repeat. **a,** Table showing mapping statistics of PTBP1, MATR3 and CELF1 CLIP-seq data for male ESCs with a tet-inducible *Xist* allele, induced for 6 hours, for reads mapping across the *Xist* locus, the E-repeat of *Xist,* and the 1^st^ kb of *Xist* (including the A-repeat) as a percentage of total mapped reads or of the reads located within *Xist*. Note that *Xist* is overexpressed in this experiment, which influences the total number of reads mapping to the locus. **b,** The first 1500nt of the terminal exon 7 of *Xist* are shown. The sequence remaining after splicing of the *Xist* ΔE transcript is underlined (see Extended Data Figure 6). The C/U/G tandem repeats within the 5’ half of the E-repeat are indicated (pink – full repeats, blue-truncated repeats) as are the CU-tracts (shown as CT tracks in the DNA sequence, green) in the 3’ half of the sequence. Potential TDP-43 sites are shown in orange at the 3’ terminal end of the E-repeat. **c,** Alignment of the 25 full C/U/G-tandem repeats (pink) in (b). Brown, green, red and black text indicate similar nucleotide runs within each tandem repeat. Brown C/U tracts encode putative PTBP1 and/or MATR3 binding sites. Red G/U tracts encode putative CELF1 and/or TDP-43 binding sites. **d,** Alignment of the 9 truncated C/U/G-tandem repeats (blue) in (b)). Orange nucleotides are variable within each truncated repeat unit.

**Extended Data Figure 4:**
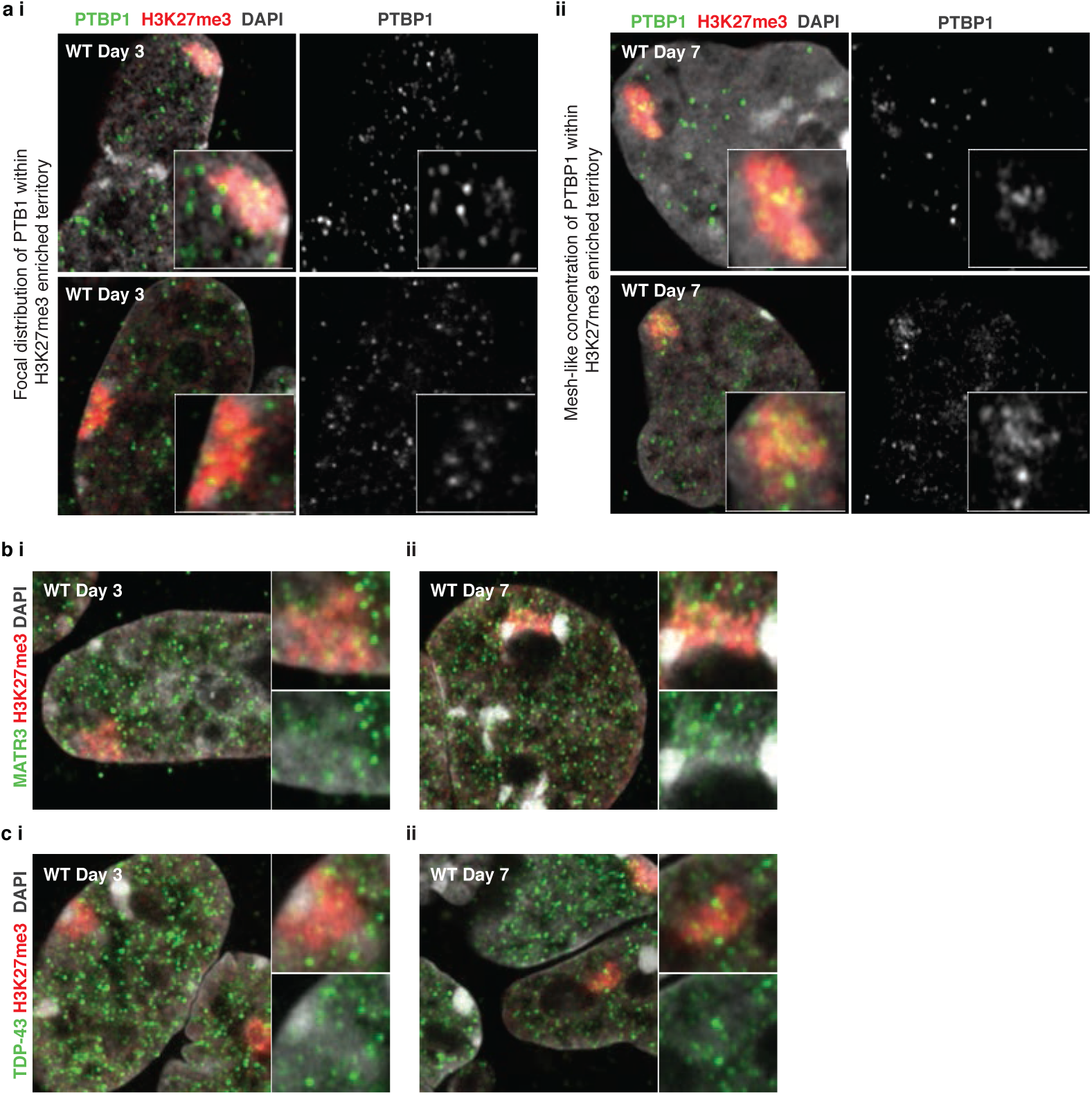
PTBP1 concentrates in a unique pattern within the *Xist-*coated X-chromosome territory at day 7 of XCI initiation. **a i,** Left: Two independent micrographs of confocal-airyscan optical sections through nuclei of WT female ESCs at day 3 of differentiation immunostained for PTBP1 (green) and H3K27me3 (red, detecting the Xi) and counterstained with DAPI (grey). Inset: enlargement of the Xi-territory, showing a punctate, focal pattern of PTBP1 within the Xi. Right: same as left, except showing only the PTBP1 signal in grayscale. **ii,** Same as in (i), except showing WT female ESCs differentiated for 7 days, showing a mesh-like PTBP1 concentration within the Xi territory that is distinct from that observed in the nucleoplasm of these cells or from the pattern within the day 3 Xi. **b i,** Confocal-airyscan optical sections through nuclei of WT female ESCs at day 3 of differentiation immunostained for MATR3 (green) and H3K27me3 (red) and counterstained with DAPI (gray). Right: enlargement of the Xi territory showing MATR3 and H3K27me3 signals (upper) or only MATR3 signal (lower) with DAPI counterstaining (gray). **ii,** Same as in (i), except showing cells differentiated for 7 days. **c i,** Same as in (b-i) except showing immunostaining for TDP-43 at day 3 of differentiation. **ii,** Same as in (b-ii) except showing immunostaining for TDP-43 at day 7 of differentiation.

**Extended Data Figure 5:**
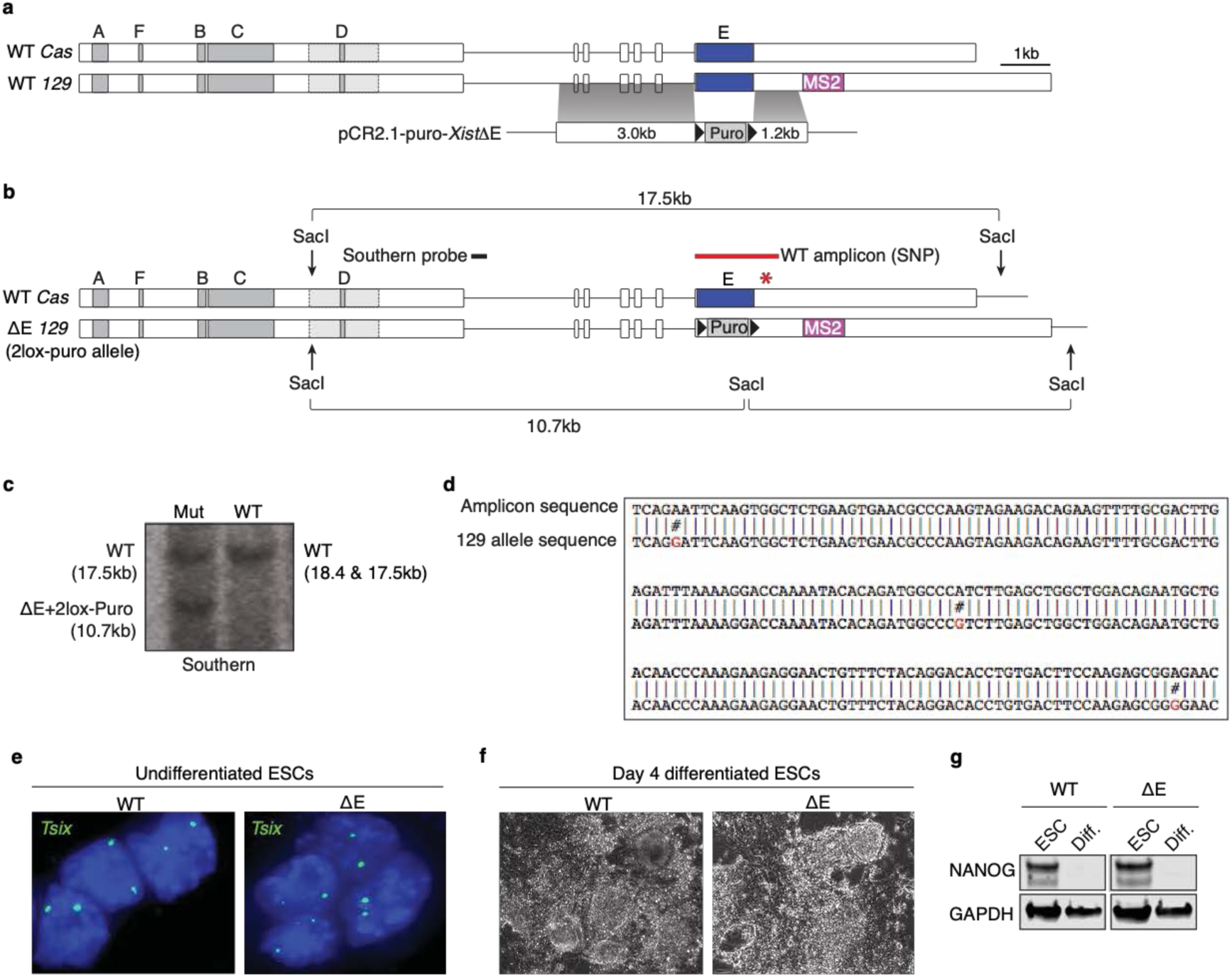
ΔE cells undergo differentiation similar to WT cells. **a,** Scale diagram of the *Xist* gene annotated with repeat regions A-F (see Figure 2a for complete description). Also shown is the targeting strategy used to delete the *Xist* E-repeat region in F1 2-1 MS2 female ESCs with the targeting plasmid pCR2.1-Puro-*Xist*ΔE. **b,** Diagram of the southern blot strategy with a 5’ external probe, used to identify deletion clones. This strategy did not distinguish between targeting of the E-repeat on the 129 or cas allele of *Xist*. Location of southern probe is shown, as is the location of the PCR amplicon (red) used to identify which allele was deleted (see (d)). Asterisk denotes location of SNPs used for defining the allelic origin of the deleted allele (see (d)). **c,** Southern blot as described in (b) on targeted female ESCs with a loxP-flanked puromycin cassette in place of the E-repeat on one *Xist* allele. The puromycin cassette was subsequently deleted by Cre-recombinase transfection into the mutant line and confirmed by PCR after sub-cloning. **d,** Sequencing analysis of the long amplicon (shown in b). SNPs conforming to the *129* allele are shown in red and are not detected in the amplicon, confirming that the E-repeat remains present on the *Cas* allele and is deleted on the *129* allele. **e,** *Tsix* RNA FISH (green) on undifferentiated WT and ΔE cells confirming the presence of two *Tsix* nascent transcription units, used as a proxy to confirm that the targeted ΔE cells maintain two X-chromosomes. DAPI stained nuclei are shown in blue. **f,** Brightfield images of WT and ΔE cells at day 4 of differentiation, showing that the cells do not appear morphologically different upon induction of differentiation. **g,** Immunoblot of lysates from WT and ΔE cells at 2 days of differentiation showing early and equal loss of Nanog expression, a pluripotency gene quickly down-regulated with the induction of differentiation.

**Extended Data Figure 6:**
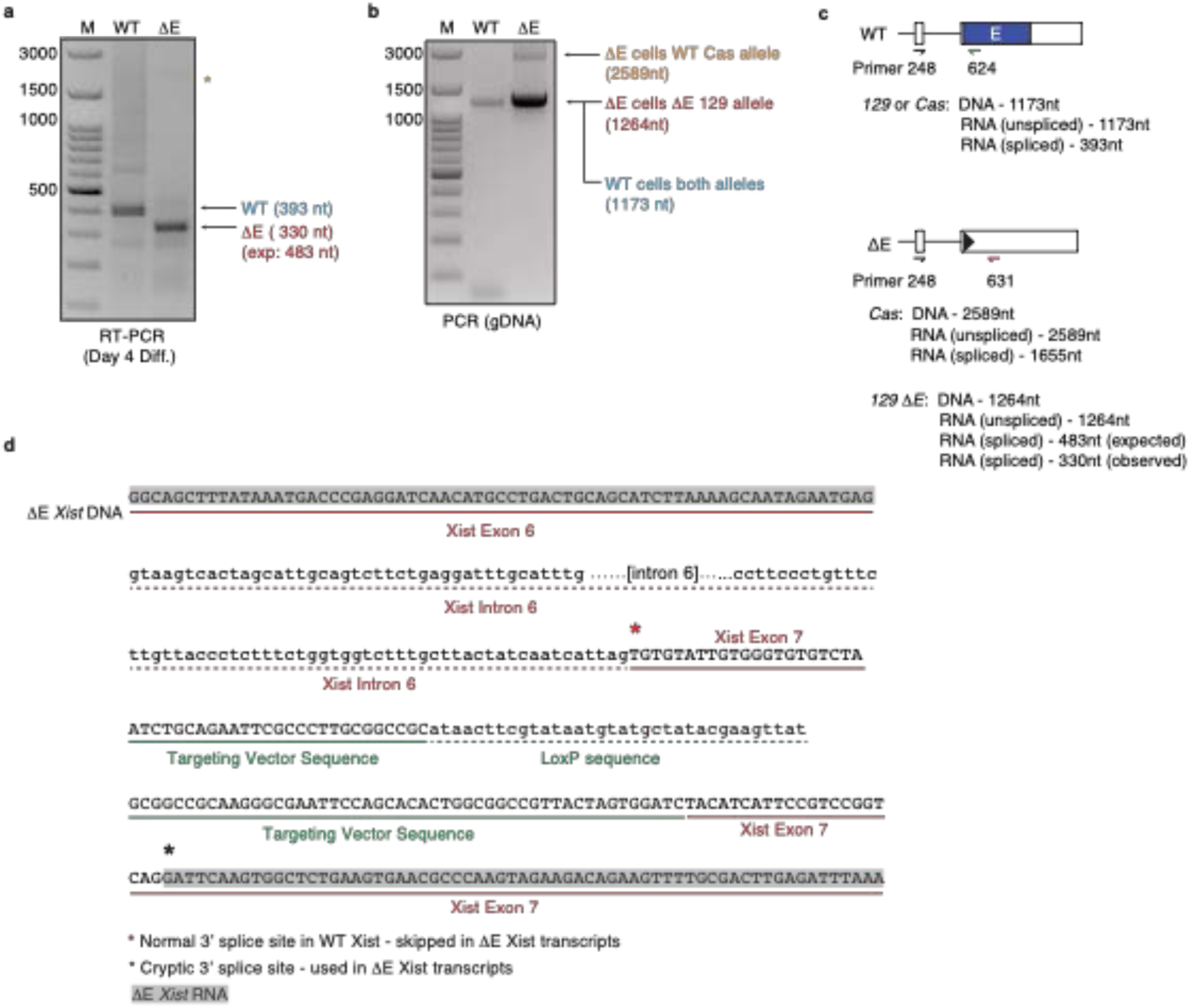
Splicing of intron 6 proceeds in the absence of the E-repeat. **a,** RT-PCR amplicons generated from RNA isolated from day 4 differentiated WT (with primers APJ248/624) or ΔE (with primers APJ248/631) cells across exon 6 – exon 7 to confirm splicing of intron 6. (see (c) for primer locations). We found that that the PCR amplicon was shorter than expected for the ΔE transcript. Sequencing across the modified exon 6 - 7 junction revealed the use a cryptic 3’ splice site downstream of the LoxP site which extended the E repeat deletion within the *Xist* transcript (but not the *Xist* genomic DNA) (as designed) by 42nt. This caused the LoxP site and associated vector sequences to be excluded from the mature ΔE *Xist* transcripts, such that the 3’ terminus of exon 6 splices to nucleotide 1479 of exon 7 to form a scar-less junction (see (d)). Orange asterisk indicates a faint band at the expected location for the amplicon generated from the WT *Cas* allele in ΔE cells. **b,** PCR amplicons from WT or ΔE genomic DNA using the same primers as in (a) showing that the intron 6-containing products can be amplified and thus, non-detection of intron 6-containing *Xist* transcripts is due to splicing of ΔE *Xist* and not amplification problems. **c,** Schematic outlining primers used to assess *Xist* gDNA and RNA in (a) and (b) across the exon 6 / intron 6 and exon 7 region in WT and ΔE cells that have been differentiated for 4 days. **d,** Diagram of sequenced genomic and cDNA amplicons (using APJ248/631) confirming correct integration of the targeting plasmid and cre recombinase-mediated removal of the puromycin cassette as well as the sequence of the spliced *Xist*ΔE transcript from exon 6 to exon 7. Red asterisk denotes wild-type 3’ splice site on exon 7. Black asterisk denotes splice site used in in ΔE transcripts.

**Extended Data Figure 7:**
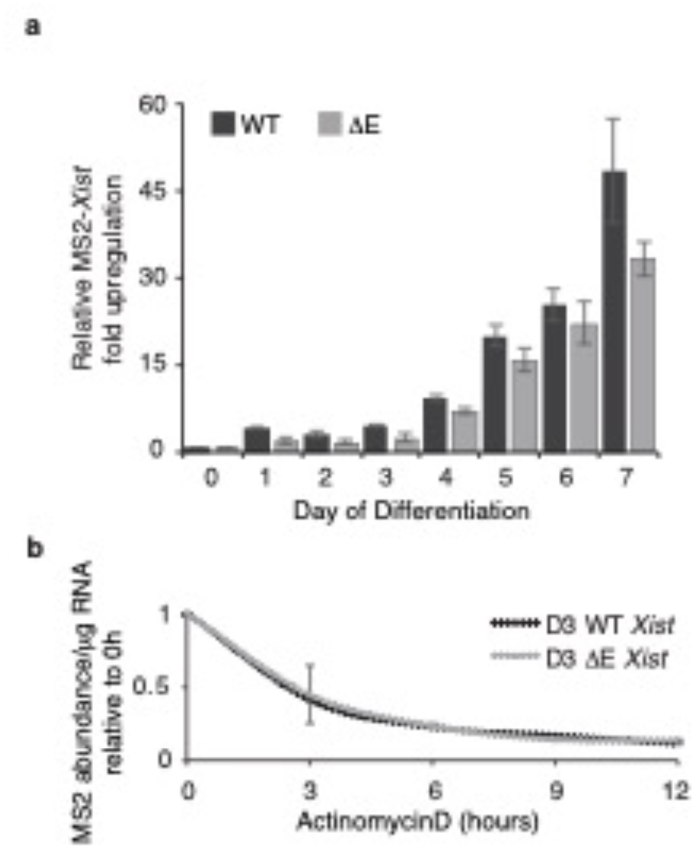
Deletion of the E-repeat does not affect *Xist* abundance or stability. **a,** Histogram showing RT-qPCR quantification of the fold upregulation of MS2+*Xist^129^* RNA in WT and ΔE cells over 7 days of differentiation, normalized relative to undifferentiated samples and against an internal control (*Rrm2*). Error bars represent the standard error of the mean for three independent samples measured in triplicate. Differences were not significant by 2-tailed students t-test. **b,** Line plot showing RT-qPCR measurements of *Xist* RNA half-life in in WT and ΔE cells treated for 3, 6, 9 or 12h with ActinomycinD at day 3 of differentiation. *Xist* RNA amounts were calculated as transcript copy number per microgram of total RNA. Note that ΔE *Xist* RNA displays a similar stability as WT *Xist* at day 3 of differentiation, which is before the major XCI phenotype in ΔE cells manifests, indicating that the deletion of the E-repeat does not make *Xist* inherently less stable. Samples measured in triplicate from 3 independent samples. Error bars indicate the SEM.

**Extended Data Figure 8:**
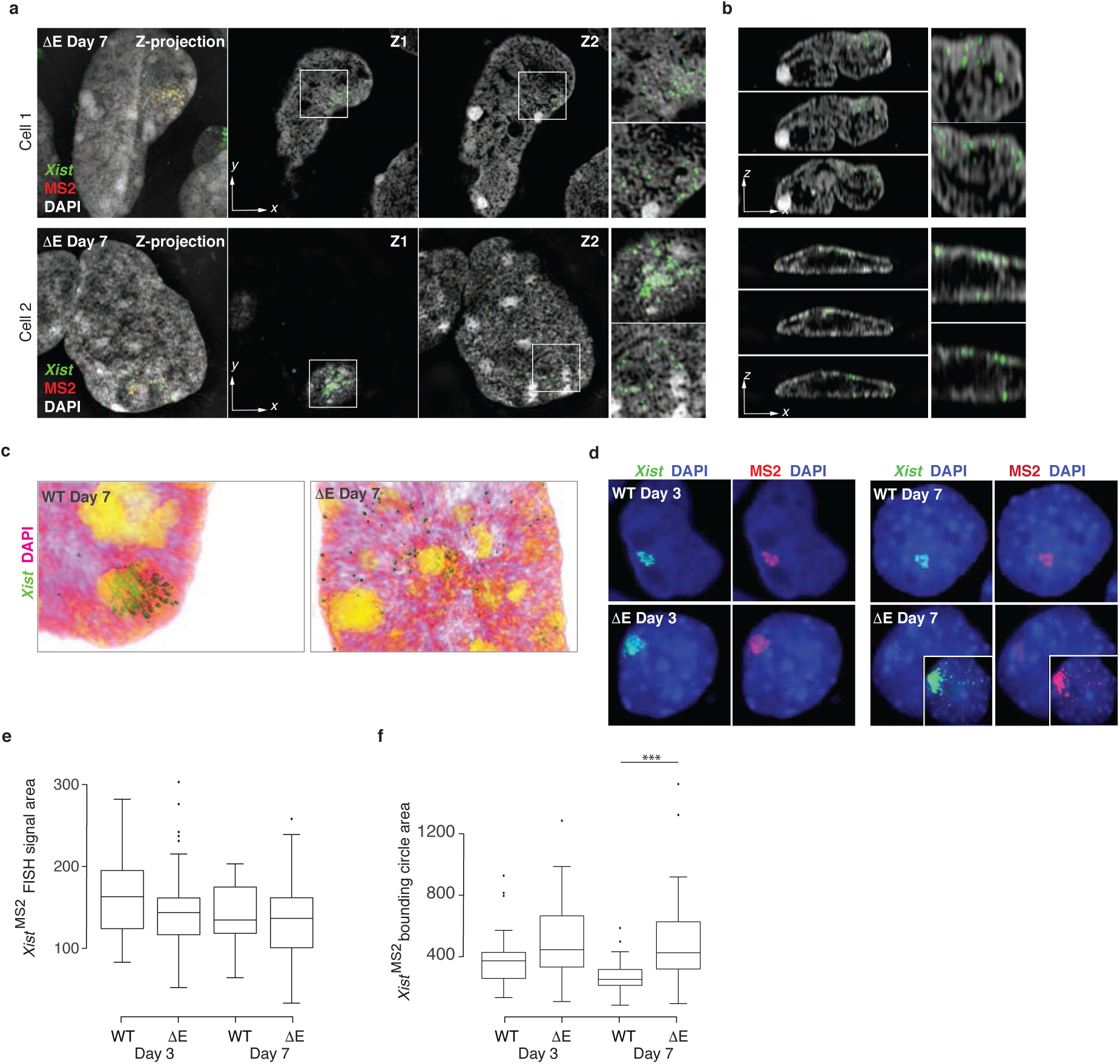
ΔE *Xist* transcripts migrate to the nuclear lamina at late stages of XCI-initiation. **a,** Left: MIP of 3D-SIM optical sections across a nucleus in ΔE cells at day 7 of differentiation hybridized with *Xist* and MS2 probes. *Xist* is shown in green, MS2 in red and DAPI in gray. Next images to the right: Two different lateral sections (Z1, Z2) through the same nucleus. Far Right: Magnifications of marked regions corresponding to the *Xist*-MS2-coated X-chromosome from each plane (Z1, top; Z2, bottom). Two independent nuclei are shown in addition to the cell shown in Figure 2h. Scale bar: 5µm. **b,** Left top to bottom: consecutive axial cross-sections of the nuclei shown in (a). Right: Magnification of regions with the *Xist* RNA FISH signal in the top and bottom left panels. Scale bar: 5µm. Note the aberrant localization of *Xist* signal at the nuclear lamina in ΔE cells. **c,** 3D Amira reconstructions of the *Xist*-MS2-coated/expressing X-chromosome and surrounding nuclear region in WT and ΔE cells shown in Figure 2h *Xist* is shown in green, DAPI with intensity rendering (white/low to orange/high). **d,** Representative conventional epifluorescence images of RNA FISH against *Xist* (green) and MS2 (red) in day 3 and 7 differentiating WT and ΔE cells. Chromatin staining with DAPI is shown in blue. Inset: Enhanced image of the *Xist*/MS2 RNA FISH signal in ΔE cells at Day 7. Epifluorescent images are shown for comparison to super-resolution images shown in Figure 2e/h and (a). **e,** Box plot showing the distribution of the area (in pixels) covered by *Xist* RNA FISH signal at day 3 and 7 of differentiation in WT and ΔE cells, for MS2+*Xist^129^* clouds, used for the calculation of the *Xist* aggregation score in Figure 2f. 30-34 cells were counted per sample from one experiment. 2-sample KS test revealed no significant differences between samples. **f,** Box plot showing the distribution of the area (in pixels) of the bounding circle encompassing *Xist* RNA FISH signal at day 3 and 7 of differentiation in WT and ΔE cells, for MS2+*Xist^129^* clouds, used for the calculation of the *Xist* aggregation score in Figure 2f. 30-34 cells were counted per sample from one experiment (p-value: **** < 0.00005, 2-sample KS test).

**Extended Data Figure 9:**
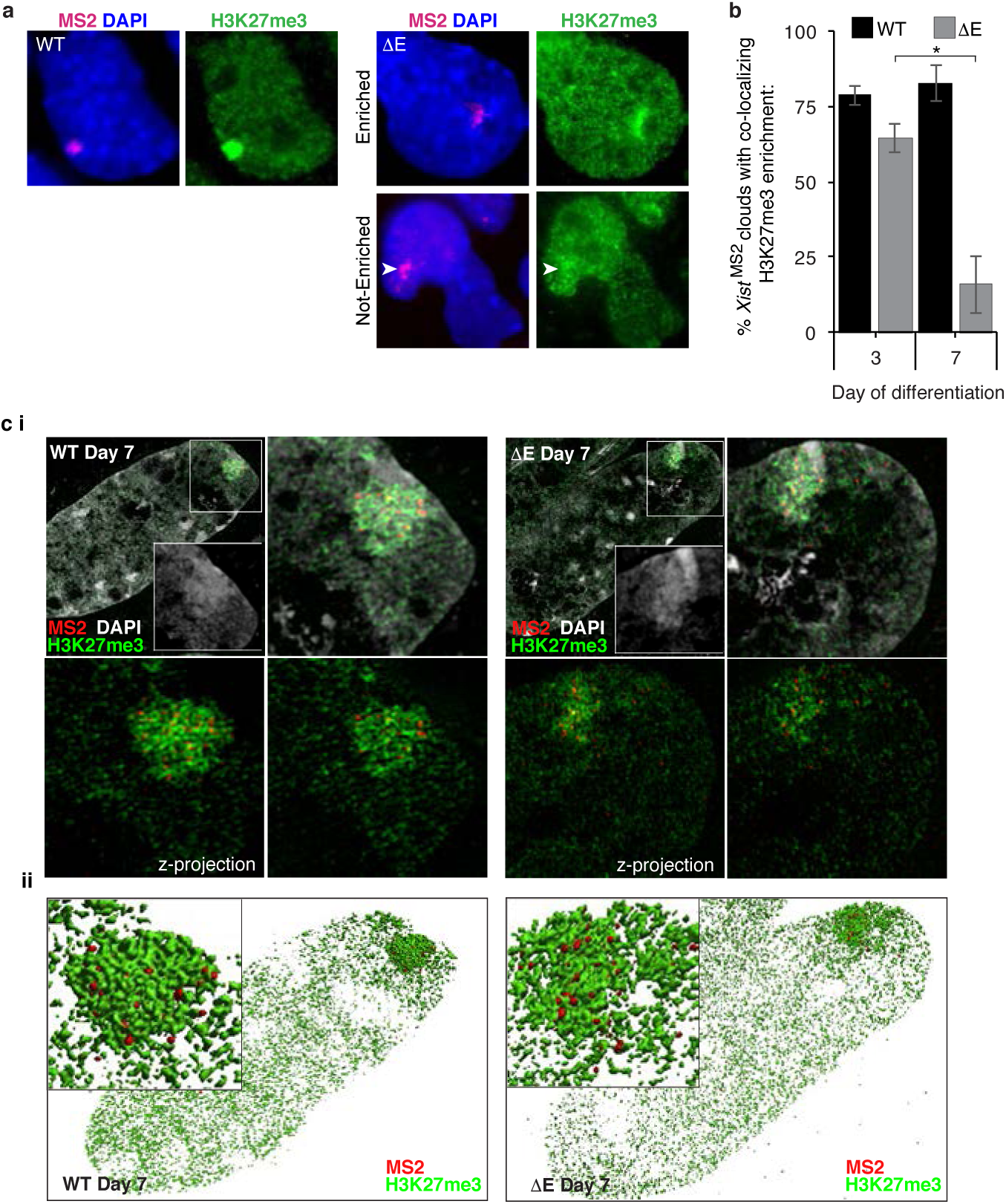
At later stages of XCI initiation, the H3K27me3 accumulation on the ΔE *Xist-*coated X-chromosome is less compact than in WT cells. **a,** Epifluorescence images of WT and ΔE cells at day 7 of differentiation, stained for H3K27me3 (green), a well-established heterochromatin marker of the Xi, and probed for MS2 by RNA FISH (red). In WT cells, nearly all cells displayed H3K27me3 Xi-enrichment, whereas ΔE cells often lacked this Xi-enrichment when *Xist*-MS2 was expressed. Arrowheads indicate expected position of MS2 co-localizing H3K27me3 immunofluorescent signal. **b,** Histogram showing the number of cells with an MS2+ *Xist* RNA FISH cloud that also displayed a co-localizing accumulation of H3K27me3 at days 3 and 7 of differentiation in WT and ΔE cells. 60 cells with *Xist*^MS2^ signal were counted per sample, from 3 coverslips over 2 independent experiments. (p-value: * < 0.05, 2-sample students t-test). To discern enrichment, we first screened cells using the H3K27me3 channel to identify potential H3K27me3 Xi-enrichment, which was then confirmed by checking for a co-localizing MS2 FISH signal. An inability to confidently discern the Xi prior to confirmation with MS2 co-localization was scored as not enriched. **c i,** Top left: Lateral 3D-SIM optical sections through the nucleus of day 7 differentiating WT (left) and ΔE (right) cells stained for H3K27me3 (green) and hybridized with MS2 probes (red). Chromatin counterstaining with DAPI is shown in grey. Inset: DAPI staining of H3K27me3-enriched region corresponding to the *Xist*-coated X-chromosome. Right: Magnification of inset area showing the H3K27me3, MS2 and DAPI staining (top right) or without DAPI (bottom right). Bottom left: Z-stack MIP of the same cell shown in the bottom right, allowing a comparison between z-stack projection or single optical section of the magnified inset area showing H3me3K27 (green) and MS2 (red) signals. **ii,** 3D Amira reconstruction of the data shown in (i). H3K27me3 shown in green, MS2 in red. Note less dense packaging of super-resolved H3K27me3 segments around dispersed *Xist* MS2 signal in the ΔE nucleus compared to WT, suggestive of chromatin de-compaction as shown in Extended Data Figure 10 with DAPI staining.

**Extended Data Figure 10:**
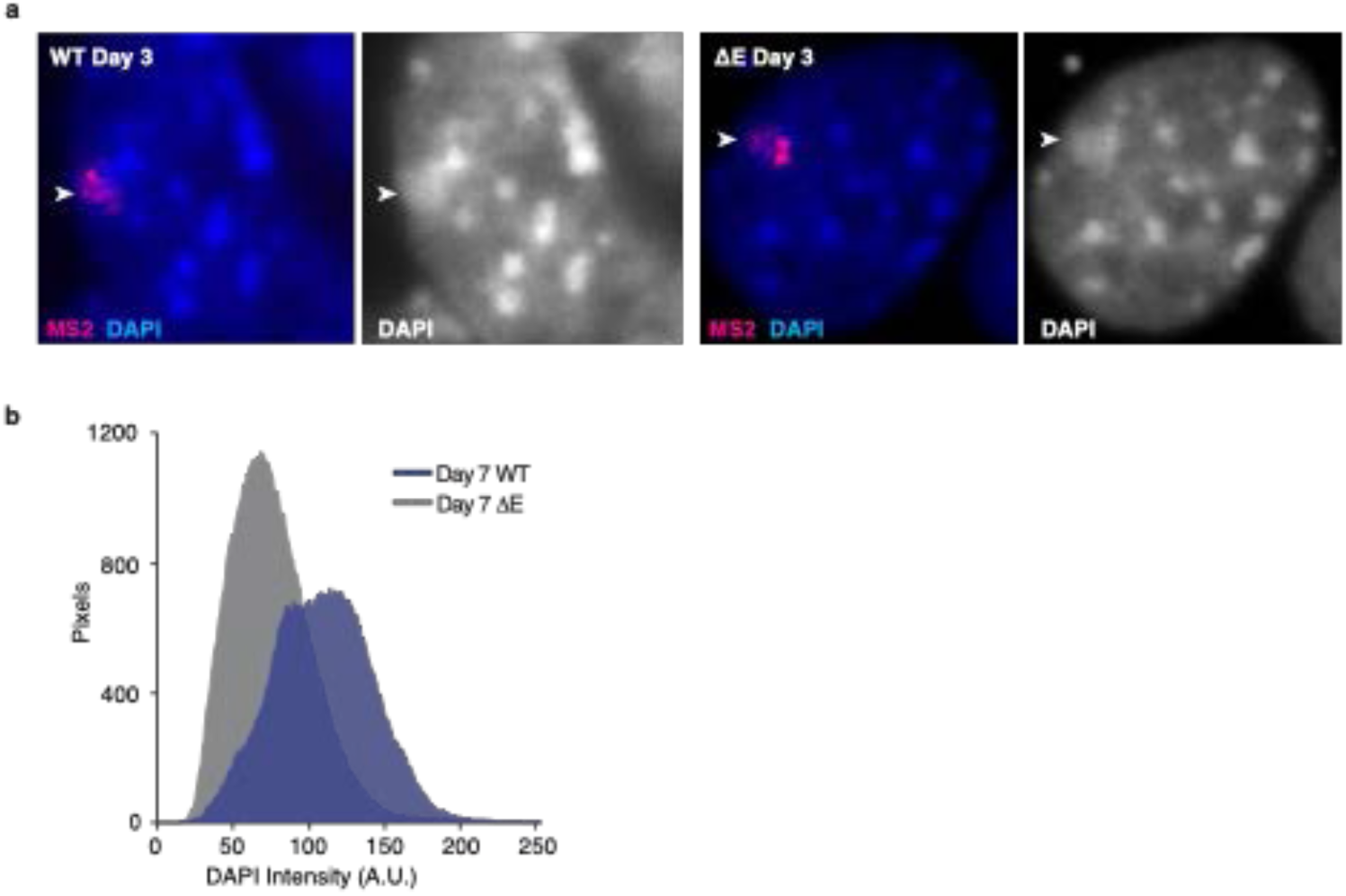
DAPI staining intensity of the ΔE *Xist-*coated X-chromosome decreases compared to WT cells at day 7 of differentiation and XCI-initiation. **a,** Representative epifluorescence images of day 3 differentiated WT and ΔE cells, probed for the MS2 RNA tag by RNA FISH (red). DNA was stained with DAPI (displayed in blue (left) and in grey (right)). Arrowheads point to location of the *Xist* cloud, highlighting the DAPI-intense staining for WT and ΔE *Xist*-coated X-chromosomes at day 3 of differentiation. **b,** Line plot showing DAPI fluorescence intensity within the MS2-*Xist*-expressing X-chromosome masked by H3K27me3 enrichment in WT (blue) and ΔE (gray) cells, differentiated for 7 days. 10 cells were analyzed per cell line from one experiment. These data indicate that the loss of ΔE *Xist* late in differentiation is accompanied by the loss of intense DAPI staining, a hallmark of the Xi.

**Extended Data Figure 11:**
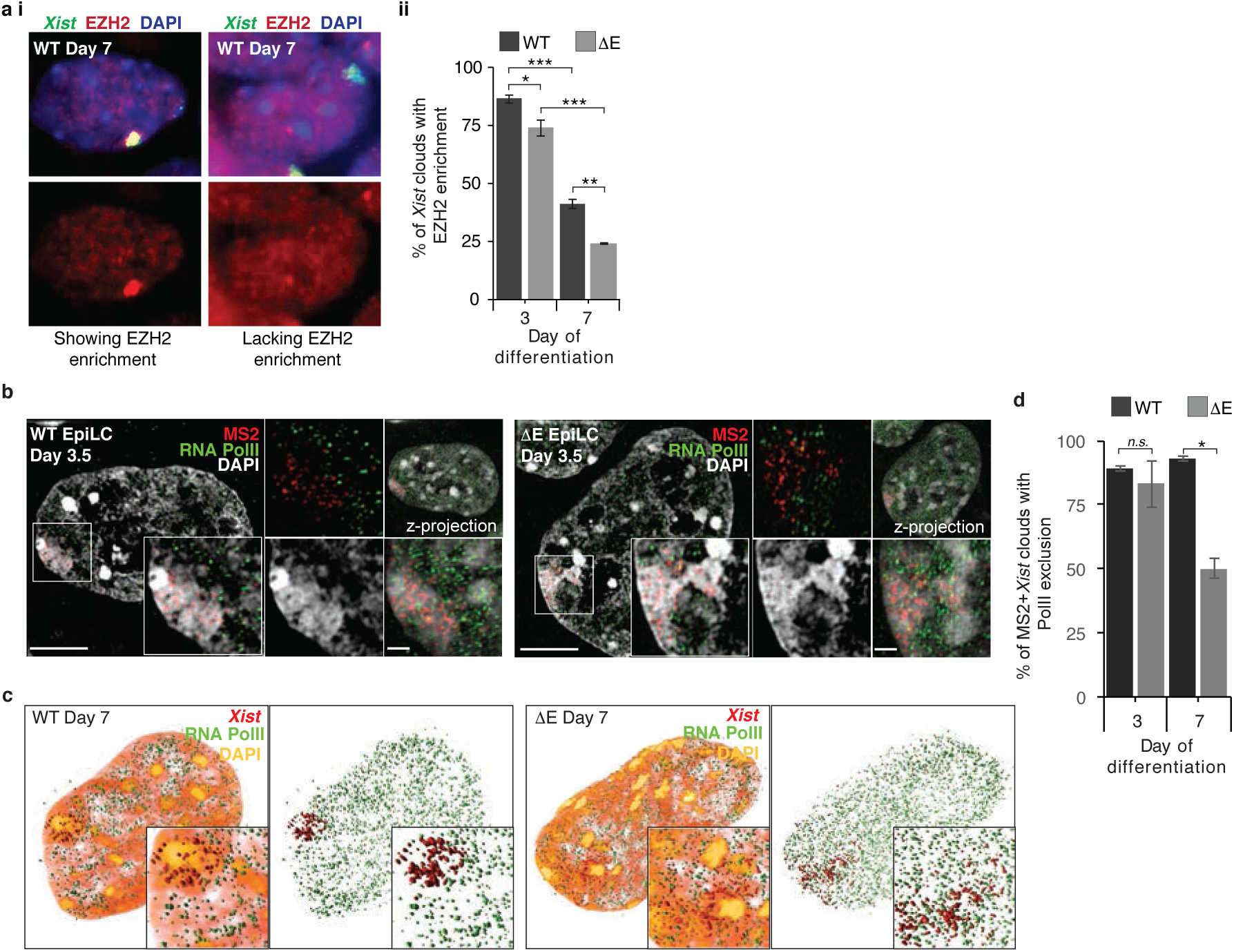
EZH2 accumulation and RNA PolII exclusion occurs on the ΔE-*Xist-*coated X-chromosome. **a i,** Epifluorescence images of WT female ESCs at day 7 of differentiation stained for EZH2 (red) and probed for *Xist* by RNA FISH (green), showing cells with (left) and without (right) EZH2 Xi-enrichment. **a ii,** Histogram showing the percentage of *Xist* clouds with a co-localized EZH2 enrichment in WT and ΔE cells at day 3 and 7 of differentiation. 60-70 *Xist* clouds were counted per sample, from 3 coverslips from 2 independent experiments. (p-value: * < 0.05, ** < 0.005, *** < 0.0005, 2-sample students t-test). Note that the E-repeat loss-of-function phenotype arises as the EZH2 enrichment on the WT X-chromosome declines. **b,** Lateral 3D-SIM optical sections through the nuclei of WT and ΔE female cells at an early stage of XCI-initiation (epiblast-like cells (EpiLCs) at day 3.5 of differentiation from ESCs) stained for RNA-pol II (green) and probed for *Xist* (red). Chromatin counterstaining with DAPI is shown in grey. Inset: RNA-pol II and *Xist* signals derived from the single optical section through the *Xist*-associated X-chromosome. Right: Same as inset except that PolII and *Xist* signals are presented independently. Far right: Z-stack projections of the cell (upper) and *Xist*-coated X-chromosome (lower). Scale bar: 5µm, Inset: 1µm. **c,** 3D Amira reconstruction of WT and ΔE cells at day 7 of differentiation (optical slices for the same cell are shown in Figure 3E) stained for RNA-polII (green), *Xist* (red), and DAPI (intensity rendering white/low to orange/high). Inset: Enlargement of the MS2-*Xist*-expressing X-chromosome. Right: Same as left with the DAPI channel removed. **d,** Histogram showing quantification of RNA-PolII exclusion from *Xist*^MS2^-coated territory in WT and ΔE cells at day 3 and 7 of differentiation. 50 cells with *Xist*^MS2^ clouds were counted from 2 coverslips from 1 differentiation experiment. Significance testing was done using a 2-tailed students t-test (p-values: * < 0.05).

**Extended Data Figure 12:**
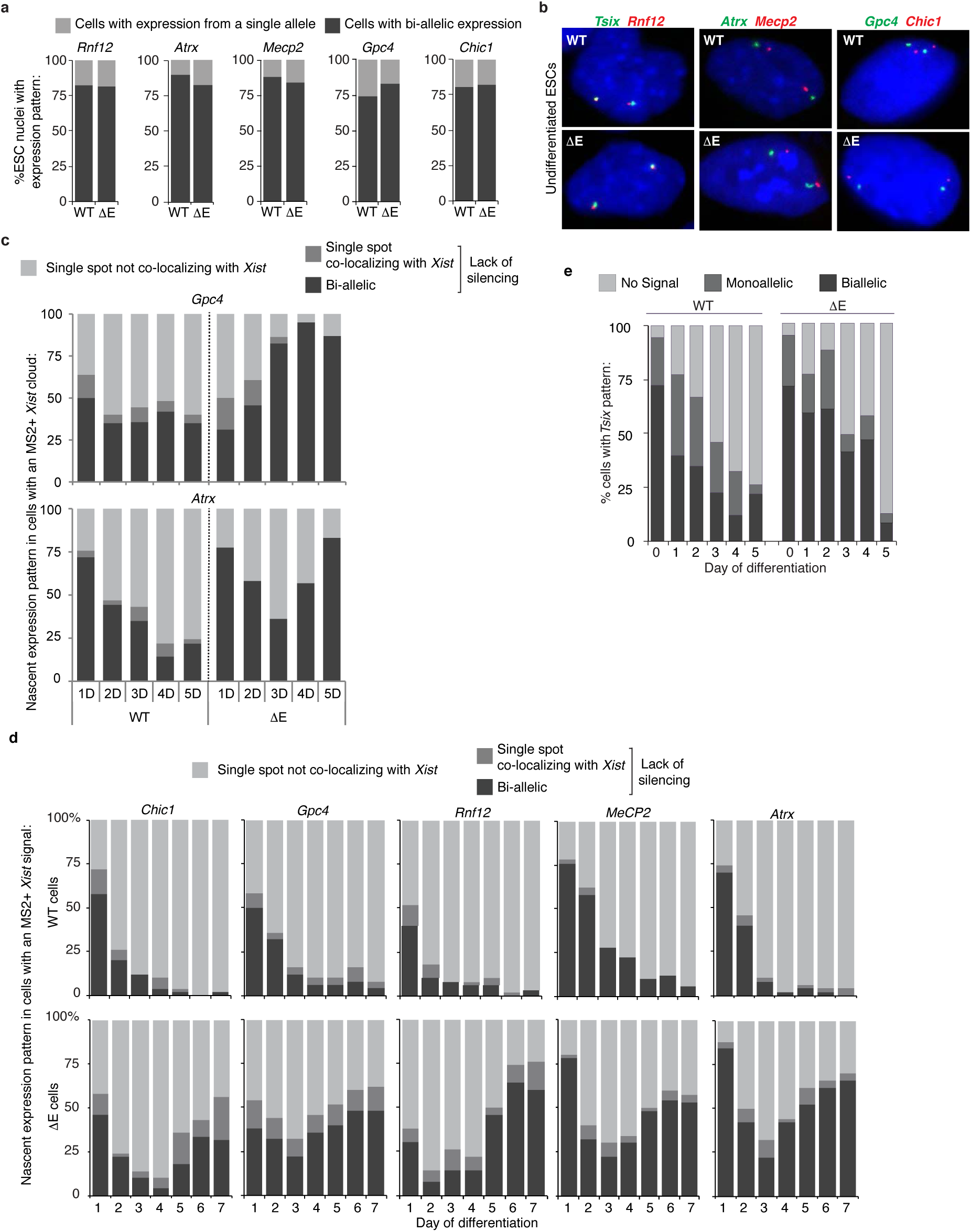
Loss of the E-repeat does not interfere with X-linked gene expression in ESCs but prevents continued gene silencing during the initiation of XCI. **a,** Histograms showing the nascent transcription pattern (mono- or bi-allelic) of indicated X-linked genes *(Rnf12 (Rlim), Atrx, Mecp2, Gpc4 and Chic1)* in undifferentiated female WT and ΔE ESCs. 64-67 cells were counted per sample from one experiment. **b,** Representative epifluorescent images of cells counted in (a). FISH signals against nascent transcripts are shown in green and red as indicated. Co-localized foci appear yellow. DAPI is shown in blue. **c,** Histograms showing the quantification of nascent expression patterns of the X-linked genes *Gpc4 and Atrx* in WT and ΔE cells displaying an MS2+*Xist^129^-*coated X-chromosome, across 5 days of differentiation. These data are derived from an independent differentiation from that shown in Figure 3c. 50 cells with an MS2+*Xist^129^* cloud were counted per time point. **d,** Histograms showing the quantification of nascent expression patterns of indicated X-linked genes in WT and ΔE cells displaying an MS2+*Xist^129^-*coated X-chromosome, across 7 days of differentiation. 50 cells with an MS2+*Xist^129^* clouds were counted per time point. These data are derived from an independent differentiation from that shown in (c) and Figure 3c. **e,** Histogram showing the quantification of nascent expression patterns of the X-linked gene *Tsix* in WT and ΔE cells across 5 days of differentiation. Note that these data were not scored relative to *Xist*^MS2^ expression (ie are derived from cells expressing either the *129* or *cas* allele of *Xist*). 77-167 cells were counted per time point, except for the ΔE cells at day 5, for which only 47 cells were counted.

**Extended Data Figure 13:**
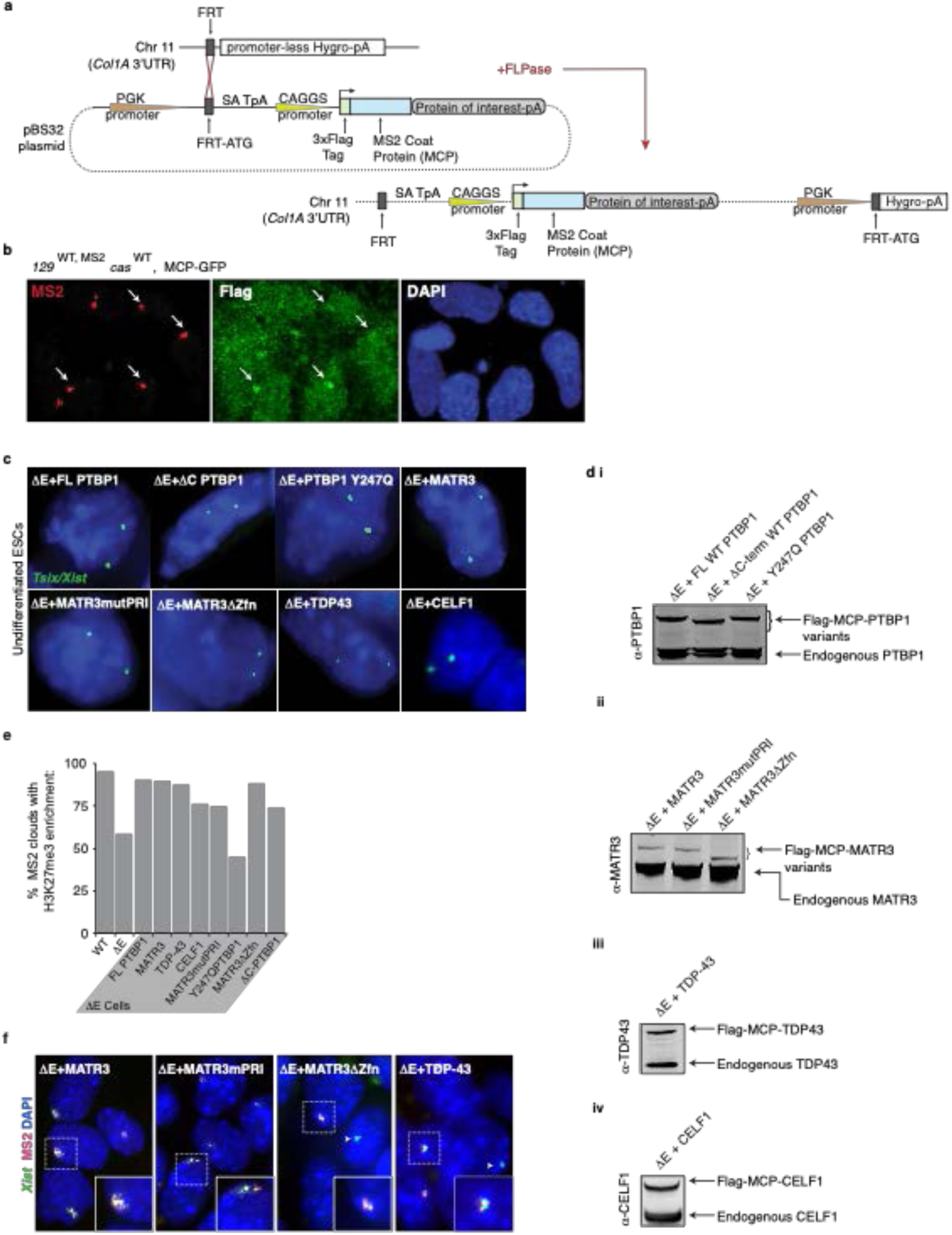
A site-specific recombination-based approach to rescue phenotypes associated with loss of the E-repeat via expression of MCP-fusions of E-repeat binding proteins. **a,** Diagram illustrating the Flp-In approach taken to constitutively express the MCP fusion proteins in WT and ΔE cells (see methods). Note that only the MCP-GFP fusion protein was expressed in WT cells. **b,** MCP-GFP fusion protein recruitment to MS2+*Xist* in WT cells at day 7 of differentiation is shown with representative epifluorescent images. Cells were CSK-extracted (see methods) prior to immunostaining for the Flag epitope (green) and RNA FISH against MS2 (red). DAPI staining is shown in blue. Arrows indicate MS2+*Xist*^1*29*^ clouds along with co-localizing Flag-MCP-GFP enrichment. **c,** RNA FISH images of *Tsix* transcripts (using a DNA probe detecting both *Xist* (which is not expressed in undifferentiated ESCs) and its antisense transcript *Tsix*) in undifferentiated ΔE ESCs expressing the indicated MCP fusion proteins as illustrated in Figure 4b. *Tsix* signal is in green, DAPI in blue. 2 *Tsix* spots were used as a proxy to confirm presence of 2 X-chromosomes in the targeted ESC lines. **d i,** Immunoblot against PTBP1 using lysates from undifferentiated ΔE ESCs expressing full-length PTBP1 or PTBP1 mutants fused to MCP. **ii,** Immunoblot against MATR3 using lysates from undifferentiated ΔE ESCs expressing full-length MATR3 and MATR3 mutants fused to MCP. **iii,** Immunoblot against TDP-43 using lysates from the undifferentiated ΔE ESCs expressing full-length TDP-43 fused to MCP. **iv,** Immunoblot against CELF1 using lysates from the undifferentiated ΔE ESCs expressing full-length CELF1 fused to MCP. **e,** Histogram showing the percentage of MS2+*Xist^129^* clouds that also show enrichment of H3K27me3 in WT or ΔE cells, or ΔE cells expressing the indicated MCP-fusion protein, differentiated for 7 days. 83-103 cells were counted per line, from one experiment. **f,** Representative epifluorescence images of RNA FISH against *Xist* (green) and MS2 (red) in day 7 differentiated ΔE cell lines expressing the indicated variants of Flag-tagged MCP fusion proteins. Inset: enlargement of MS2+*Xist^129^* cloud outlined with hatched lines. Chosen clouds are representative of the MS2+*Xist^129^* cloud phenotype in ΔE cells upon rescue with the indicated protein variant. Arrowheads indicate WT *Xist* clouds in ΔE cells, derived from the *cas* allele.

**Extended Data Figure 14:**
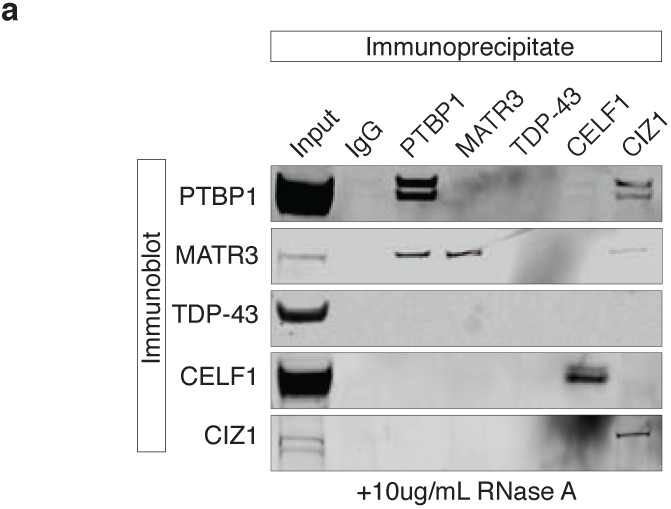
PTBP1 and MATR3 interact in the absence of RNA. Immunoprecipitation of PTBP1, MATR3, CELF1, TDP-43 and CIZ1 from ESC extracts (RNase treated) and detection of co-precipitated proteins with the same antibodies by immunoblotting (to go with Figure 4f, where no RNAse treatment was performed).

**Extended Data Figure 15:**
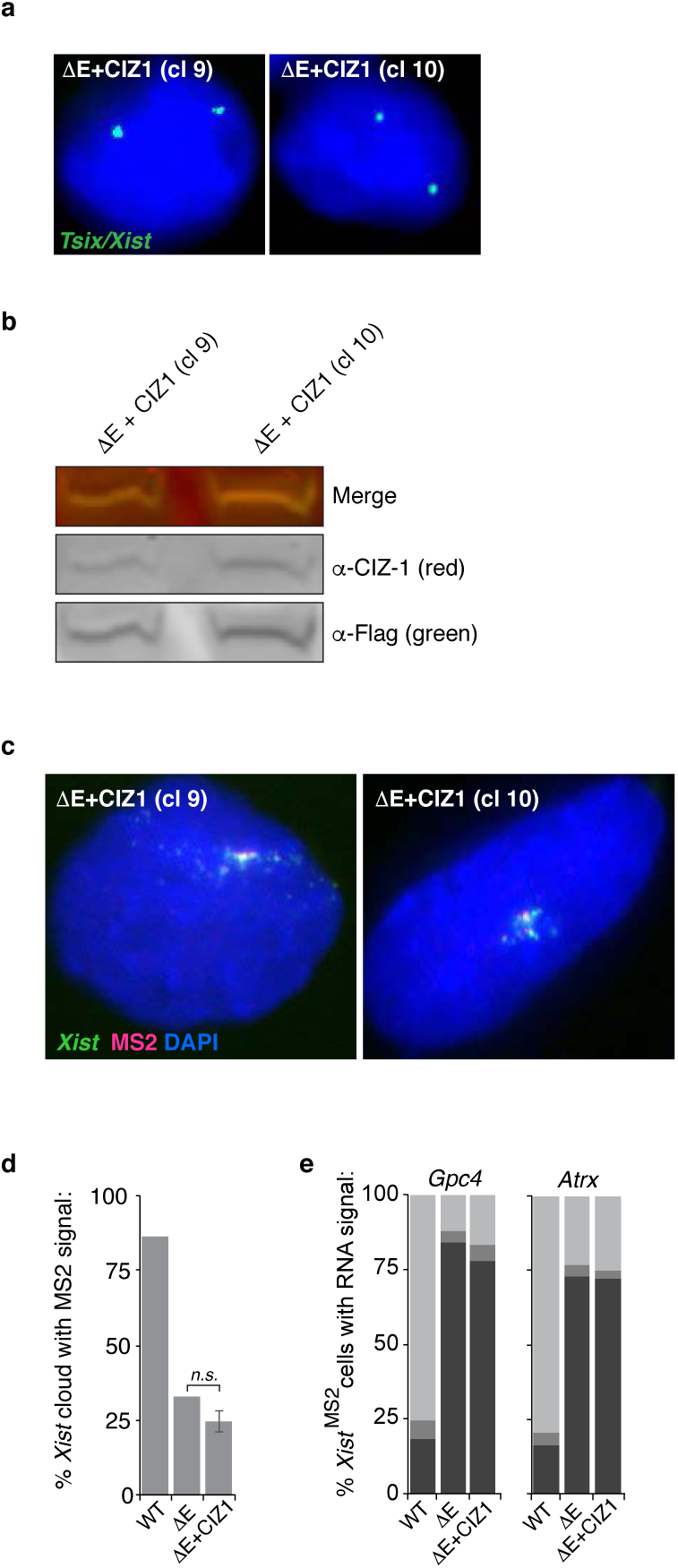
Expression of a Flag-MCP-CIZ1 fusion protein does not rescue phenotypes due to loss of the E-repeat. **a,** Representative RNA FISH images showing biallelic *Tsix* expression in MCP-CIZ1 expressing undifferentiated ΔE ESCs. *Tsix* FISH signal is in green, DAPI in blue. 2 *Tsix* spots were used as a proxy to confirm presence of 2 X-chromosomes. Two ΔE clones expressing the CIZ1 fusion protein (9 and 10) are shown. **b,** Immunoblot against CIZ1 and Flag using lysates from undifferentiated MCP-CIZ1 expressing ΔE ESCs. **c,** Representative epifluorescence images of RNA FISH s against *Xist* (green) and MS2 (red) in day 7 differentiated MCP-CIZ1 expressing ΔE cells. Two independent clones are shown. **d,** Histogram showing the number of nuclei with an *Xist* cloud that also displayed a co-localizing MS2 signal, as detected by RNA FISH, in WT, ΔE, or ΔE cells expressing Flag-tagged MCP-CIZ1. Error bar is derived from the analysis of two CIZ1 rescue clones (9 and 10) 120 – 189 cells were counted per sample from one experiment. There was no significant difference between the ΔE cells and ΔE cells MCP-CIZ1 rescue lines (2-tailed students t-test). **e,** Histograms showing the quantification of nascent *Gpc4* or *Atrx* nascent expression patterns in WT, ΔE, or ΔE cells expressing Flag-tagged MCP-CIZ1 (clone 9) displaying MS2+*Xist^129^* expression at differentiation day 7 as described in (d). 50 cells with an *Xis*t-MS2 cloud/FISH signal were counted per sample.

**Extended Data Figure 16:**
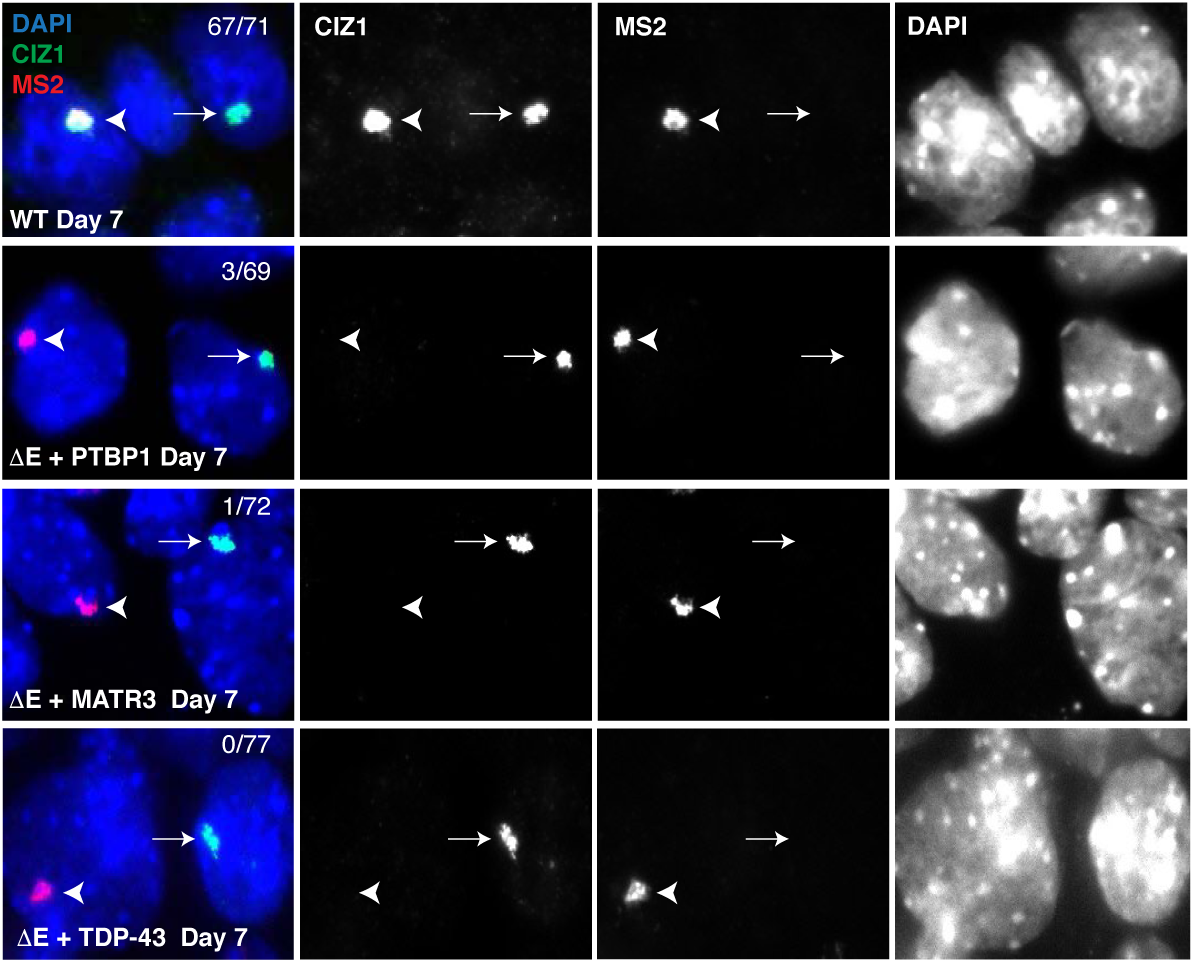
CIZ1 does not enrich on the Xi upon rescue of the E-repeat phenotypes with MCP-PTBP1, -MATR3 or -TDP-43. Representative fluorescence images showing the Xi-enrichment of CIZ1 (immunostaining, green) on the MS2+*Xist^129^* chromosome (RNA FISH, red) in WT cells at 7 days of differentiation. DAPI stained nuclei are shown in blue. The same staining for ΔE cells expressing MS2-CP-PTBP1, MS2-CP-MATR3 or MS2CP-TDP43 at day 7 of differentiation is shown below and reveals that CIZ1 does not accumulate in these cell lines. The CIZ1 Xi-enrichment seen in WT and ΔE cells expressing MCP-PTBP1, MCP-MATR3 or MCP-TDP43 that does not co-localize with the MS2+ signal derives from the *Cas* Xi that expresses a wild-type *Xist* (indicated by arrow). Arrowhead indicates rescued cloud derived from the ΔE 129 allele. Numbers at the top right in the merged image represent the percentage of MS2+ *Xist^129^* clouds showing CIZ1 co-localization. 100 cells with an *Xist*^MS2^ cloud were counted per sample, from a single differentiation experiment.

**Extended Data Figure 17:**
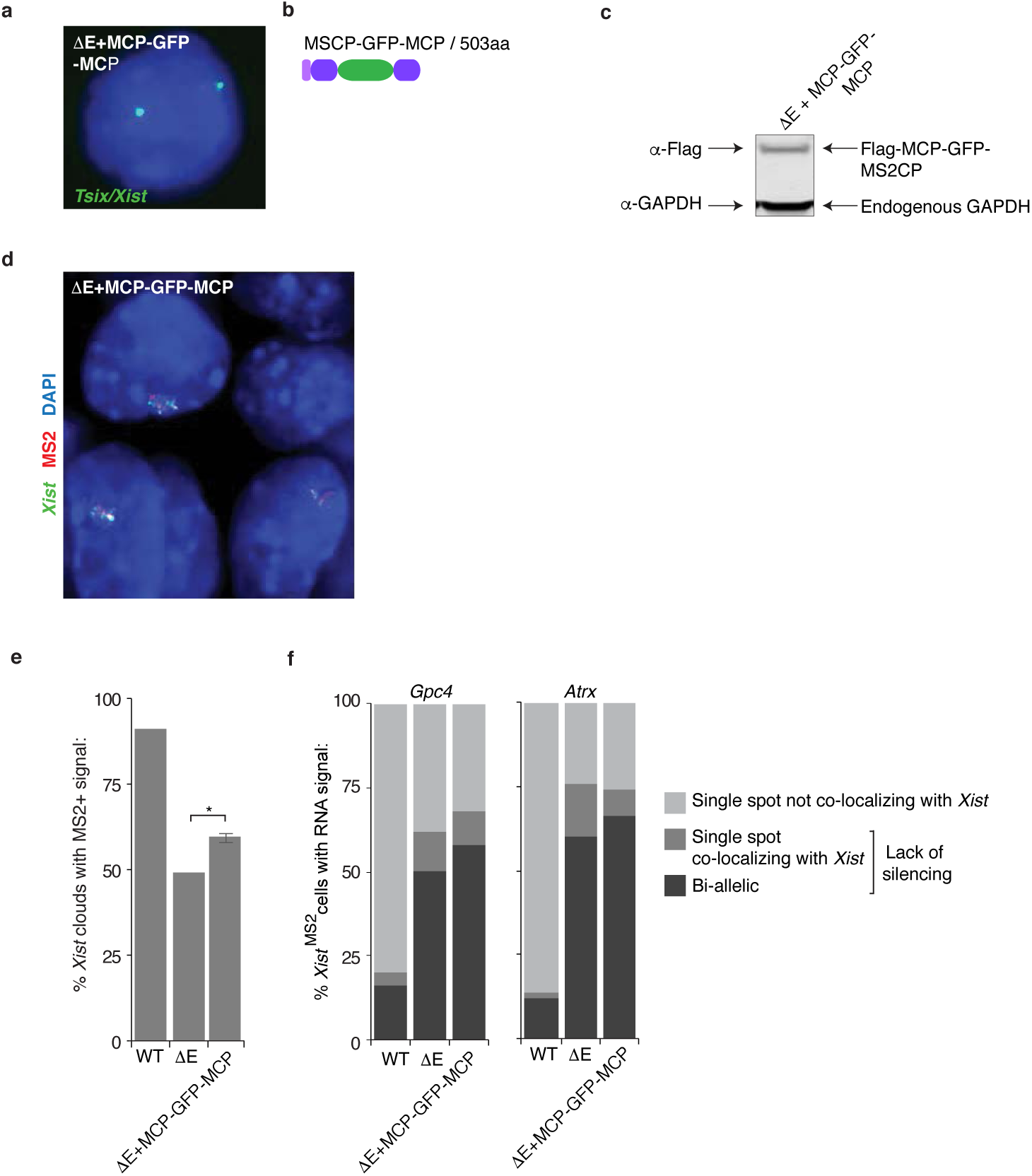
Expression of a Flag-tagged MCP-GFP-MCP fusion protein does not rescue the phenotypes associated with E-repeat loss. **a,** Representative RNA FISH images showing biallelic *Tsix* expression in MCP-GFP-MCP expressing undifferentiated ΔE ESCs. *Tsix* FISH signal is in green, DAPI in blue. 2 *Tsix* spots were used as a proxy to confirm presence of 2 X-chromosomes. **b,** Illustration of Flag-tagged MCP-GFP-MCP fusion protein. See Figure 4b for key. **c,** Immunoblot against the Flag-tag and GAPDH using lysates from undifferentiated MCP-GFP-MCP ΔE ESCs. **d,** Representative epifluorescence images of *Xist* (green) and MS2 (red) detected by RNA FISH in day 7 differentiated MCP-GFP-MCP-expressing ΔE cells. **e,** Histogram showing the percentage of *Xist* clouds with co-localizing MS2 signal at day 7 of differentiation for WT or ΔE cells and ΔE cells expressing the Flag-MCP-GFP-MCP fusion protein. 108-144 cells were counted per sample from 2 independent clones from one differentiation experiment. There was no significant difference between the ΔE cells and experimental ΔE MCP-GFP-MCP rescue lines (2-tailed students t-test). **f,** Histograms showing the quantification of nascent *Gpc4* or *Atrx* nascent expression patterns in cells displaying MS2+ *Xist* expression at differentiation day 7 in WT or ΔE cells and ΔE cells expressing the Flag-MCP-GFP-MCP fusion protein. 50 cells were counted per sample from one experiment.

**Extended Data Figure 18:**
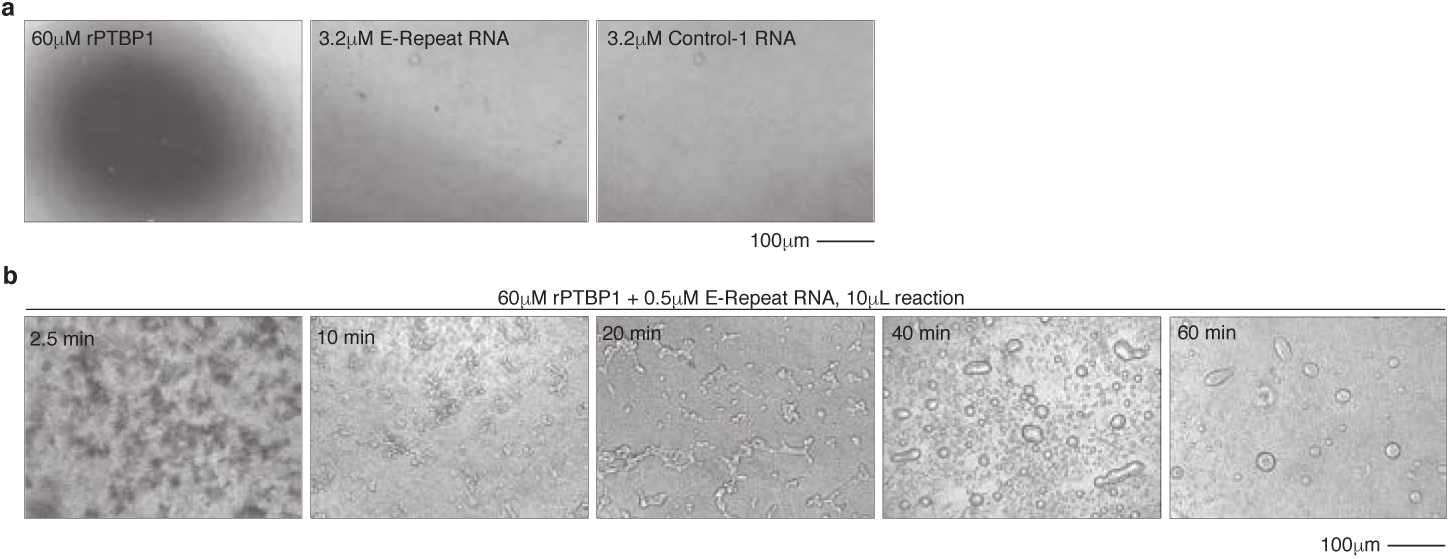
Droplet formation by PTBP1 and *Xist* E-repeat RNA occurs over time. **a,** Brightfield images showing the absence of droplet formation with 60μM rPTBP1 only, 3.2μM E-Repeat RNA only, or 3.2μM Control RNA only. **b,** Brightfield images of droplets formed with 60μM rPTBP1 and 3.2μM E-repeat RNA taken over a 1h time course.

**Extended Data Figure 19:**
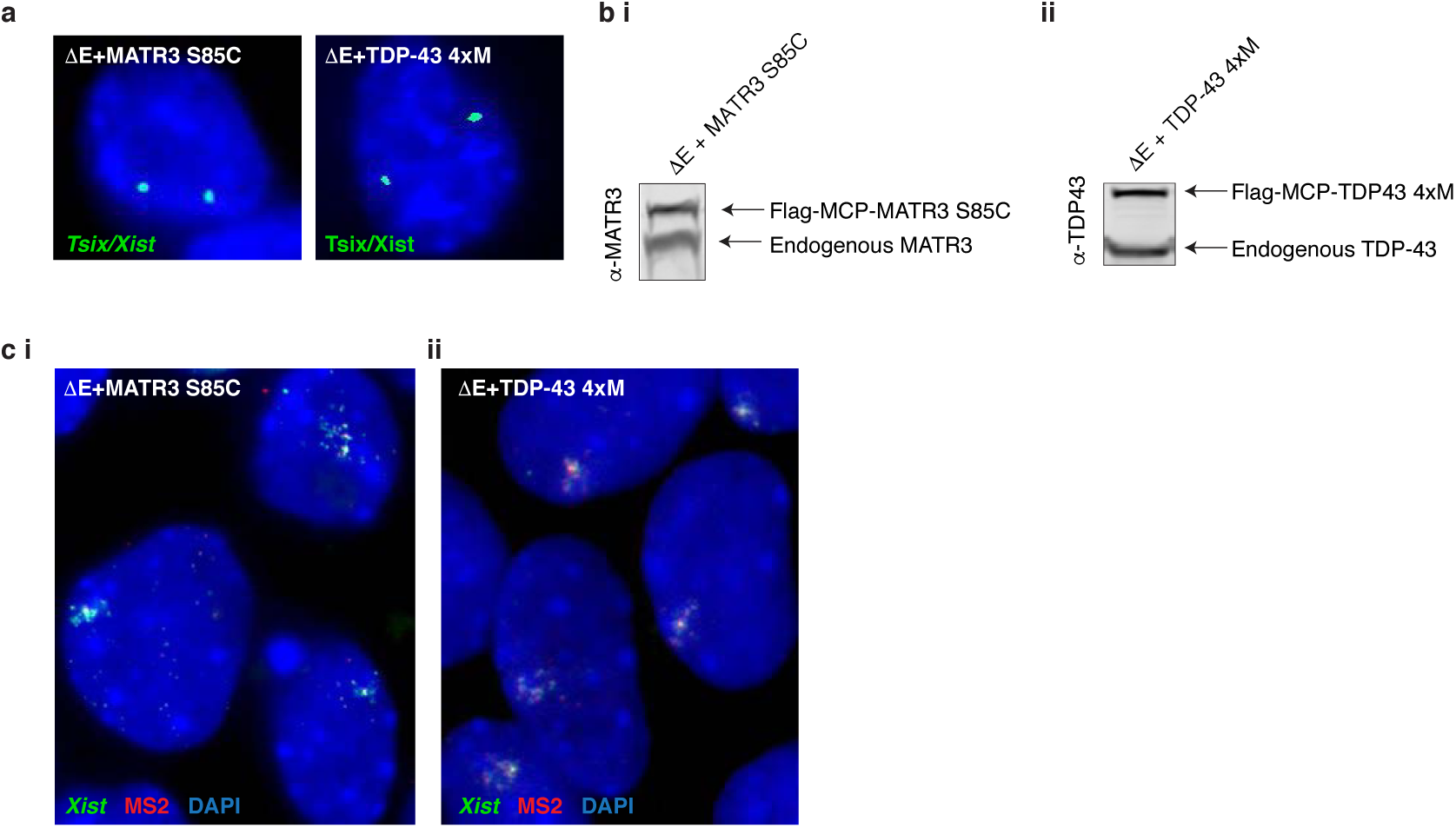
Mutations in MATR3 and TDP-43 that prevent self-association of these factors do not rescue the phenotypes observed upon loss of the E-repeat. **a,** Representative RNA FISH images showing biallelic *Tsix* expression in MCP-MATR3-S85C and MCP-TDP-43-4xM expressing undifferentiated ΔE ESCs. *Tsix* FISH signal is in green, DAPI in blue. 2 *Tsix* spots were used as a proxy to confirm presence of 2 X chromosomes. **b i,** Immunoblot against MATR3 using lysates from MCP-MATR3 S85C expressing undifferentiated ΔE ESCs. **ii,** Immunoblot against TDP-43 using lysates from MCP-TDP-43-4xM expressing undifferentiated ΔE ESCs. **c,** Representative fluorescence images of *Xist* (green) and MS2 (red) in day 7 differentiated Flag-MCP-MATR3-S85C expressing ΔE cells **(i),** and Flag-MCP-TDP-43-4xM expressing ΔE cells **(ii)**.

**Extended Data Figure 20:**
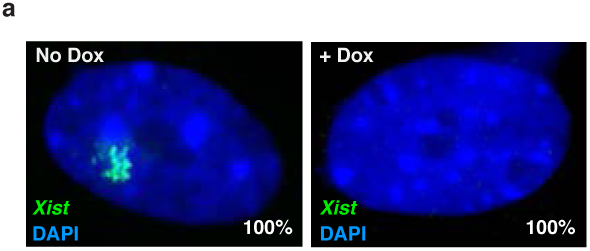
Dox treatment effectively deletes *Xist* in engineered MEFs. **a,** Images showing *Xist* as detected by RNA FISH in MEFs (*Xist*^2lox/2lox^, Rosa26^M2rtTA/tetO-Cre-recombinase^) before or after 96h of dox treatment. Numbers indicate percentage of cells with the displayed *Xist* expression pattern across 50 cells from 2 biological replicates. These data indicate that *Xist* excision completes in all cells.

**Extended Data Table 1:**
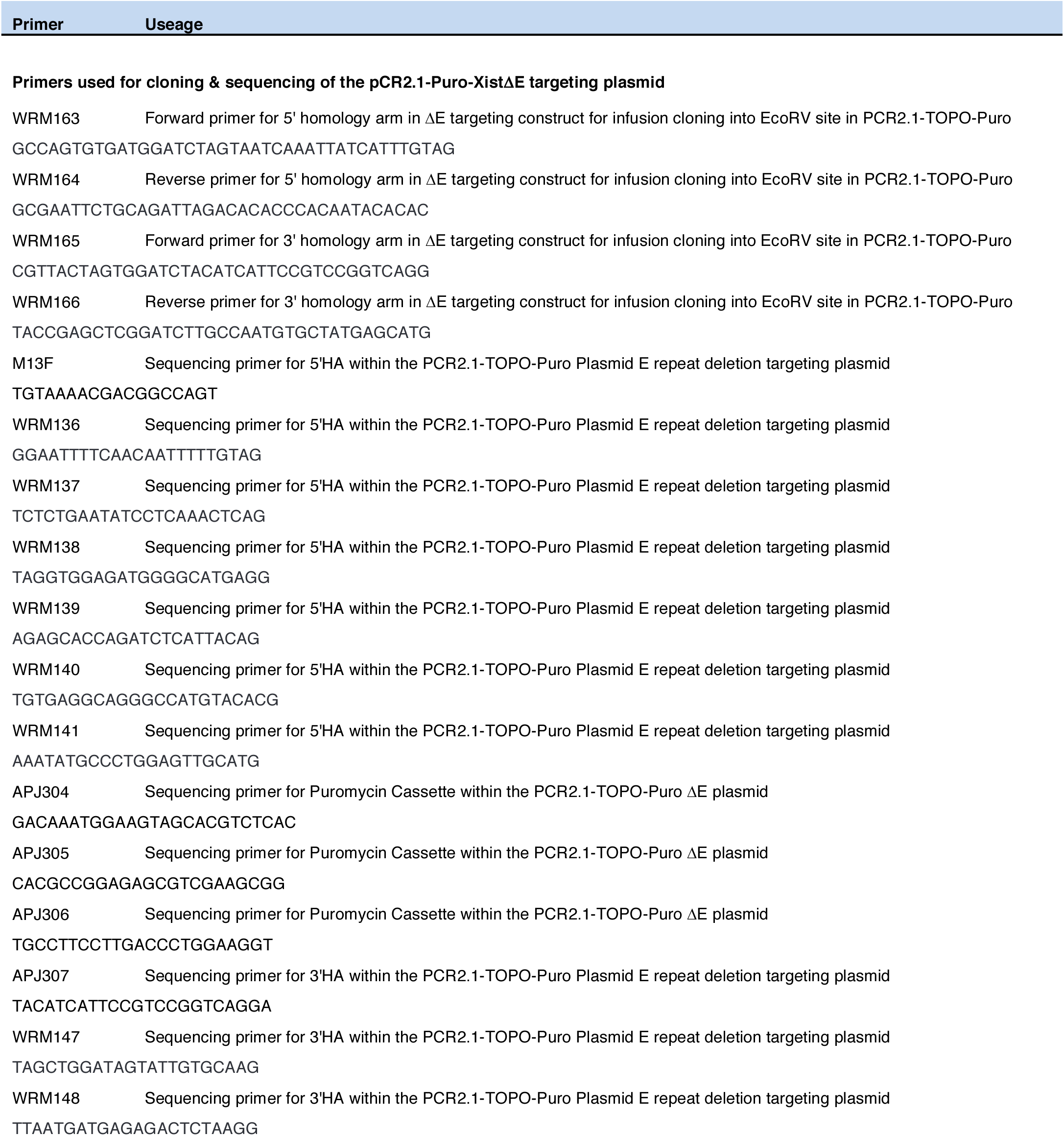
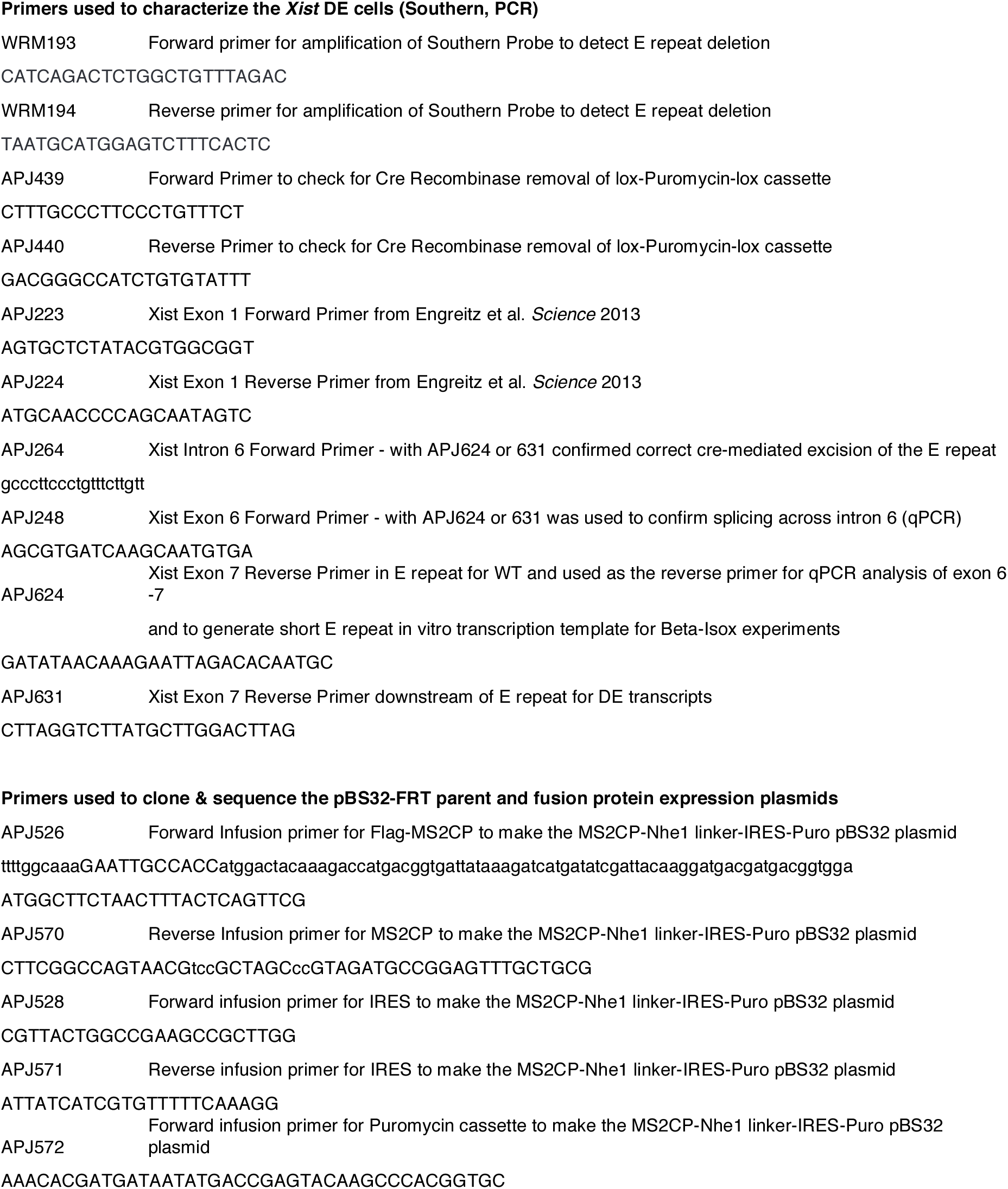
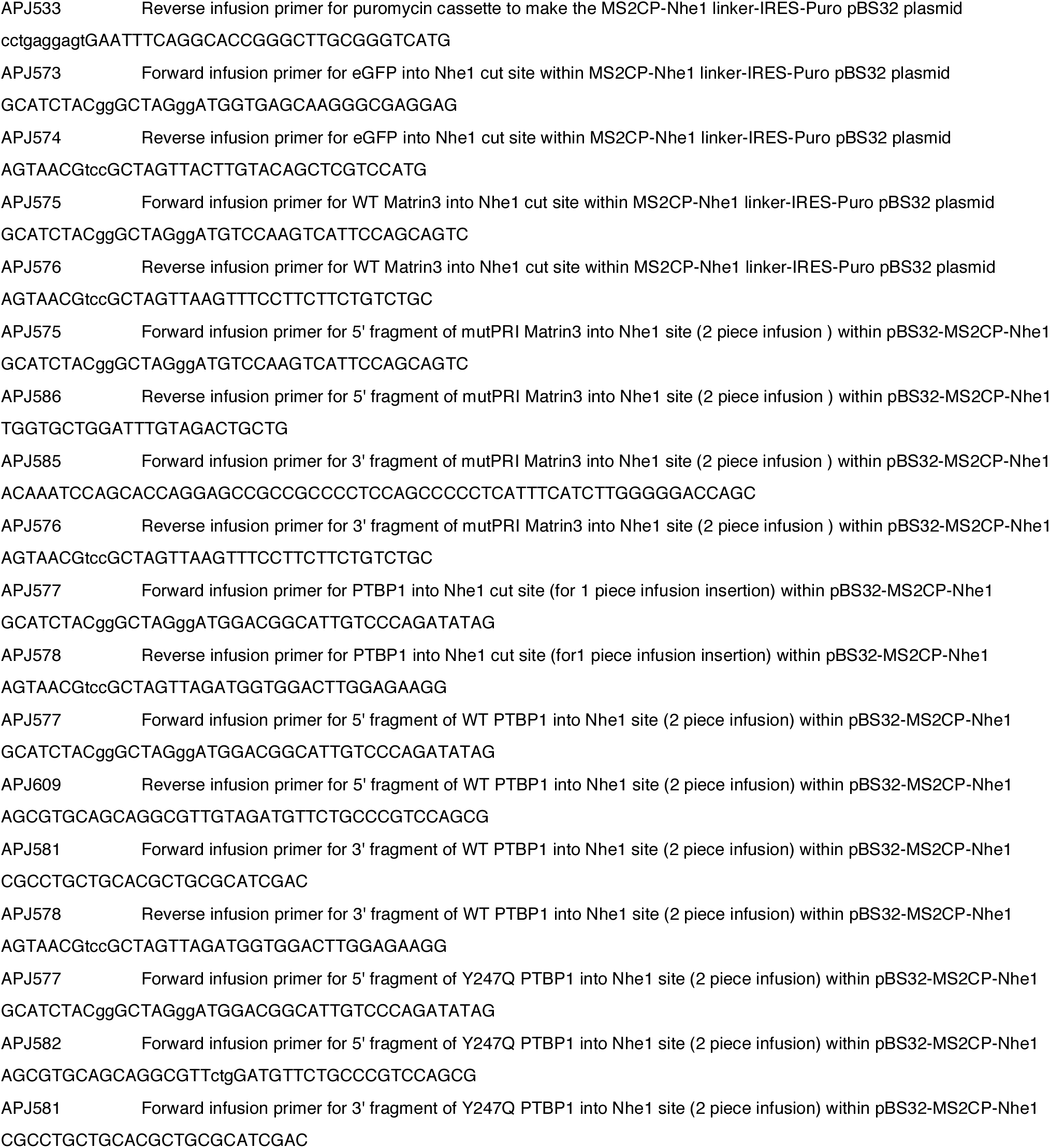
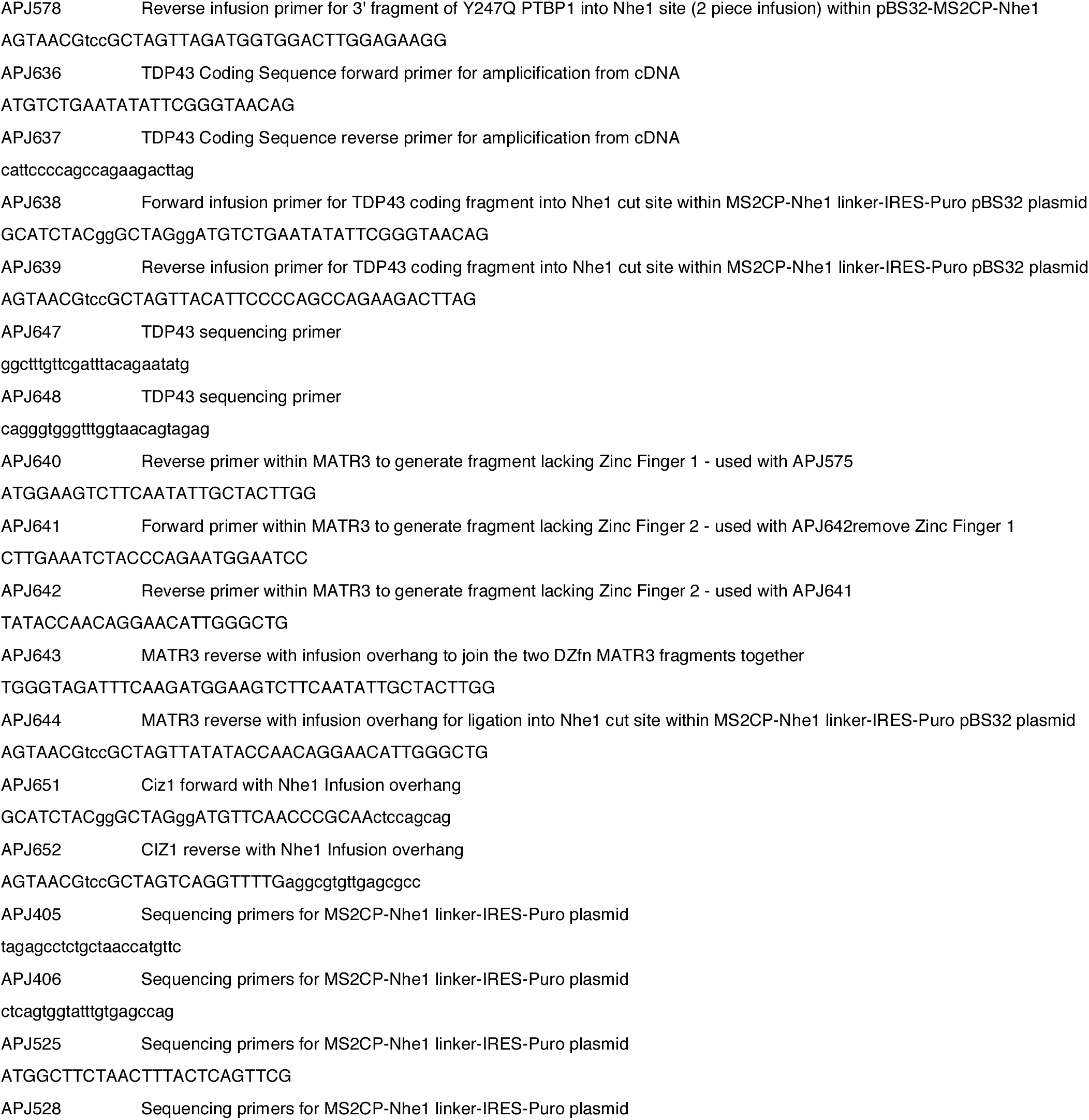
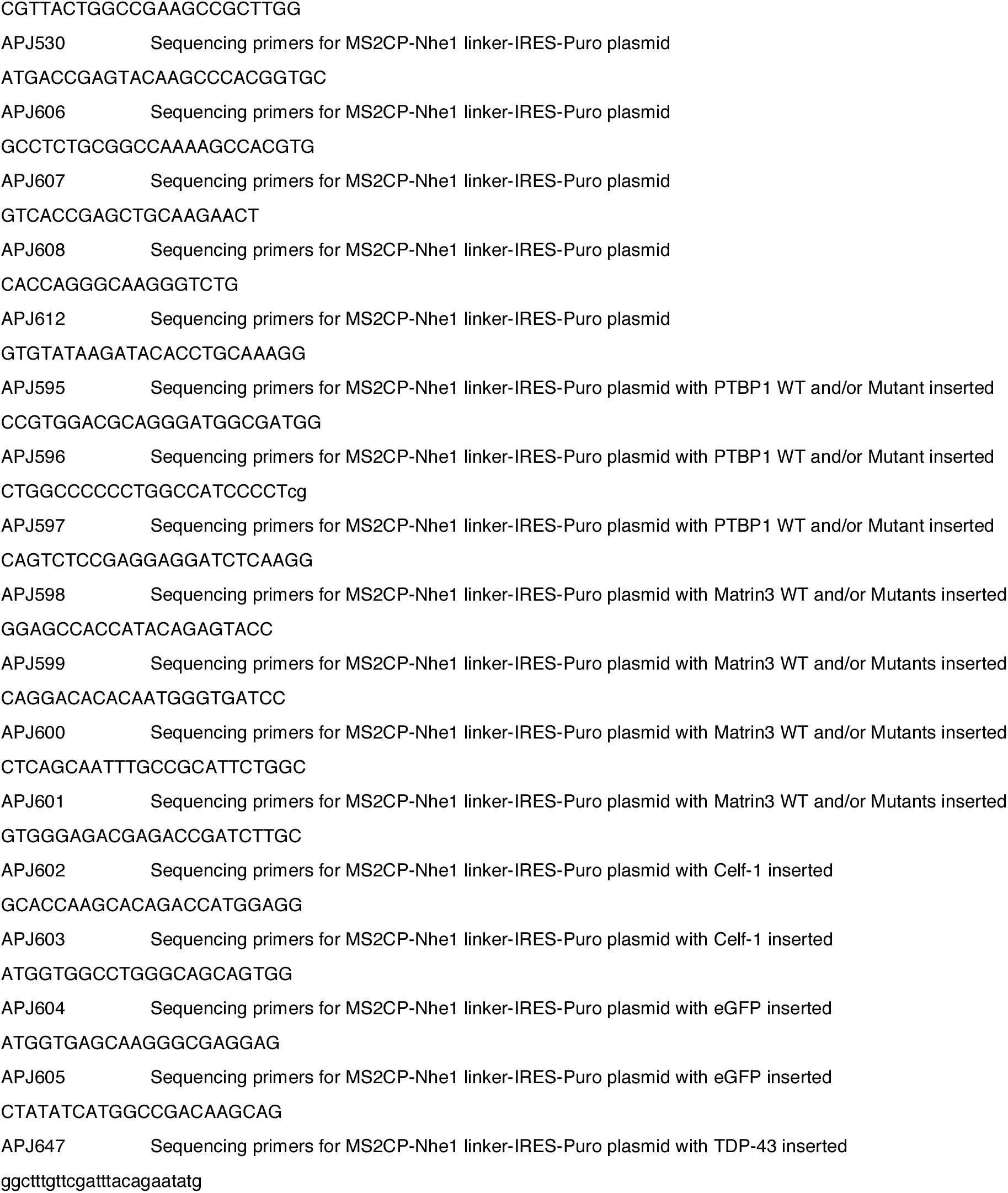
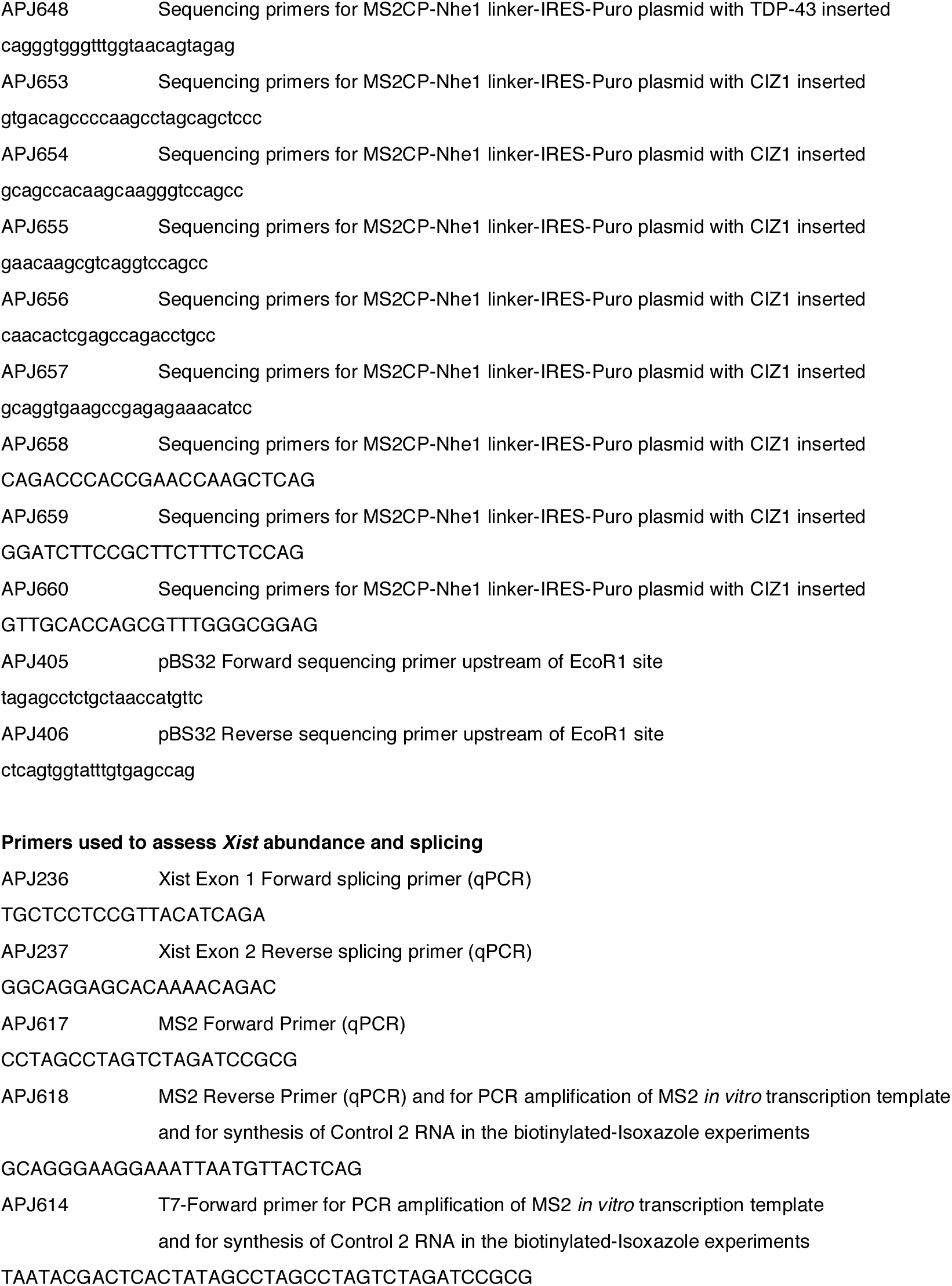
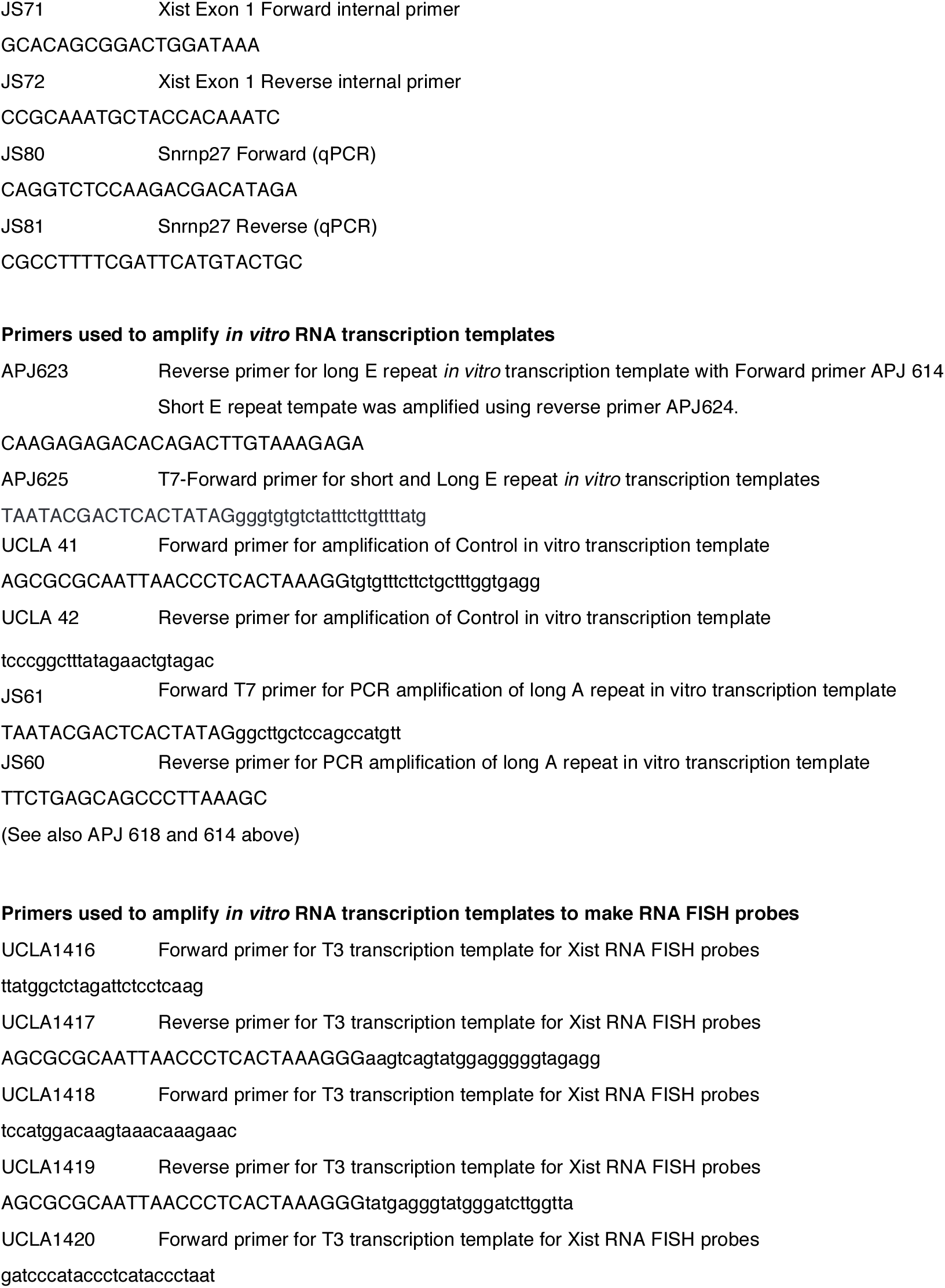
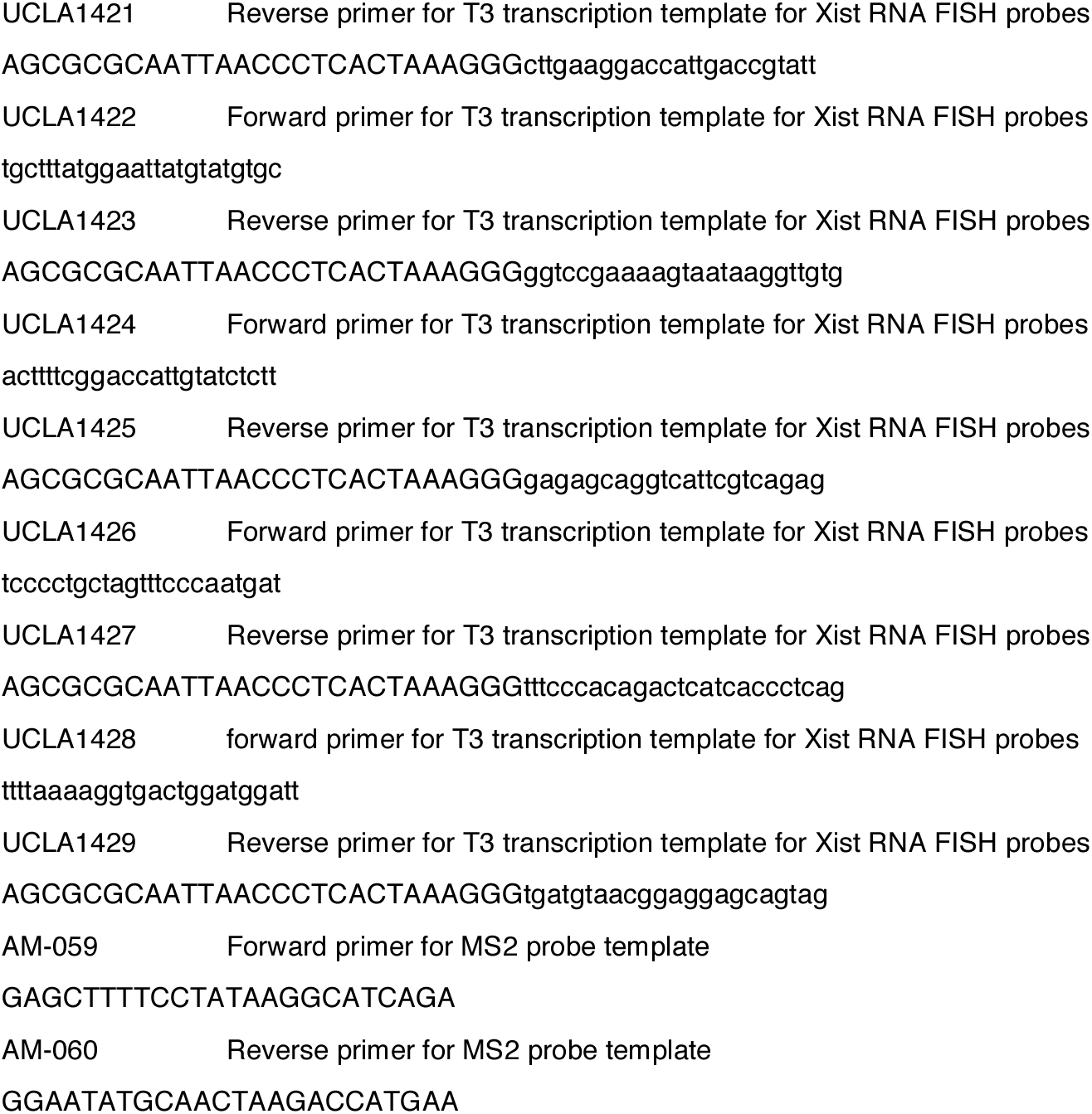
Primer information

**Extended Data Table 2:**
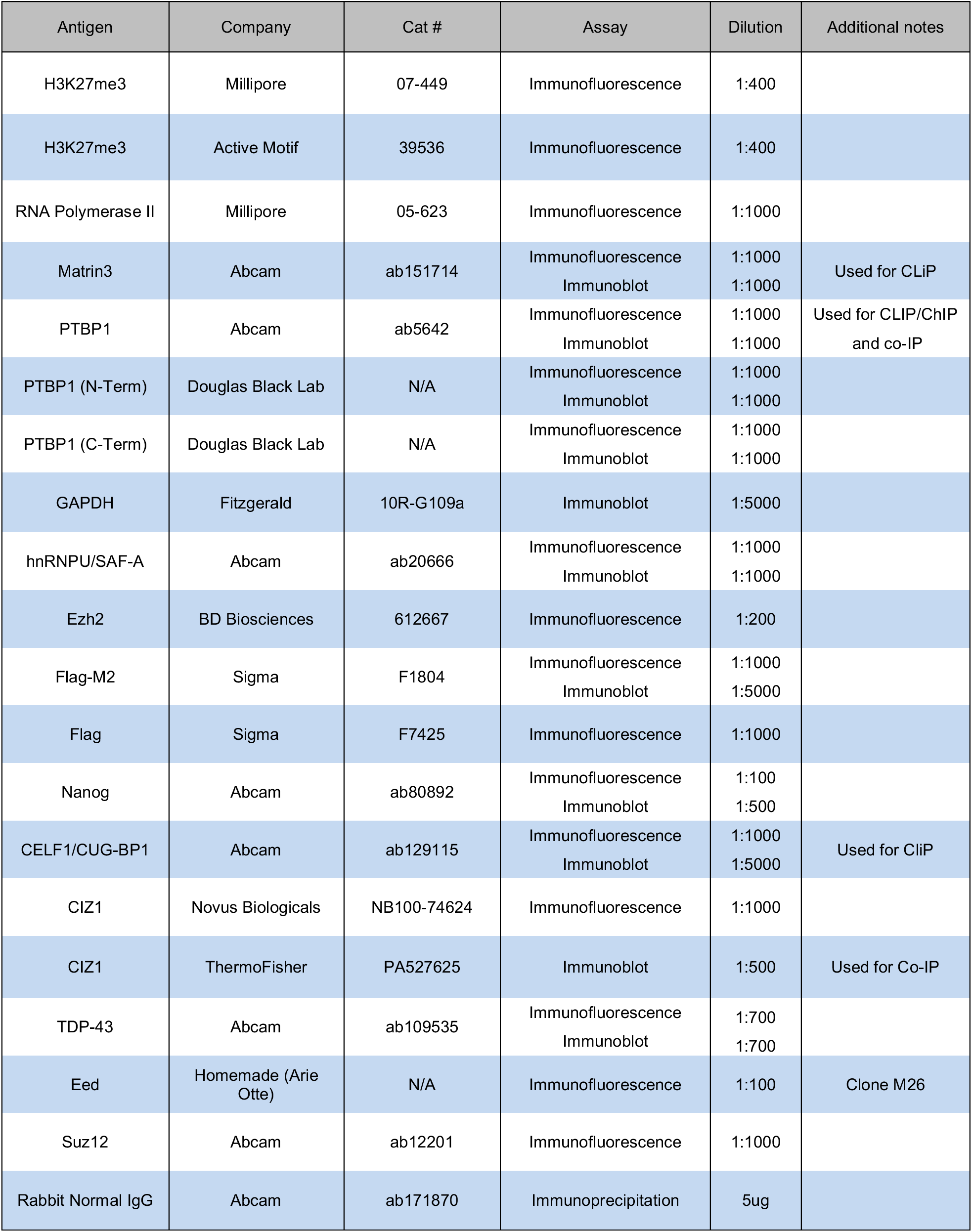
Antibody Information

